# Machine Learning-Assisted Directed Evolution Navigates a Combinatorial Epistatic Fitness Landscape with Minimal Screening Burden

**DOI:** 10.1101/2020.12.04.408955

**Authors:** Bruce J. Wittmann, Yisong Yue, Frances H. Arnold

## Abstract

Due to screening limitations, in directed evolution (DE) of proteins it is rarely feasible to fully evaluate combinatorial mutant libraries made by mutagenesis at multiple sites. Instead, DE often involves a single-step greedy optimization in which the mutation in the highest-fitness variant identified in each round of single-site mutagenesis is fixed. However, because the effects of a mutation can depend on the presence or absence of other mutations, the efficiency and effectiveness of a single-step greedy walk is influenced by both the starting variant and the order in which beneficial mutations are identified—the process is path-dependent. We recently demonstrated a path-independent machine learning-assisted approach to directed evolution (MLDE) that allows *in silico* screening of full combinatorial libraries made by simultaneous saturation mutagenesis, thus explicitly capturing the effects of cooperative mutations and bypassing the path-dependence that can limit greedy optimization. Here, we thoroughly investigate and optimize an MLDE workflow by testing a number of design considerations of the MLDE pipeline. Specifically, we (1) test the effects of different encoding strategies on MLDE efficiency, (2) integrate new models and a training procedure more amenable to protein engineering tasks, and (3) incorporate training set design strategies to avoid information-poor low-fitness protein variants (“holes”) in the training data. When applied to an epistatic, hole-filled, four-site combinatorial fitness landscape of protein G domain B1 (GB1), the resulting focused training MLDE (ftMLDE) protocol achieved the global fitness maximum up to 92% of the time at a total screening burden of 470 variants. In contrast, minimal-screening-burden single-step greedy optimization over the GB1 fitness landscape reached the global maximum just 1.2% of the time; ftMLDE matching this minimal screening burden (80 total variants) achieved the global optimum up to 9.6% of the time with a 49% higher expected maximum fitness achieved. To facilitate further development of MLDE, we present the MLDE software package (https://github.com/fhalab/MLDE), which is designed for use by protein engineers without computational or machine learning expertise.

## Introduction

Enzyme engineering has revolutionized multiple industries by making chemical processes cheaper, greener, less wasteful, and overall more efficient. Enzymes and other proteins are engineered by searching the fitness landscape, a surface in a high-dimensional space that relates a desired function (“fitness”) to amino acid sequences.^1,2^ Exploring this landscape is extremely challenging: the search space grows exponentially with the number of amino acid positions considered, functional proteins are extremely rare, and experimental screening of proteins can be resource-intensive, with researchers often limited to testing a few hundred or thousand variants. Directed evolution (DE) can overcome these challenges by employing a greedy local search to optimize protein fitness.^3^ In its lowest-screening-burden form (hereafter referred to as “traditional DE”), DE starts from a protein having some level of the desired function, then iterates through rounds of mutation and screening, where in each round single mutations are made (e.g. by site saturation mutagenesis) to create a library of variants and the best variant is identified and fixed; iteration continues until a suitable level of improvement is achieved (Figure 1A).

**Figure 1.**
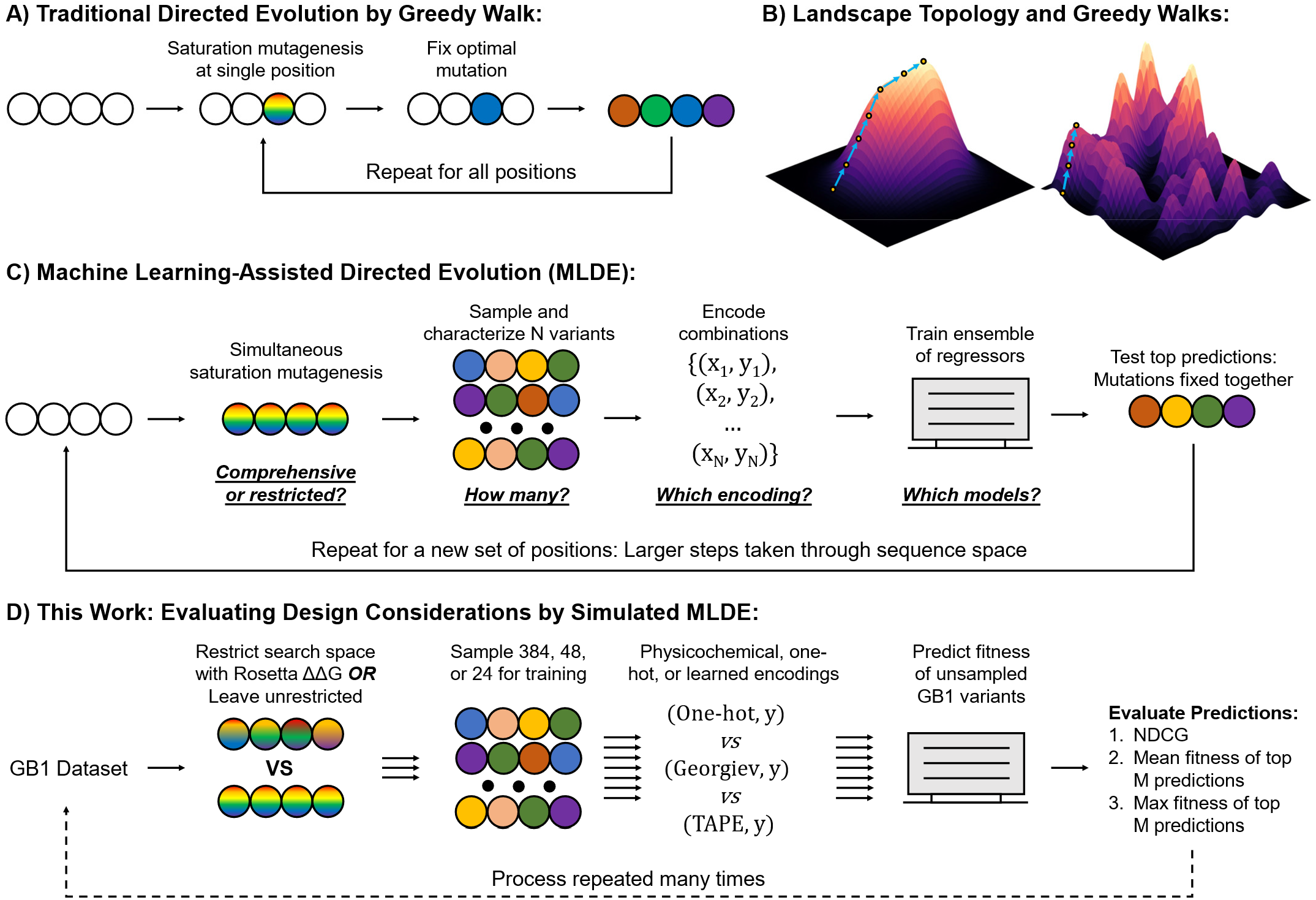
(A) Directed evolution (DE) by single-mutation greedy walk (“Traditional DE”). In this approach, mutations are fixed iteratively by walking up the steepest fitness gradient. (B) Smooth (left) vs rugged (right) fitness landscapes. A smooth fitness landscape contains a single fitness maximum, so traditional DE is guaranteed to eventually reach the global optimum, though the number of steps needed will depend on the topology of the peak. A rugged fitness landscape contains multiple fitness maxima. Traditional DE is only guaranteed to reach a local fitness optimum here; the maximum achieved will depend on the starting protein variant and the order in which positions are chosen for mutagenesis and testing. (C) Machine learning-assisted directed evolution (MLDE). In this approach, a sample drawn from a multi-site simultaneous saturation mutagenesis (“combinatorial”) library at a set of positions is used to train an ensemble of regressors. This ensemble is used to predict the best combinations of mutations not seen in the initial draw, which are then tested experimentally. Because the best mutations are fixed simultaneously, MLDE operates in a path-independent manner, so the global optimum of a combinatorial space can be achieved regardless of the starting point. Once mutations are fixed for a given set of positions, a new set is chosen and the procedure is repeated, allowing for larger, more efficient steps through sequence space. The MLDE procedure has many design considerations, which are highlighted as questions under each step. (D) The simulation procedure used throughout this study to evaluate improvements to the MLDE workflow, with tests performed to evaluate the different design considerations given above each step. The simulation procedure is repeated many times using data from the GB1 landscape. The effectiveness of a round of simulated MLDE is determined by (1) calculating the normalized discounted cumulative gain (NCDG) over all predictions in the simulation and (2) evaluating the mean and max true fitness of the M variants with highest predicted fitness.

By focusing on single mutations rather than combinations of mutations, traditional DE can be used to optimize protein fitness with a low screening burden. The process is highly effective when the beneficial effects of mutations made at different sequence positions are additive; however, focusing on single mutants ignores the codependence of mutations (epistasis).^4,5^ Epistasis is commonly observed, for example, between residues close together in an enzyme active site or protein binding pocket, where mutations often affect function. Epistatic effects can decrease the efficiency of DE by altering the shape of the protein fitness landscape. Specifically, epistasis can alter gradients on the fitness landscape to make the route to a global optimum very long,^6^ or it can introduce local optima at which traditional DE can become trapped (Figure 1B). Both lower the average fitness that can be achieved for a given screening burden. The only way to account for epistasis during optimization is to evaluate and fix combinations of mutations, bypassing the path-dependence of traditional DE. Due to limited screening capacity, however, this is intractable for most protein engineering projects.

Increasingly, machine learning (ML) is being used to ease experimental screening burden by evaluating proteins *in silico*.^7–10^ Data-driven ML models learn a function that approximates the protein fitness landscape, and they require little to no physical, chemical, or biological knowledge of the problem. Once trained, these models are used to predict the fitness of previously unseen protein variants, dramatically increasing screening capacity and expanding the scope of the protein fitness landscape that can be explored. We recently demonstrated a machine learning-assisted directed evolution (MLDE) strategy for navigation of epistatic fitness landscapes that cover a small number of amino acid sites.^11^ MLDE works by training an ML model on a small sample (10^1^–10^2^) of variants from a multi-site simultaneous saturation mutagenesis (“combinatorial”) library, each with an experimentally determined fitness; the model is then used to predict the fitness of all remaining variants in the combinatorial library (10^4^–10^5^), effectively exploring the full combinatorial space. Combinations with the highest predicted fitness are experimentally evaluated, the best combination is fixed, and another round of MLDE is started at a new set of positions (Figure 1C). The iterative nature of MLDE is identical to that of traditional DE, but by evaluating and fixing multiple cooperative mutations, MLDE avoids some local fitness traps or long paths to the global optimum for each combinatorial library.

Our original MLDE work serves as a baseline, as it did not explore the many design considerations of MLDE (Figure 1C, bold and underlined questions).^11^ Two notable considerations are (1) the choice of encoding strategy and (2) the handling of low-fitness variants in combinatorial libraries. Protein sequences must be numerically encoded to be used in ML algorithms, and the choice of encoding will affect the outcome of learning. In our original implementation, we used a one-hot encoding scheme that captures no information about the biochemical nature of different amino acids. Mutating an amino acid to a similar one (in terms of size, charge, etc.) is less likely to significantly affect protein fitness than mutating it to a very different one, however, and this knowledge can be transferred into ML models via the encoding strategy. The effectiveness of an ML model is also determined by the information content of the data used to train it, and so the choice of variants to use for the training stage of MLDE is important. Combinatorial libraries tend to be enriched in zero- or extremely low-fitness variants, particularly in regions critical to protein function like an enzyme active site.^12–14^ These “holes” provide minimal information about the topology of the regions of interest in a fitness landscape (i.e. they provide no information about regions with functioning proteins and no information about the *extent* to which different mutations affect fitness) and can bias ML models to be more effective at predicting low-fitness variants than high-fitness ones, the opposite of our goal. In our original implementation, we opted to sample randomly from full combinatorial spaces to generate training data with high sequence diversity. Because combinatorial landscapes tend to be dominated by holes, however, this random draw primarily returned sequences with extremely low or zero fitness, resulting in training data that, despite containing diverse sequences, was information-poor.

In this work, we evaluate various design considerations by simulating MLDE on the empirically determined four-site combinatorial fitness landscape (total theoretical size of 20^4^ = 160,000 protein variants) of protein G domain B1 (GB1) (Figure 1D).^15^ Containing multiple fitness peaks (the routes to which are not always direct) and heavily populated by zero- and low-fitness variants (92% have fitness below 1% of that of the global maximum), this landscape not only presents an ideal testing ground in which to compare the abilities of traditional DE and MLDE to navigate epistatic fitness landscapes, but also serves to test the ability of ML methods to navigate hole-filled regions of protein fitness landscapes. We begin by evaluating a number of alternate encoding strategies to one-hot, including physicochemical encodings and embeddings derived from five different natural language processing models.^16,17^ Next, we demonstrate how integration of models and training procedures better tailored for protein fitness landscapes (in particular 1D convolutional neural networks and gradient boosted Tweedie regression) into the workflow can improve MLDE performance.^18–20^ We then show the importance of reducing uninformative holes in MLDE training sets and propose integrating some form of zero-shot prediction (i.e. prediction of variant fitness prior to data collection) into the MLDE pipeline to generate more informative training data. We call the general strategy of running MLDE with training sets designed to avoid holes “focused training MLDE” (ftMLDE). As a demonstration, we show how predicted ΔΔG of protein stability upon mutation can be used as an effective zero-shot predictor of GB1 fitness, and then use this predictor to generate information-rich training data. Using this training data, we finally test the effect of training set size on the outcome of ftMLDE, finding that our improved procedure achieved the GB1 global maximum up to 77-fold more frequently than traditional DE.

To summarize our contributions, this paper describes significant improvements to our original method. It also highlights (1) the importance of considering the unique attributes of fitness landscapes when applying ML to protein engineering problems and (2) how the three major paradigms of protein engineering (directed evolution, rational design, and, increasingly, machine learning) can be combined into a cohesive, highly efficient engineering pipeline. To improve access to such a pipeline, we introduce the MLDE software package, made available on the Arnold Lab GitHub (https://github.com/fhalab/MLDE). Designed to be accessible to non-ML and non-computational experts, this repository contains Python scripts that allow execution of MLDE on arbitrary combinatorial fitness landscapes, thus enabling wet-lab application.

## Results and Discussion

### MLDE Procedure and Simulated MLDE

MLDE attempts to learn a function that maps protein sequence to protein fitness for a multi-site simultaneous saturation mutagenesis (“combinatorial”) library. The procedure begins with measuring the fitness values of a small subsample from the library. These labeled variants are then used to train an ensemble of models with varied architectures (models come from scikit-learn,^21^ Keras, and XGBoost^18^ (Supporting Information: *Inbuilt Models*)), employing k-fold cross validation to measure a validation error for each model class. Predictions from the top-performing trained models (as measured by validation error) are averaged to predict fitness values for the unsampled (“unlabeled”) variants. Unlabeled variants are ranked according to predicted fitness, and the top M are evaluated experimentally to identify the best-performing ones (Figure 1C). More detailed information about the implementation of MLDE can be found in the supporting information (*MLDE Programmatic Implementation*).

Throughout this work, we demonstrate improvements to MLDE through simulation on the empirically determined four-site combinatorial fitness landscape of protein G domain B1 (GB1).^15^ Originally reported by Wu *et al.*, this landscape consists of 149,361 experimentally determined fitness measurements for 160,000 possible variants, where fitness is defined by both the ability of the protein to fold and the ability of the protein to bind antibody IgG-Fc. To our knowledge, this landscape is the only published one of its kind (i.e. the only almost-complete combinatorial landscape where fitness is reported as scalar values amenable to training the regression models used in MLDE). By imputing the fitness of the remaining 10,639 variants and evaluating the resultant complete landscape, Wu *et al.* identified 30 local optima, the routes to which were often indirect (e.g. if a local optimum was four mutations away from a starting point, it would take more than four mutations to travel by single-mutation greedy walk from the starting point to the optimum). Epistatic interactions are thus highly prevalent in the GB1 landscape. The goal of simulated MLDE is to mimic what we would expect had we performed thousands of MLDE experiments on GB1. Thus, to ensure that our simulations match what would have been observed experimentally had our simulated experiments actually been performed, we do not use the variants with imputed fitness in this study.

A round of simulated MLDE begins with generating labeled variants (Figure 1D). Here, we draw a small set of variants from the GB1 landscape and attach their known fitness values—this stage of the simulation is analogous to building a combinatorial library, picking colonies from an agar plate, then expressing and assaying the variants harbored by the colonies. The labeled variants are then fed into the MLDE pipeline and the average predictions of the top three models (as measured by cross-validation error) are used to rank the unlabeled variants by predicted fitness. The quality of the returned ordering is evaluated using a combination of metrics, including (1) the ranking metric “normalized discounted cumulative gain” (NDCG) calculated over the full ordering of unlabeled data and using the true fitness as the gain, (2) the mean fitness of the M-highest-ranked unlabeled variants, and (3) the max fitness of the M-highest-ranked unlabeled variants (Supporting Information: *Evaluation Metrics*). The NDCG score evaluates the predictive performance of a model over all unlabeled datapoints, while the mean and max fitness of the M-highest-ranked variants evaluate how well our process predicts the highest-fitness variants in the fitness landscape. Evaluating the M-highest variants is analogous to experimentally constructing and characterizing the M protein variants predicted to have highest fitness. Whenever reported, mean and max fitness are normalized to the highest fitness in the unlabeled dataset and so can typically be interpreted as a fraction of the global maximum in the GB1 dataset.

### More Informative Encodings Improve MLDE Outcome

Protein sequences must be numerically encoded to be used in ML algorithms. Our previous implementation of MLDE used one-hot encoding, an uninformative categorical encoding strategy that captures no information about amino acid similarities and differences. To investigate the effects of more informative encodings, we tested physicochemical parameters as well as learned embeddings. Physicochemical parameters are manually curated indices that describe amino acid qualities such as hydrophobicity, volume, mutability, etc. In this work, we used the set of physicochemical parameters developed by Georgiev, which is a low-dimensional representation of over 500 amino acid indices from the AAIndex database.^16,22,23^ Learned embeddings are vectors of continuous values extracted from machine learning models trained on unlabeled data, and they capture similarities and differences between specific amino acids as well as contextual information about amino acid positions in a protein. Here, we evaluate the effectiveness of the learned protein embeddings generated by all fully unsupervised models made available by Rao *et al.*^17^ These include embeddings generated from models with a transformer architecture (“Transformer”),^24^ three separate LSTM-based architectures (“LSTM”, “UniRep”, and “Bepler”),^25–27^ and a dilated residual network architecture (“ResNet”),^28^ all trained on ~30 million protein sequences from the Pfam database.^29^

To compare the different encoding strategies, we performed 2000 rounds of simulated MLDE, using 384 randomly drawn GB1 variants as training data in each simulation (Methods: *Encoding Comparison Simulations*). For the sake of computational efficiency, only XGBoost and Keras models were used in the ensemble of MLDE models for the largest embeddings (LSTM and UniRep). Importantly, the training data and cross-validation indices were kept the same for each encoding strategy in each simulation, allowing for pairwise comparison of simulation results. As shown in Figure 2, more informative encodings generally achieved a higher NDCG than one-hot. These results become more apparent when making pairwise comparisons (Supporting Figure S6), with LSTM-derived encodings achieving a higher NDCG than one-hot in 97.4% of simulations, transformer-derived in 92.1%, UniRep-derived in 89.6%, Bepler-derived in 79%, ResNet-derived in 67.15% and Georgiev in 90.4%. For all encodings except UniRep-derived, those that yielded a higher NDCG than one-hot also had a higher expected mean fitness in the top 96 predictions (Supporting Figures S7–S12 for pairwise comparisons). This pattern did not hold for the expected max fitness in the top 96 predictions, with only Georgiev encodings yielding a higher expected max fitness than one-hot.

**Figure 2.**
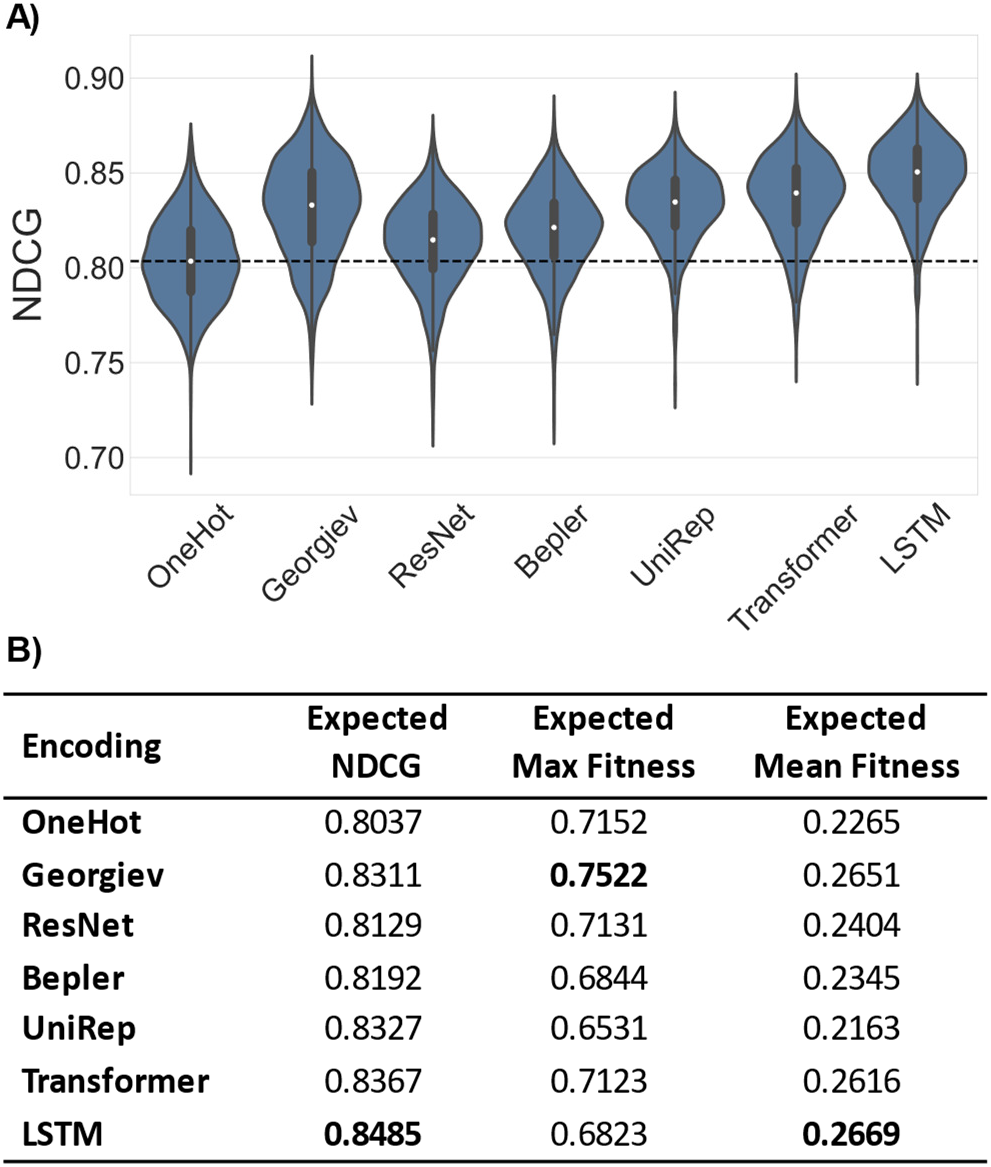
Results of simulated MLDE comparing seven different encoding strategies. Note that, for the sake of computational efficiency, only nine models were in the ensemble of models trained for simulations using the large UniRep- and LSTM-derived encodings, while 22 were in the ensemble for all others. (A) The normalized discounted cumulative gain (NDCG) of the 2000 simulations for each encoding shown as violin plots. The dashed line is the median NDCG from simulations using one-hot encoding. More informative encodings tend to outperform one-hot encoding in terms of NDCG. (B) Expectation values over the 2000 simulations for the NDCG, maximum fitness of the top 96 predictions, and mean fitness of the top 96 predictions. The LSTM achieves the highest expected NDCG and mean fitness of the top 96 predictions while physicochemical encoding (Georgiev) achieves the highest expected maximum fitness. Encoding using physicochemical parameters was the only strategy tested that achieved a higher expected maximum fitness than one-hot encoding.

It is not immediately clear why the improved NDCG using learned embeddings did not translate to a higher maximum fitness achieved. These results may align, however, with the recent observations of Biswas *et al.*, who proposed that embeddings generated from unsupervised models guide a subsequent supervised search away from sequences the unsupervised model deems to be “unnatural”.^30^ In other words, when using embeddings derived from an unsupervised model for protein encoding, sequences different from those used to train that model are less likely to be predicted as fit; stronger evidence must be presented by the labeled data to identify fit “unnatural” sequences. Indeed, from homology analysis (Supporting Information: *EVcouplings Alignments*), we see that the sequence motifs defining the highest-fitness GB1 variants are not well represented in related sequences, and so could be considered “un-natural” by the unsupervised models used to generate the encodings used here. Assuming this rationale to be true, we would thus expect MLDE using embeddings to be more effective at identifying the highest-fitness variants on protein fitness landscapes where those variants are more similar to “natural” sequences. Using embeddings from larger models trained on more sequences such as that provided by Rives *et al.* could also potentially improve the effectiveness of embeddings by including more motifs in the model training data and allowing the model to learn a richer representation of protein sequences.^31,32^ Further investigation into the generalizability of embeddings for use in predicting beneficial “unnatural” mutations (both in MLDE and otherwise) appears to be emerging as an important direction for future work.

### Models/Training Procedures More Tailored for Combinatorial Fitness Landscapes Improve MLDE Predictive Performance

Many of the learned embeddings used in the previous section are extremely high-dimensional, with the largest (LSTM) describing each combination with 8192 features (Supporting Information: *Encoding Preparation*). To better handle the high dimensionality introduced by learned embeddings, in our new implementation of MLDE we added two 1D convolutional neural network (CNN) architectures to the ensemble of models trained (Supporting Information: *Inbuilt Models*). CNNs rely on spatial dependencies of high-dimensional input data to extract the most relevant high-level features.^33^ Most commonly, they are applied to image processing tasks, where 2D convolutions are used to extract high-level features from local groupings of pixels. CNNs can also be applied, however, to sequential data such as protein and DNA sequences, where 1D convolutions extract high-level features from nearby members of the sequence.^10,20,34^ Indeed, recent evidence suggests that 1D CNNs are a particularly effective model class for protein engineering.^10^ For the simulations presented in the previous section, we found that the 1D CNNs defined in MLDE were particularly effective for the highest-dimensional encodings, where they were consistently among the top-ranking models in terms of cross-validation error during training (Supporting Figures S13–S19, Supporting Table S6).

In addition to 1D CNN architectures, we also integrated XGBoost models trained with the Tweedie regression objective to better handle the zero-inflated nature of fitness landscapes. To explain, most mutations are detrimental to protein activity, and as more mutations are made to a protein the probability that it will still fold and function drops off significantly.^12^ The result is that combinatorial fitness landscapes tend to be dominated by proteins with zero or extremely low fitness,^13,14^ something that is highlighted by the distribution of fitness for GB1 (Figure 3A). Gradient-boosted Tweedie regression was developed to handle regression for datasets with zero-inflated labels that cannot be monotonically transformed to normality.^19,35^ In the simulations discussed in the previous section, the ensemble of models trained contained XGBoost models with both a linear and tree base model, each trained with either the Tweedie regression objective or the default root mean squared error objective (four models total). Models trained with the Tweedie objective consistently achieved a higher NDCG than models trained with the default objective regardless of base model and encoding (Supporting Figure S20); only models with a tree base model consistently showed improved max and mean fitness in the top M predictions, however (Supporting Figures S21–S34).

**Figure 3.**
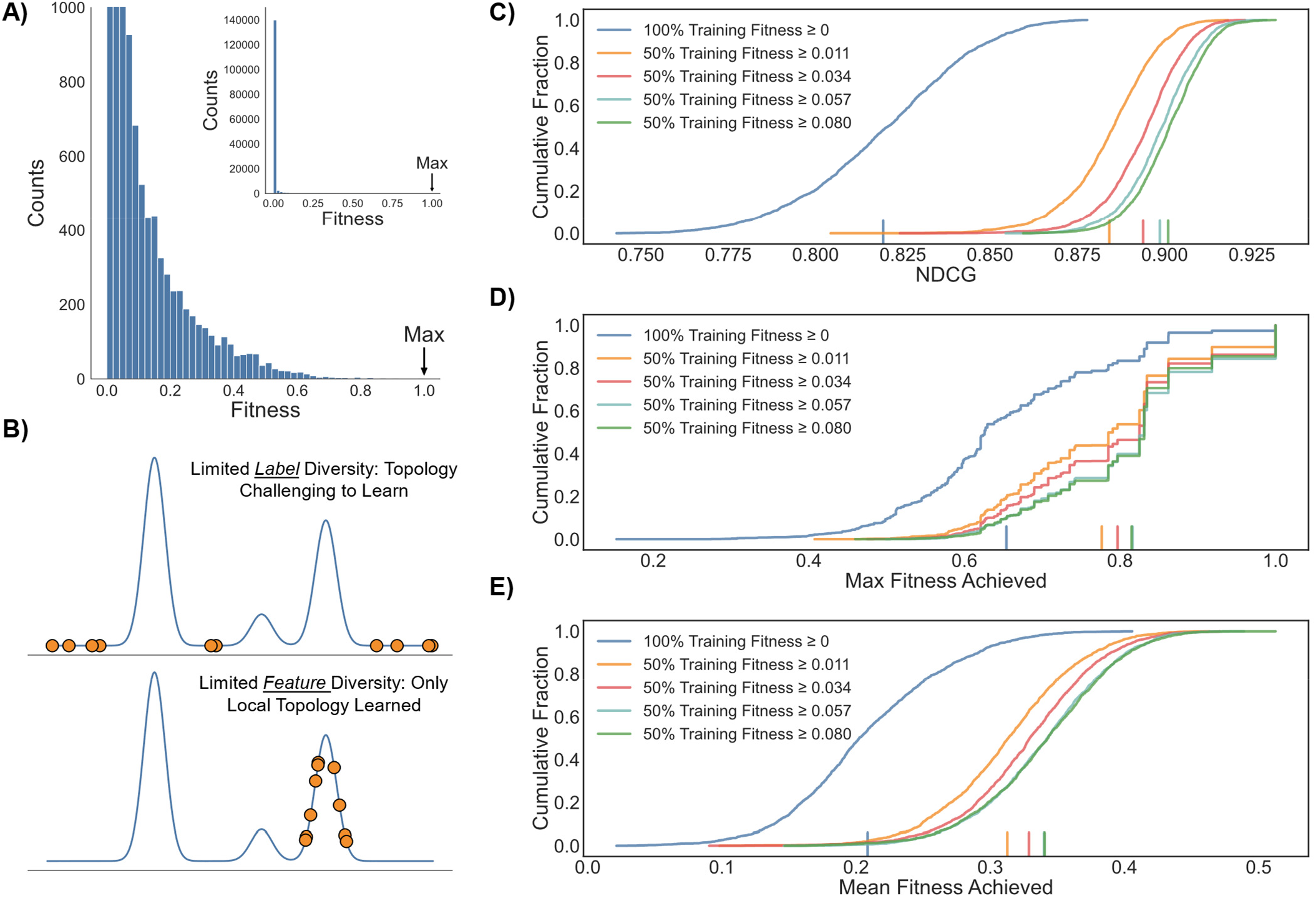
(A) The distribution of fitness in the GB1 landscape shown as a histogram. Most variants in this epistatic landscape have extremely low fitness, and the highest-fitness variants are very rare. (B) A demonstration of the importance of diversity in both the labels and features of training data for machine learning. Learning detailed topology is challenging if the labels are not representative of it, even if sampled from diverse regions of feature space. Only local topology can reliably be learned if points are sampled from a restricted region of feature space. (C) The NDCG values for simulated ftMLDE using training data sampled with simulated classifiers of varying strength. Simulated classifiers enforce that at least 50% of training data is from non-zero regions of protein space, thus committing limited training resources to regions more likely to contain the highest-fitness protein variants. Stronger classifiers enforce a higher mean training fitness. All data is shown as empirical cumulative distribution functions (ECDFs); vertical lines on the x-axis give the expectation value of the distribution. (D) The maximum fitness achieved for simulated ftMLDE using training data sampled with simulated classifiers of varying strength. (E) The mean fitness of the top 96 predictions for simulated ftMLDE using training data sampled with simulated classifiers of varying strength.

### The Challenge of Holes in Combinatorial Fitness Landscapes and the Importance of Informative Training Data

Diversity within training data is critical to constructing an effective machine learning model. Often, training set diversity is thought of in terms of exploration of the feature space, where limited resources are intelligently committed to minimize the amount of extrapolation that must be performed when making predictions. Equally important, however, is diversity in the labels; patterns in the ground truth will not be identified if there are no patterns in the training data (Figure 3B). The overabundance of “dead” (zero- or very low-fitness) variants in combinatorial fitness landscapes thus poses an additional challenge beyond that discussed in the previous section: a random draw for the generation of training data is likely to be populated by primarily zero- or extremely low-fitness variants. While useful for classifying dead vs functional proteins, these “holes” provide no information about the extent to which specific combinations of mutations benefit or harm fitness, and so have limited utility when training the regression models used in MLDE.

We thus propose a general strategy of running MLDE with training sets designed to contain a minimal number of holes. In this strategy, which we call “focused training MLDE” (ftMLDE), training data is not randomly drawn from the full combinatorial landscape (which will return primarily holes), but instead drawn from diverse regions of sequence space believed to contain functional variants. Romero *et al.* used such an approach while building ML models to predict P450 thermo-stability, where a classifier was used to build a training dataset enriched in functional protein variants.^14^ To demonstrate the concept and test the effectiveness of ftMLDE, we simulated a series of classifiers that, with 50% accuracy, could identify variants above fitness thresholds of 0.011, 0.034, 0.057, and 0.080 (Methods: *High-Fitness Simulations*). To avoid “cheating” by inclusion of the highest-fitness variants in the training data, we enforced that our classifiers could never recommend variants with fitness greater than 34% of the global maximum. We then used these simulated classifiers to generate training data enriched in functional, but not the fittest, protein variants (Supporting Figure S35), which was in turn used to perform simulated ftMLDE. Performing 2000 rounds of simulated ftMLDE using these classifiers (Methods: *High-Fitness Simulations*) for training data generation showed greatly improved NDCG, mean fitness achieved in the top 96 predictions, and max fitness achieved in the top 96 predictions compared to standard MLDE performed with randomly sampled training data (Figure 3C–E, Supporting Table S7). Notably, the difference in expected results when comparing the weakest and strongest classifiers for training data generation (at 0.017 ΔNDCG, 0.028 Δmean of top 96, and 0.038 Δmax of top 96) was less than the difference in results when comparing the weakest classifier and no classifier for training data generation (at 0.082 ΔNDCG, 0.132 Δmean of top 96, 0.161 Δmax of top 96). This result suggests that, while achieving a higher mean fitness in the training data is beneficial to ftMLDE, the more important factor is elimination of holes. In turn, this suggests that ftMLDE using even the weakest training set design predictors can achieve superior results than standard MLDE.

### Predicted ΔΔG of Stabilization for the Design of Fitness-Enriched Training Data

We next wanted to demonstrate a practical approach to building training data enriched in higher-fitness variants, thus allowing a practical demonstration of ftMLDE. One way to accomplish this would be to take an active learning approach, where a diverse set of higher-fitness variants is identified from a round of standard MLDE and then used to train models in a round of ftMLDE. Indeed, a strategy like this was taken by Romero *et al.* when evolving for improved P450 thermostability.^14^ While this active learning approach has proven success, it adds an additional round of data collection to the workflow, which is undesirable. We thus chose to investigate zero-shot prediction strategies for training set design.

We define zero-shot prediction strategies as those capable of predicting protein fitness without the need for further labeled training data collection, and thus they do not affect the overall screening burden of ftMLDE. While a number of zero-shot strategies exist—ranging from scoring protein variants based on evolutionary sequence conservation,^36–38^ to computational modeling (e.g. prediction of ΔΔG upon mutation)^39–43^ and generative modeling,^38,44,45^ to name a few—the optimal strategy will vary by protein.^46^ For instance, we attempted to build multiple sequence alignments (MSAs) to the GB1 protein sequence by querying the UniRef100 database, but found too few homologs to build an MSA with high coverage of all mutated positions in the GB1 landscape, thus disfavoring the use of evolutionary conservation approaches (Supporting Information: *EVcouplings Alignments*). In contrast, the fitness of GB1 is considered to be, at least in part, a function of stability, suggesting that approaches like predicted ΔΔG of protein stability upon mutation might be more effective.^15,47^ Indeed, we find a correlation between single-mutant fitness data and literature GB1 ΔΔG data (|Spearman ρ| = 0.58, Supporting Figure S36A).^48^ Wu *et al.* also previously presented evidence suggesting that predicted ΔΔG could be correlated to GB1 fitness.^15^

To test the effectiveness of ΔΔG predictions as a zero-shot predictor for GB1 fitness, we used the Triad protein design software suite (Protabit, Pasadena, CA, USA: https://triad.protabit.com/) with a Rosetta energy function to predict the stability of each of the 149,361 GB1 variants with measured fitness, then calculated a predicted ΔΔG of stabilization for each variant relative to the parent amino acid sequence (Supporting Information: *ΔΔG Calculations*). Both fixed backbone and flexible backbone calculations were performed using a previously determined GB1 crystal structure (PDB: 2GI9) as a scaffold.^49^ The predicted ΔΔGs from each calculation correlated with literature values of experimentally determined ΔΔG values for the single mutants, though the fixed backbone calculations were more effective (Spearman ρ = 0.61 for fixed backbone, Spearman ρ = 0.42 for flexible backbone, Supporting Figure S36B–C). Interestingly, despite both approaches having predictive power for single mutant ΔΔG, only the fixed backbone calculations were effective at identifying GB1 variants enriched in fitness when ranking by predicted ΔΔG (|Spearman ρ| = 0.27, Figure 4, Supporting Figures S37–S39). If instead, however, the GB1 variants were ranked by root mean squared deviation (RMSD) of variant structures produced during flexible backbone calculations, those variants with the lowest RMSD tended to be enriched in fitness, though not as strongly as in the fixed backbone calculations (Supporting Figure S40).

**Figure 4.**
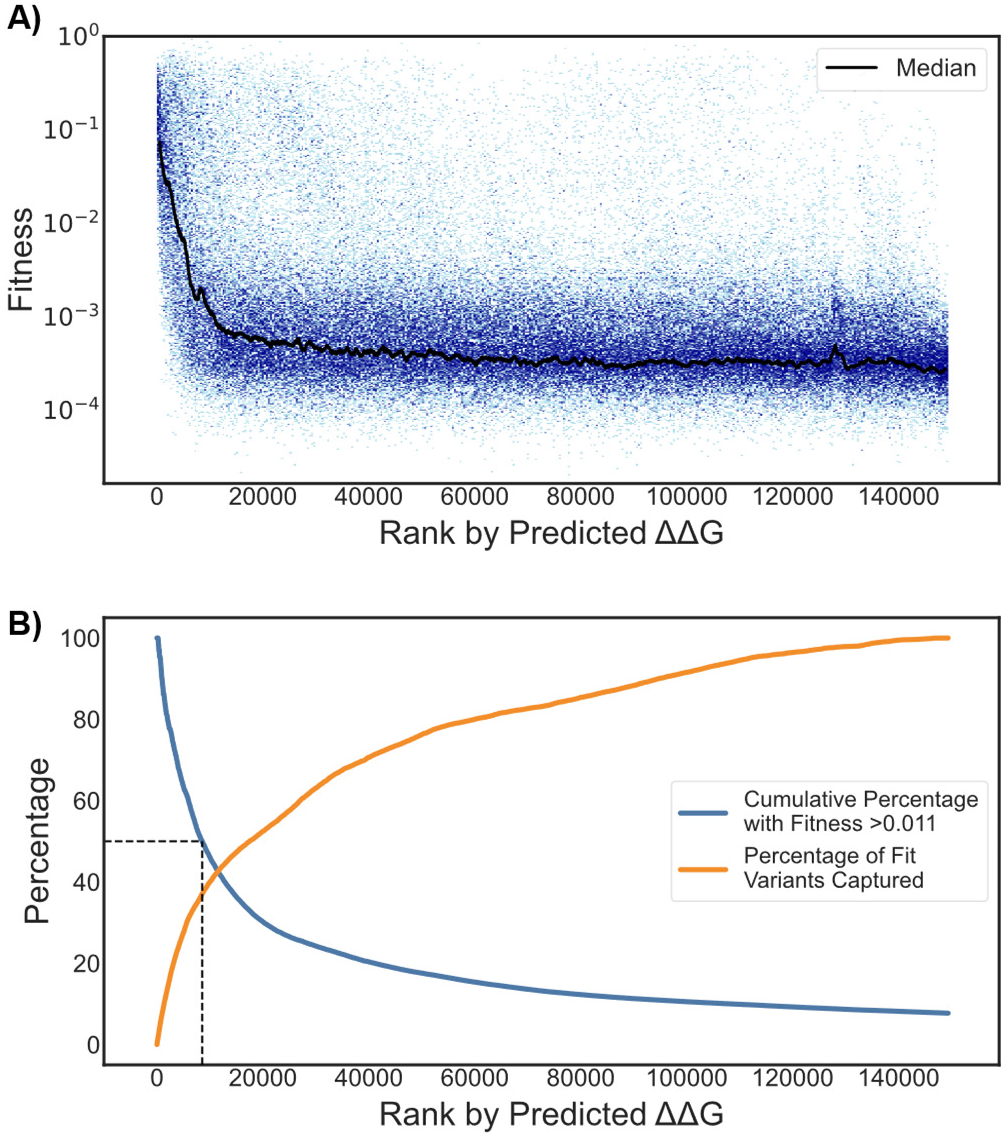
Results of zero-shot prediction using fixed backbone Triad ΔΔG of protein stability calculations. (A) The fitness of all GB1 variants ranked by predicted ΔΔG, where lower ΔΔG (and so lower rank) corresponds to a more energetically favorable mutation. Blue dots are all individual variants while the black line is the sliding median (window size = 1000) of fitness. (B) Cumulative fitness metrics for the Triad zero-shot predictions. The blue curve gives the percentage of variants ranked up to and including a given Triad rank that have fitness greater than 0.011 (the cutoff of the weakest simulated classifier in Figure 3). The orange curve gives the percentage of all “fit” (defined as fitness greater than 0.011) variants encompassed in the set up to and including a given Triad rank. The dotted line marks the first rank (8545) at which the Triad calculations have weaker classification power than the weakest simulated classifier in Figure 3. Sets up to the ranks after and including this cutoff have weaker classification power than the weakest simulated classifier and sets up to the ranks before this cutoff have stronger or equal.

Structurally conservative mutations are generally less likely to disrupt protein function, and so the observation that RMSD can be used for zero-shot prediction is not entirely surprising. Because fixed backbone calculations will tend to heavily penalize mutations that would require significant backbone movement to stabilize, an interesting question arises over the extent to which structural conservation or accurate prediction of ΔΔG allows effective fixed backbone zero-shot prediction of fitness in GB1. If structural conservation dominates, it is possible that Triad could be used for zero-shot prediction with other combinatorial libraries in proteins where variant fitness is not related to stability, particularly when the mutations in question are tightly packed together and/or buried in the protein core as they are for the GB1 landscape used in this study (Supporting Figure S41). Answering this question and evaluating the generalizability of fixed backbone Triad calculations is beyond the scope of this work, but as more fully combinatorial datasets become available this question should be investigated in more detail.

### Predicted ΔΔG of Stabilization for Training Set Design Enables Highly Effective ftMLDE on the GB1 Landscape

As a final test, we evaluated the performance of ftMLDE using GB1 training data predicted to be higher in fitness by the Triad ΔΔG calculations. To begin, we generated training data by randomly sampling 2000 training sets of 24, 48, and 384 variants from the top 1600 (1.1%), 3200 (2.1%), 6400 (4.3%), 9600 (6.4%), 12,800 (8.6%), 16,000 (10.7%), and 32,000 (21.4%) of variants as ranked by predicted ΔΔG (21 “Triad training conditions” in total, each made up of 2000 training sets); completely random training data (i.e. from the full landscape) were also drawn so that standard MLDE could be performed as a control (three additional training conditions). Predictive algorithms (Triad calculations included) will tend to predict that similar sequences have similar fitness, so sampling from different percentiles of the top predictions explores the exploration-exploitation tradeoff of using zero-shot predictions for training set design. In other words, sampling from a larger top percentile of the ranked variants allows greater sequence diversity in the training data (thus potentially enabling exploration of more fitness peaks as depicted in Figure 3B) at the expense of confidence that the variants will have non-zero fitness (Supporting Figure S42). Different sample sizes are tested to enable comparison of ftMLDE (and standard MLDE) with the most efficient implementation of traditional DE. To explain, throughout this work we have used 384 training samples with 96 tested predictions to evaluate simulations on a scale that approximates the typical experimental screening burdens for standard DE approaches.^11^ In principle, traditional DE could be performed on a four-site library by deterministically evaluating all 20 amino acids at each position, requiring only 80 measurements for the GB1 landscape. Due to the cost of synthesizing variants individually, this approach is rarely taken, and researchers instead opt to stochastically sample from pools of mutants (thus raising the required screening burden above 80). However, here we wish to directly compare the *algorithms* of traditional DE and ftMLDE; use of 24- and 48-variant training sets (with 56 and 32 tested predictions, respectively) allows for direct comparison of ftMLDE and this most efficient implementation of traditional DE.

For each of the 24 training conditions, simulated MLDE was performed using each training set (Methods: *Zero-Shot Simulations*). Importantly, after prediction, only the top-predicted unsampled combinations that could be constructed by recombining combinations in the training data were evaluated (e.g. if “AAAA” and “CCCC” were the only training examples, then only “AAAC”, “AACC”, “CAAA”, etc. could be in the top M proteins chosen for fitness evaluation). This approach enforces a confidence threshold on our predictions and focuses all resources on regions believed to contain the highest-fitness protein variants. The distributions of the achieved mean and max fitness for the simulations are shown in Figure 5 and summary statistics are given in Supporting Table S8. Most notably, when using any Triad training condition, all simulations at all screening burdens achieved both a higher expected maximum fitness as well as reached the global fitness optimum at a higher rate than traditional DE (Methods: *Traditional Directed Evolution Simulations*); ftMLDE with Triad training sets is thus a superior evolution strategy to traditional DE on the fitness landscape used here. The results when training on 384 samples are particularly impressive, with the global maximum achieved 15% of the time with training data from the top 1600 Triad-ranked samples, 92% for the top 3200, 82% for the top 6400, 77% for the top 9600, 67% for the top 12,800, 45% for the top 16,000, 30% for the top 32,000, and 8.8% for pure random sampling. By comparison, traditional DE reached the global optimum just 1.2% of the time. The difference in these rates highlights the importance of balancing exploration and exploitation, with the most effective sampling strategy having neither the highest-fitness training data nor the most sequence-diverse training data (Supporting Figure S42).

**Figure 5.**
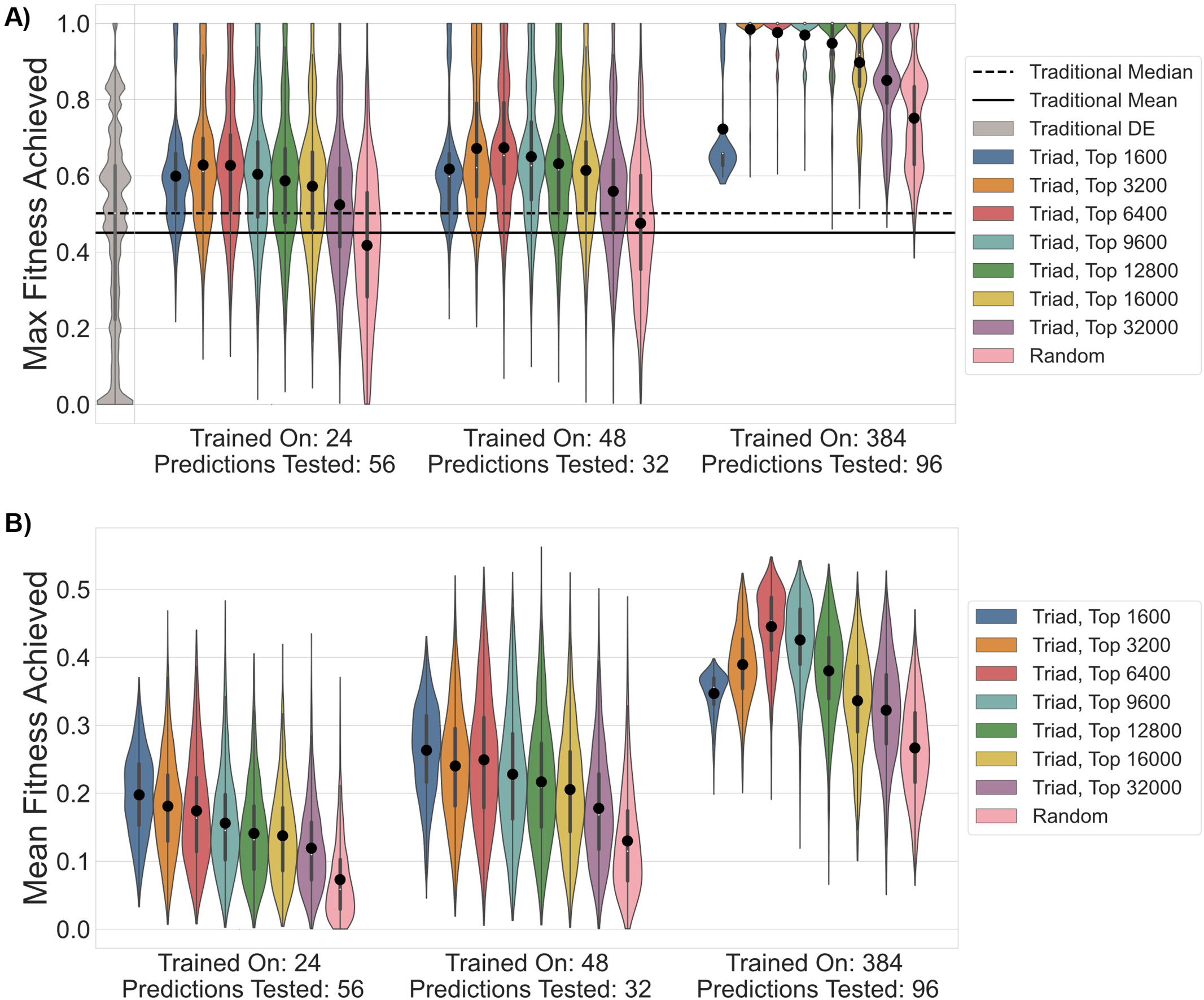
The results of simulated ftMLDE using training data generated from Triad zero-shot predictions shown as violin plots. Different colors distinguish sampling strategies (e.g. “Triad, 6400” means that training data was generated by randomly sampling from the top 6400 variants as predicted by Triad zero-shot calculations); different x-axis categories give the screening burden for the set of simulations; the large black circles give the expectation values of each distribution. (A) The max fitness achieved for the different sampling strategies and screening burdens. The simulated results of traditional DE are given on the far left in grey; the expectation value for traditional DE is given as a solid black line and the median is given as a dotted black line. For an equivalent screening burden, ftMLDE outcompetes traditional DE on the GB1 landscape after incorporation of zero-shot learning for training data selection; without zero-shot learning (standard MLDE, pink distributions) the results are comparable. (B) The mean fitness achieved in the tested predictions for the different sampling strategies and screening burdens.

### MLDE Software Enables Wet-Lab Application

To facilitate further development of MLDE, as well as to allow for its practical wet-lab application, we developed the MLDE software package, available on the Arnold Lab GitHub (https://github.com/fhalab/MLDE). This repository contains Python scripts for (1) generating any of the encodings presented in this work for any combinatorial library and (2) performing MLDE as described in this work using any encoding strategy (used in this work or otherwise). The software package was designed for use by non-computational and non-ML experts and can be executed with a simple command line call—all that is required for execution is a fasta file with the parent protein sequence and a csv file of combination-fitness data for training.

## Conclusion

We have demonstrated a series of improvements to MLDE that, all together, make it a more efficient process than the lowest-possible-screening-burden form of DE for navigating an epistatic, hole-filled, combinatorial protein fitness landscape. While incorporation of more informative encodings and models/regression strategies more amenable to combinatorial protein fitness landscapes proved beneficial to MLDE outcome, by far the most significant improvement came from training set design strategies. Specifically, we show that a focused training MLDE (ftMLDE) strategy that uses some type of predictor to avoid uninformative ex-tremely-low-fitness variants in the training data is more capable than standard MLDE at identifying the most-fit variants in a combinatorial landscape. From simulated experiments, we note that the predictor used for training set design in ftMLDE does not need to be strong— ftMLDE with even the weakest simulated classifier tested was more effective than standard MLDE—nor does it need to be capable of identifying particularly high-fitness variants—eliminating holes in the training data had a larger effect on outcome than subsequently raising training data mean fitness. The ability of the predictor to identify diverse sequences, however, is important for improving the probability of identifying the global maximum of a combinatorial landscape. This concept is highlighted when using predicted ΔΔG of protein stability as a zero-shot strategy for building training sets with functional GB1 variants, where a balance between sequence diversity and sequence fitness in the training data proved important for maximizing ftMLDE effectiveness.

There is, of course, no guarantee that predicted ΔΔG would be effective for other proteins or other functions, though we do note evidence that, by favoring conformationally conservative mutations, this strategy could generalize to other combinatorial libraries. Choosing an optimal training set design strategy will depend on the protein, and while we have mainly discussed unsupervised zero-shot strategies in this work,^36–45^ many alternate strategies can be imagined. For instance, if a protein scaffold has been used in previous protein engineering studies, a crude non-computational approach would be to avoid mutations that previously destroyed protein function. More robustly, a transfer learning approach could be taken, where an ML model trained using information from related experiments. (e.g. evolution of the same protein for a different task, evolution of a different protein for the same task, or even data from previous rounds of MLDE at different positions) is used to predict the effects of mutations in the present experiment.^50^ Perhaps even more effectively, fitness information from single-site saturation mutagenesis or error-prone PCR random mutagenesis libraries could be used to predict the fitness of combinations. Indeed, Biswas *et al.* and Hie *et al.* each recently demonstrated approaches where ML models trained on single-site or random mutation data were capable of predicting the fitness of combinations of those mutations.^30,51^ The use of Gaussian processes in the application of Hie *et al.* is particularly interesting, as it enables use of the upper confidence bound algorithm to explicitly balance exploration and exploitation, thus providing a more principled way to inject sequence diversity into training set design while maintaining high fitness.^51,52^

Whatever training set design approach is taken, we would expect its impact on the outcome of ftMLDE to be specific to the shape and makeup of the fitness landscape. For instance, on a non-epistatic landscape, minimalistic traditional DE will deterministically reach the global (and only) fitness maximum; in this case, ftMLDE could at best perform as well as traditional DE regardless of the training set design strategy used (though it may still be able to do so with a lower screening burden). Similarly, as the number of holes in a landscape increases, the probability of a random draw returning primarily uninformative zero-fitness variants increases, and so implementation of an effective training set design strategy will have a greater impact. Thus, the effectiveness of ftMLDE will vary as a function of the shape of the landscape, the number of holes in the landscape, and the availability of robust training set design strategies. We cannot expect that ftMLDE will always outcompete traditional DE.

Thorough evaluation of the effectiveness of ftMLDE will only be possible once more combinatorial landscape data beyond that provided by the GB1 landscape become available. For now, however, ftMLDE can most confidently be used on combinatorial landscapes known to be highly epistatic and that either contain few holes or else for which confident training set design strategies can be employed. The strategies, concepts, and technology presented in this work will serve as a foundation for further evaluation of the generalizability of different encodings, model architectures, regression strategies, and training set design strategies for ftMLDE on combinatorial fitness landscapes. By achieving the GB1 global maximum up to 92% of the time with a total screening burden of 470 protein variants, or up to 9.6% of the time with a screening burden of just 80 variants, the ftMLDE protocol presented here significantly outcompeted both traditional DE—which achieved the global optimum just 1.2% of the time—and our original implementation—which achieved the global optimum 8.5% of the time with a screening burden of 570 variants.^11^ This work thus presents a significant advance over our previous publication and is, to the best of our knowledge, the first proven example of a machine learning approach directly outcompeting minimalistic DE. Given the degree to which ftMLDE outcompetes traditional DE on the GB1 landscape, we hope for many more examples to come.

## Methods

### Encoding Comparison Simulations

The simulation procedure for comparing encoding strategies was designed to enable pairwise comparison of simulation results using the different encodings. For a given simulation, each of the tested encodings shared the same training set (same variant identities) and cross validation indices. A random seed was not used for model training, however, so there may be some variance introduced by models that rely on randomness for training.

All encoding comparison simulations were run with a randomly drawn training set of 384 variants (drawn without replacement) and five-fold cross validation. Except for the simulations run with LSTM- and UniRep-derived encodings, all 22 inbuilt MLDE models were trained; due to the computational expense of performing calculations on the very large LSTM- and UniRep-derived encodings, only the XGBoost and Keras models were run for simulations using them (See Supporting Information: *Inbuilt Models* for architectures and default parameter values). Trained models were then ranked according to their cross-validation mean squared error (MSE) and the predictions of the top three were averaged to predict the fitness of the remaining 148,977 variants (Supporting Information: *MLDE Programmatic Implementation* for details on model averaging). The values of NDCG, max achieved fitness, and mean achieved fitness reported for this set of simulations are all based on these predictions.

### High-Fitness Simulations

Simulated classifiers have two parameters: “cutoff” and “limit”. When used for sampling, 50% of variants chosen must have a fitness greater than or equal to the value of “cutoff” and 50% must have a fitness below. Cutoff values of 0.011, 0.034, 0.057, and 0.080 were used to define four different classifiers in this study. “Limit” gives the highest allowed fitness value sampled from the GB1 dataset. Throughout this study, “limit” was set to 0.34 for all simulated classifiers.

To generate training data using a simulated classifier, the GB1 dataset was first filtered to exclude all variants with fitness greater than “limit”. The remaining data was then split into two sets: one set had all variants with fitness greater than or equal to the threshold and the other set had all variants with fitness less than the threshold. Equal numbers of samples were then drawn at random from the two sets without replacement. Training data for the “no classifier” control discussed in the results section and presented in Figure 3C-E (“100% Training Fitness ≥ 0”) was generated by sampling at random from the limit-filtered GB1 dataset.

For each of the four simulated classifiers and the no classifier control, 2000 training sets were generated, each containing 384 variants (10,000 training sets of 384 variants in total). Each training set was then fed into the simulated MLDE pipeline using Georgiev parameters for variant encoding and 5-fold cross validation. Trained models were then ranked according to their cross-validation MSE and the predictions of the top three were averaged to predict the fitness of the unlabeled variants. The values of NDCG, max achieved fitness, and mean achieved fitness reported for this set of simulations are all based on these predictions.

### Zero-Shot Simulations

To generate training data using predicted ΔΔG, the GB1 dataset was first ranked by Triad score (from lowest to highest predicted ΔΔG). We defined ΔΔG such that a lower predicted ΔΔG corresponded to a predicted more energetically favorable mutation. Next, the top 1600 variants (i.e. the 1600 variants with the lowest predicted ΔΔG) were identified, and 2000 random samples of 384 were drawn at random without replacement. This process was repeated for the top-ranked 3200, 6400, 9600, 12,800, 16,000, and 32,000 variants, resulting in 14,000 total training sets, each containing 384 random samples.

For the 384-training-sample simulations, each of the 14,000 training sets was then fed into the simulated MLDE pipeline. For the sake of computational efficiency, only CPU-bound (scikit-learn and XGBoost) models were evaluated. All simulations were performed using Georgiev parameters for variant encoding and 5-fold cross validation. Trained models were ranked according to their cross-validation MSE and the predictions of the top three were averaged to predict the fitness of the remaining variants. Only the top-predicted un-sampled combinations that could be constructed by recombining combinations in the training data were evaluated, enforcing a confidence threshold on our predictions and focusing all resources on regions believed to contain the highest-fitness protein variants. The reported values of max achieved fitness and mean achieved fitness reported for this set of simulations are all derived from this restricted set of evaluated proteins, though the “fitness” value returned is still normalized to the full unsampled set. For a given simulation, the global maximum is considered to be achieved if it is present in either the training data or the evaluated predictions. The random controls presented in the results and in Figure 5 are derived from the *Encoding Comparison Simulations* when using Georgiev encodings, but only evaluating the CPU models and employing the same confidence threshold strategy for evaluating predictions.

For the 24- and 48-training-sample simulations, the first 24 and 48 variants in each of the full 384-sample training sets were used for training, respectively. For random controls, the first 24 and 48 variants from the full 384-sample training sets used in the *Encoding Comparison Simulations* were used for training. Otherwise, the procedure was the same as for the 384-training-sample simulations.

### Traditional Directed Evolution Simulations

Traditional DE simulations were performed from every variant in the GB1 landscape with non-zero starting fitness. Zero-fitness variants were omitted from these simulations as a researcher would never begin a DE study from such a variant. As in the MLDE simulations, variants with imputed fitness in the GB1 dataset were ignored for these simulations.

A greedy walk simulation begins with 4 potential positions to evaluate. One of these positions is selected, the fitness values of all mutants at this position are evaluated, and the best mutation is fixed. In the next round, there are three positions to evaluate. One of these positions is selected, all mutants are evaluated, and the best mutation is fixed again. This process continues until all positions have been evaluated; the fitness of the best variant identified in the last round is returned. The results reported for the greedy walk simulations consider all possible paths from all non-zero-fitness starting variants (with 24 paths per starting variant and 119,814 non-zero fitness starting points, this is 2,877,216 simulated greedy walks in total).

## Acknowledgments

The authors thank Patrick Almhjell, Lucas Schaus, and Sabine Brinkmann-Chen for helpful discussion and critical reading of the manuscript, Zachary Wu, Kadina Johnston, and Amir Motmaen for helpful discussion, and Paul Chang for assistance with Triad calculations. Additionally, the authors thank NVIDIA Corporation for donation of two Titan V GPUs used in this work as well as Amazon.com Inc. for donation of AWS computing credits. This work was supported by the NSF Division of Chemical, Bioengineering, Environmental and Transport Systems (CBET 1937902), and by an Amgen Chem-Bio-Engineering Award (CBEA).

## References

(1) Maynard Smith, J. Natural Selection and the Concept of a Protein Space. Nature 1970, 225, 563–564. https://doi.org/10.1038/225563a0.

(2) Romero, P. A.; Arnold, F. H. Exploring Protein Fitness Landscapes by Directed Evolution. Nat. Rev. Mol. Cell Biol. 2009, 10, 866–876. https://doi.org/10.1038/nrm2805.

(3) Fasan, R.; Kan, S. B. J.; Zhao, H. A Continuing Career in Biocatalysis: Frances H. Arnold. ACS Catal. 2019, 9, 9775–9788. https://doi.org/10.1021/acscatal.9b02737.

(4) Starr, T. N.; Thornton, J. W. Epistasis in Protein Evolution. Protein Sci. 2016, 25, 1204–1218. https://doi.org/10.1002/pro.2897.

(5) Miton, C. M.; Tokuriki, N. How Mutational Epistasis Impairs Predictability in Protein Evolution and Design. Protein Sci. 2016, 25, 1260–1272. https://doi.org/10.1002/pro.2876.

(6) Kaznatcheev, A. Computational Complexity as an Ultimate Constraint on Evolution. Genetics 2019, 212, 245–265. https://doi.org/10.1534/genetics.119.302000.

(7) Yang, K. K.; Wu, Z.; Arnold, F. H. Machine-Learning-Guided Directed Evolution for Protein Engineering. Nat. Methods 2019, 16, 687–694. https://doi.org/10.1038/s41592-019-0496-6.

(8) Mazurenko, S.; Prokop, Z.; Damborsky, J. Machine Learning in Enzyme Engineering. ACS Catal. 2020, 10, 1210–1223. https://doi.org/10.1021/acscatal.9b04321.

(9) Siedhoff, N. E.; Schwaneberg, U.; Davari, M. D. Machine Learning-Assisted Enzyme Engineering. In Methods in Enzymology; Elsevier Inc., 2020; Vol. 643, 281–315. https://doi.org/10.1016/bs.mie.2020.05.005.

(10) Xu, Y.; Verma, D.; Sheridan, R. P.; Liaw, A.; Ma, J.; Marshall, N. M.; McIntosh, J.; Sherer, E. C.; Svetnik, V.; Johnston, J. M. Deep Dive into Machine Learning Models for Protein Engineering. J. Chem. Inf. Model. 2020, 60, 2773–2790. https://doi.org/10.1021/acs.jcim.0c00073.

(11) Wu, Z.; Kan, S. B. J.; Lewis, R. D.; Wittmann, B. J.; Arnold, F. H. Machine Learning-Assisted Directed Protein Evolution with Combinatorial Libraries. Proc. Natl. Acad. Sci. 2019, 116, 8852–8858. https://doi.org/10.1073/pnas.1901979116.

(12) Bloom, J. D.; Silberg, J. J.; Wilke, C. O.; Drummond, D. A.; Adami, C.; Arnold, F. H. Thermodynamic Prediction of Protein Neutrality. Proc. Natl. Acad. Sci. 2005, 102, 606–611. https://doi.org/10.1073/pnas.0406744102.

(13) Arnold, F. The Library of Maynard-Smith: My Search for Meaning in the Protein Universe. Microbe 2011, 6, 316–318. https://doi.org/10.1128/microbe.6.316.1.

(14) Romero, P. A.; Krause, A.; Arnold, F. H. Navigating the Protein Fitness Landscape with Gaussian Processes. Proc. Natl. Acad. Sci. 2013, 110, E193–E201. https://doi.org/10.1073/pnas.1215251110.

(15) Wu, N. C.; Dai, L.; Olson, C. A.; Lloyd-Smith, J. O.; Sun, R. Adaptation in Protein Fitness Landscapes Is Facilitated by Indirect Paths. Elife 2016, 5. https://doi.org/10.7554/eLife.16965.

(16) Georgiev, A. G. Interpretable Numerical Descriptors of Amino Acid Space. J. Comput. Biol. 2009, 16, 703–723. https://doi.org/10.1089/cmb.2008.0173.

(17) Rao, R.; Bhattacharya, N.; Thomas, N.; Duan, Y.; Chen, X.; Canny, J.; Abbeel, P.; Song, Y. S. Evaluating Protein Transfer Learning with TAPE. arXiv 2019. arXiv:1906.08230.

(18) Chen, T.; Guestrin, C. XGBoost: A Scalable Tree Boosting System. arXiv 2016. arXiv:1603.02754.

(19) Zhou, H.; Qian, W.; Yang, Y. Tweedie Gradient Boosting for Extremely Unbalanced Zero-Inflated Data. Commun. Stat. - Simul. Comput. 2020, 1–23. https://doi.org/10.1080/03610918.2020.1772302.

(20) Bai, S.; Kolter, J. Z.; Koltun, V. An Empirical Evaluation of Generic Convolutional and Recurrent Networks for Sequence Modeling. arXiv 2018. arXiv:1803.01271.

(21) Buitinck, L.; Louppe, G.; Blondel, M.; Pedregosa, F.; Mueller, A.; Grisel, O.; Niculae, V.; Prettenhofer, P.; Gramfort, A.; Grobler, J.; Layton, R.; Vanderplas, J.; Joly, A.; Holt, B.; Varoquaux, G. API Design for Machine Learning Software: Experiences from the Scikit-Learn Project. arXiv 2013. arXiv:1309.0238.

(22) Kawashima, S.; Pokarowski, P.; Pokarowska, M.; Kolinski, A.; Katayama, T.; Kanehisa, M. AAindex: Amino Acid Index Database, Progress Report 2008. Nucleic Acids Res. 2008, 36, 202–205. https://doi.org/10.1093/nar/gkm998.

(23) Ofer, D.; Linial, M. ProFET: Feature Engineering Captures High-Level Protein Functions. Bioinformatics 2015, 31, 3429–3436. https://doi.org/10.1093/bioinformatics/btv345.

(24) Vaswani, A.; Shazeer, N.; Parmar, N.; Uszkoreit, J.; Jones, L.; Gomez, A. N.; Kaiser, Ł.; Polosukhin, I. Attention Is All You Need. arXiv 2017. arXiv:1706.03762.

(25) Alley, E. C.; Khimulya, G.; Biswas, S.; AlQuraishi, M.; Church, G. M. Unified Rational Protein Engineering with Sequence-Based Deep Representation Learning. Nat. Methods 2019, 16, 1315–1322. https://doi.org/10.1038/s41592-019-0598-1.

(26) Bepler, T.; Berger, B. Learning Protein Sequence Embeddings Using Information from Structure. arXiv 2019. arXiv:1902.08661.

(27) Hochreiter, S.; Schmidhuber, J. Long Short-Term Memory. Neural Comput. 1997, 9, 1735–1780. https://doi.org/10.1162/neco.1997.9.8.1735.

(28) Yu, F.; Koltun, V.; Funkhouser, T. Dilated Residual Networks. arXiv 2017. arXiv:1705.09914.

(29) El-Gebali, S.; Mistry, J.; Bateman, A.; Eddy, S. R.; Luciani, A.; Potter, S. C.; Qureshi, M.; Richardson, L. J.; Salazar, G. A.; Smart, A.; Sonnhammer, E. L. L.; Hirsh, L.; Paladin, L.; Piovesan, D.; Tosatto, S. C. E.; Finn, R. D. The Pfam Protein Families Database in 2019. Nucleic Acids Res. 2019, 47, D427–D432. https://doi.org/10.1093/nar/gky995.

(30) Biswas, S.; Khimulya, G.; Alley, E. C.; Esvelt, K. M.; Church, G. M. Low-N Protein Engineering with Data-Efficient Deep Learning. bioRxiv 2020. https://doi.org/10.1101/2020.01.23.917682.

(31) Rives, A.; Meier, J.; Sercu, T.; Goyal, S.; Lin, Z.; Guo, D.; Ott, M.; Zitnick, C. L.; Ma, J.; Fergus, R. Biological Structure and Function Emerge from Scaling Unsupervised Learning to 250 Million Protein Sequences. bioRxiv 2020. https://doi.org/10.1101/622803.

(32) Brown, T. B.; Mann, B.; Ryder, N.; Subbiah, M.; Kaplan, J.; Dhariwal, P.; Neelakantan, A.; Shyam, P.; Sastry, G.; Askell, A.; Agarwal, S.; Herbert-Voss, A.; Krueger, G.; Henighan, T.; Child, R.; Ramesh, A.; Ziegler, D. M.; Wu, J.; Winter, C.; Hesse, C.; Chen, M.; Sigler, E.; Litwin, M.; Gray, S.; Chess, B.; Clark, J.; Berner, C.; McCandlish, S.; Radford, A.; Sutskever, I.; Amodei, D. Language Models Are Few-Shot Learners. arXiv 2020. arXiv:2005.14165.

(33) Rawat, W.; Wang, Z. Deep Convolutional Neural Networks for Image Classification: A Comprehensive Review. Neural Comput. 2017, 29, 2352–2449. https://doi.org/10.1162/neco_a_00990.

(34) Jaganathan, K.; Panagiotopoulou, S. K.; McRae, J. F.; Darbandi, S. F.; Knowles, D.; Li, Y. I.; Kosmicki, J. A.; Arbelaez, J.; Cui, W.; Schwartz, G. B.; Chow, E. D.; Kanterakis, E.; Gao, H.; Kia, A.; Batzoglou, S.; Sanders, S. J.; Farh, K. K.-H. Predicting Splicing from Primary Sequence with Deep Learning. Cell 2019, 176, 535–548. https://doi.org/10.1016/j.cell.2018.12.015.

(35) Yang, Y.; Qian, W.; Zou, H. Insurance Premium Prediction via Gradient Tree-Boosted Tweedie Compound Poisson Models. J. Bus. Econ. Stat. 2018, 36, 456–470. https://doi.org/10.1080/07350015.2016.1200981.

(36) Ng, P. C.; Henikoff, S. SIFT: Predicting Amino Acid Changes That Affect Protein Function. Nucleic Acids Res. 2003, 31, 3812–3814. https://doi.org/10.1093/nar/gkg509.

(37) Hopf, T. A.; Ingraham, J. B.; Poelwijk, F. J.; Schärfe, C. P. I.; Springer, M.; Sander, C.; Marks, D. S. Mutation Effects Predicted from Sequence Co-Variation. Nat. Biotechnol. 2017, 35, 128–135. https://doi.org/10.1038/nbt.3769.

(38) Riesselman, A. J.; Ingraham, J. B.; Marks, D. S. Deep Generative Models of Genetic Variation Capture the Effects of Mutations. Nat. Methods 2018, 15, 816–822. https://doi.org/10.1038/s41592-018-0138-4.

(39) Firnberg, E.; Labonte, J. W.; Gray, J. J.; Ostermeier, M. A Comprehensive, High-Resolution Map of a Gene’s Fitness Landscape. Mol. Biol. Evol. 2014, 31, 1581–1592. https://doi.org/10.1093/molbev/msu081.

(40) Jacquier, H.; Birgy, A.; Le Nagard, H.; Mechulam, Y.; Schmitt, E.; Glodt, J.; Bercot, B.; Petit, E.; Poulain, J.; Barnaud, G.; Gros, P.-A.; Tenaillon, O. Capturing the Mutational Landscape of the Beta-Lactamase TEM-1. Proc. Natl. Acad. Sci. U. S. A. 2013, 110, 13067–13072. https://doi.org/10.1073/pnas.1215206110.

(41) Sirin, S.; Apgar, J. R.; Bennett, E. M.; Keating, A. E. AB-Bind: Antibody Binding Mutational Database for Computational Affinity Predictions. Protein Sci. 2016, 25, 393–409. https://doi.org/10.1002/pro.2829.

(42) Sarkisyan, K. S.; Bolotin, D. A.; Meer, M. V.; Usmanova, D. R.; Mishin, A. S.; Sharonov, G. V.; Ivankov, D. N.; Bozhanova, N. G.; Baranov, M. S.; Soylemez, O.; Bogatyreva, N. S.; Vlasov, P. K.; Egorov, E. S.; Logacheva, M. D.; Kondrashov, A. S.; Chudakov, D. M.; Putintseva, E. V.; Mamedov, I. Z.; Tawfik, D. S.; Lukyanov, K. A.; Kondrashov, F. A. Local Fitness Landscape of the Green Fluorescent Protein. Nature 2016, 533, 397–401. https://doi.org/10.1038/nature17995.

(43) Yang, J.; Naik, N.; Patel, J. S.; Wylie, C. S.; Gu, W.; Huang, J.; Ytreberg, F. M.; Naik, M. T.; Weinreich, D. M.; Rubenstein, B. M. Predicting the Viability of Beta-Lactamase: How Folding and Binding Free Energies Correlate with Beta-Lactamase Fitness. PLoS One 2020, 15. https://doi.org/10.1371/journal.pone.0233509.

(44) Riesselman, A.; Shin, J.-E.; Kollasch, A.; McMahon, C.; Simon, E.; Sander, C.; Manglik, A.; Kruse, A.; Marks, D. Accelerating Protein Design Using Autoregressive Generative Models. bioRxiv 2019. https://doi.org/10.1101/757252.

(45) Madani, A.; McCann, B.; Naik, N.; Keskar, N. S.; Anand, N.; Eguchi, R. R.; Huang, P.-S.; Socher, R. ProGen: Language Modeling for Protein Generation. arXiv 2020. arXiv:2004.03497.

(46) Livesey, B. J.; Marsh, J. A. Using Deep Mutational Scanning to Benchmark Variant Effect Predictors and Identify Disease Mutations. Mol. Syst. Biol. 2020, 16. https://doi.org/10.15252/msb.20199380.

(47) Olson, C. A.; Wu, N. C.; Sun, R. A Comprehensive Biophysical Description of Pairwise Epistasis throughout an Entire Protein Domain. Curr. Biol. 2014, 24, 2643–2651. https://doi.org/10.1016/j.cub.2014.09.072.

(48) Nisthal, A.; Wang, C. Y.; Ary, M. L.; Mayo, S. L. Protein Stability Engineering Insights Revealed by Domain-Wide Comprehensive Mutagenesis. Proc. Natl. Acad. Sci. 2019, 116, 16367–16377. https://doi.org/10.1073/pnas.1903888116.

(49) Franks, W. T.; Wylie, B. J.; Stellfox, S. A.; Rienstra, C. M. Backbone Conformational Constraints in a Microcrystalline U-15N-Labeled Protein by 3D Dipolar-Shift Solid-State NMR Spectroscopy. J. Am. Chem. Soc. 2006, 128, 3154–3155. https://doi.org/10.1021/ja058292x.

(50) Shamsi, Z.; Chan, M.; Shukla, D. TLmutation: Predicting the Effects of Mutations Using Transfer Learning. bioRxiv 2020. https://doi.org/10.1101/2020.01.07.897892.

(51) Hie, B.; Bryson, B.; Berger, B. Learning with Uncertainty for Biological Discovery and Design. bioRxiv 2020. https://doi.org/10.1101/2020.08.11.247072.

(52) Srinivas, N.; Krause, A.; Kakade, S.; Seeger, M. Gaussian Process Optimization in the Bandit Setting: No Regret and Experimental Design. arXiv 2010. arXiv:0912.3995.

