## Supporting Information for "Machine Learning-Assisted Directed Evolution Navigates a Combinatorial Epistatic Fitness Landscape with Minimal Screening Burden"

### MLDE Software

This section provides details on the MLDE software used in this work. It also provides information on the MLDE software made available at https://github.com/fhalab/MLDE.

#### Encoding Preparation

We investigated 3 encoding classes in this work: One-hot, physicochemical parameters, and learned embeddings. One-hot encodings were prepared by first assigning each amino acid an index. To encode each variant, a 4×20 matrix filled with 0's was instantiated where the index of each row corresponded to the amino acid index in the variant. Each row of the matrix was then populated with a single value of "1" at the index corresponding to the appropriate amino acid.

Physicochemical encodings were prepared using the descriptors originally published by Georgiev,[1] using the values found in code published by Ofer & Linial.[2] To encode all variants, a 160,000×4×19 tensor was instantiated ("N possible combos" × "N amino acids per combo" × "N Georgiev parameters"). Every possible variant was encoded using all 19 Georgiev parameters, and the encodings were stored in the instantiated tensor. The last two dimensions of the tensor were then flattened to produce a 160,000×76 matrix, and each column of the matrix was mean-centered and unit-scaled. The final encoding tensor was generated by extracting only those rows belonging to GB1 variants with experimentally measured fitness, then reshaping the last dimension to produce a 149,361×4×19 tensor.

Learned embeddings were prepared using the pre-trained models published by Rao *et al.*[3–5] The full amino acid sequences of all 160,000 possible GB1 variants at positions 39, 40, 41, and 54 were first constructed and stored in fasta file format. The template sequence was:

```
MQYKLILNGKTLKGETTTEAVDAATAEKVFKQYANDNGVDGEWTYDDATKTFTVTE
```

This fasta file was passed into the software associated with the original publication by Rao *et al.* (https://github.com/songlab-cal/tape-neurips2019) to generate tensors of shape 160,000×56×L ("N possible combos" × "Length of GB1" × "L latent dimensions"). The value of L varied by model used, and is given in Table S1. Next, the indices corresponding to amino acids varied in the GB1 dataset (indices 38, 39, 40, and 53, using 0-indexing) were extracted from the output tensor. Using the same procedure as with the physicochemical properties, the resultant 160,000×4×L tensor was mean-centered and unit-scaled to produce a 160,000×4L matrix. Finally, the appropriate rows were isolated and the last dimension reshaped to produce a final learned embedding encoding tensor of shape 149,361×4×L.

**Table S1.** The number of latent dimensions (number of features describing each amino acid) in the encodings generated by each of the models provided by Rao *et al.*

| Model | Latent Dimensions |
|---|---|
| Bepler | 100 |
| ResNet | 256 |
| Transformer | 512 |
| UniRep | 1900 |
| LSTM | 2048 |

#### MLDE Programmatic Implementation

This section details the machine learning portion of MLDE. Scripts to execute MLDE can be found on the Arnold Lab GitHub (https://github.com/fhalab/MLDE).

The MLDE algorithm takes as input all encodings corresponding to the combinations of amino acids found in the training data along with their measured fitness values. During the training stage, these sampled combinations are used to train a version of all inbuilt models (see *Inbuilt Models*, below) using k-fold cross validation and the default model parameters; mean validation error from the k-fold cross validation is recorded. Using the same k-fold splits, in the next stage, H rounds of Bayesian hyperparameter optimization are optionally performed using the package "hyperopt".[6] Hyperparameters that minimize mean validation error are recorded. Note that, due to computational limitations, hyperparameter optimization was skipped for all simulations presented in the main text, and models trained on default model values were used. For making predictions, the top N model architectures (those with the lowest cross-validation error after hyperparameter optimization) are first identified. For each of the top N model architectures, predictions are made on the unsampled combinations by averaging the predictions of the k models trained during cross validation. The predictions made by each of the N models are then averaged to return a single final prediction of the unsampled values. In total, this means that $k \times N$ models are averaged to generate a single prediction for MLDE (k from each of the top N model architectures).

#### Inbuilt Models

##### *Keras*

Five separate neural network architectures were implemented using the Python package Keras: three fully connected neural network architectures and two 1D-convolutional neural network architectures. Summaries of each Keras model initialized with default MLDE hyperparameters and using the encodings derived from the LSTM model used in this work are given in Figures S1–S5. Note that both the layer sizes and the number of filters were defined as a fraction of the number of latent dimensions in the encoding. As a result, the total number of parameters varied by encoding used, with lower-dimensionality encodings having fewer. The identities and default values of tunable hyperparameters are given in Table S2. All neural networks were trained with a batch size of 32 using the “adam” optimizer for at most 1000 epochs with early stopping after 10 epochs with no improvement in validation error (calculated against the cross-validation test data).

The fully connected neural networks differed in the number of hidden layers: zero, one, or two. After each hidden layer, a batch normalization layer was employed, followed by a rectified linear unit (ReLu) nonlinearity. A single dropout layer was used before the output layer. The output layer was a scalar value passed through a ReLu nonlinearity.

The convolutional neural networks differed in the number 1D convolutional layers: one or two. After each convolutional layer, a batch normalization layer was employed followed by a ReLu nonlinearity. Following the convolutional layers, the output matrix was flattened with a GlobalAveragePooling1D layer. After flattening, a single dropout layer was used before the output layer. The output layer was a scalar value passed through a ReLu nonlinearity.

```
_________________________________________________________________
Layer (type)                 Output Shape              Param #
=================================================================
dropout_1 (Dropout)          (None, 8192)              0
_________________________________________________________________
dense_1 (Dense)              (None, 1)                 8193
=================================================================
Total params: 8,193
Trainable params: 8,193
Non-trainable params: 0
```

**Figure S1.** Example Keras summary for the no-hidden-layers fully connected neural network. This example is defined for use with the LSTM model used in this work and with default MLDE hyperparameters.

```
_________________________________________________________________
Layer (type)                 Output Shape              Param #
=================================================================
dense_1 (Dense)              (None, 2048)              16779264
_________________________________________________________________
batch_normalization_1 (Batch (None, 2048)              8192
_________________________________________________________________
activation_1 (Activation)    (None, 2048)              0
_________________________________________________________________
dropout_1 (Dropout)          (None, 2048)              0
_________________________________________________________________
dense_2 (Dense)              (None, 1)                 2049
=================================================================
Total params: 16,789,505
Trainable params: 16,785,409
Non-trainable params: 4,096
```

**Figure S2.** Example Keras summary for the one-hidden-layer fully connected neural network. This example is defined for use with the LSTM model used in this work and with default MLDE hyperparameters. Note that the hidden layer size is defined as a fraction of the number of latent dimensions in the encoding, so the total number of parameters will vary.

```
_________________________________________________________________
Layer (type)                 Output Shape              Param #
=================================================================
dense_1 (Dense)              (None, 2048)              16779264
_________________________________________________________________
batch_normalization_1 (Batch (None, 2048)              8192
_________________________________________________________________
activation_1 (Activation)    (None, 2048)              0
_________________________________________________________________
dense_2 (Dense)              (None, 512)               1049088
_________________________________________________________________
batch_normalization_2 (Batch (None, 512)               2048
_________________________________________________________________
activation_2 (Activation)    (None, 512)               0
_________________________________________________________________
dropout_1 (Dropout)          (None, 512)               0
_________________________________________________________________
dense_3 (Dense)              (None, 1)                 513
=================================================================
Total params: 17,839,105
Trainable params: 17,833,985
Non-trainable params: 5,120
```

**Figure S3.** Example Keras summary for the two-hidden-layer fully connected neural network. This example is defined for use with the LSTM model used in this work and with default MLDE hyperparameters. Note that the hidden layer sizes are defined as a fraction of the number of latent dimensions in the encoding, so the total number of parameters will vary.

```
_________________________________________________________________
Layer (type)                 Output Shape              Param #
=================================================================
conv1d_1 (Conv1D)            (None, 3, 128)            524416
_________________________________________________________________
batch_normalization_1 (Batch (None, 3, 128)            512
_________________________________________________________________
activation_1 (Activation)    (None, 3, 128)            0
_________________________________________________________________
global_average_pooling1d_1 ( (None, 128)               0
_________________________________________________________________
dropout_1 (Dropout)          (None, 128)               0
_________________________________________________________________
dense_1 (Dense)              (None, 1)                 129
=================================================================
Total params: 525,057
Trainable params: 524,801
Non-trainable params: 256
_________________________________________________________________
```

**Figure S4.** Example Keras summary for the 1D convolutional neural network with one convolutional layer. This example is defined for use with the LSTM model used in this work and with default MLDE hyperparameters. Note that the number of filters is defined as a fraction of the number of latent dimensions in the encoding, so the total number of parameters will vary.

```
_________________________________________________________________
Layer (type)                 Output Shape              Param #
=================================================================
conv1d_1 (Conv1D)            (None, 3, 128)            524416
_________________________________________________________________
batch_normalization_1 (Batch (None, 3, 128)            512
_________________________________________________________________
activation_1 (Activation)    (None, 3, 128)            0
_________________________________________________________________
conv1d_2 (Conv1D)            (None, 2, 16)             4112
_________________________________________________________________
batch_normalization_2 (Batch (None, 2, 16)             64
_________________________________________________________________
activation_2 (Activation)    (None, 2, 16)             0
_________________________________________________________________
global_average_pooling1d_1 ( (None, 16)                0
_________________________________________________________________
dropout_1 (Dropout)          (None, 16)                0
_________________________________________________________________
dense_1 (Dense)              (None, 1)                 17
=================================================================
Total params: 529,121
Trainable params: 528,833
Non-trainable params: 288
```

**Figure S5.** Example Keras summary for the 1D convolutional neural network with two convolutional layers. This example is defined for use with the LSTM model used in this work and with default MLDE hyperparameters. Note that the number of filters is defined as a fraction of the number of latent dimensions in the encoding, so the total number of parameters will vary.

**Table S2.** The tunable parameters with their default values for the different neural network architectures used in MLDE.

| Architecture | Parameter | Description | Default |
|---|---|---|---|
| NoHidden | dropout | Dropout value for model | 0.2 |
| OneHidden | dropout | Dropout value for model | 0.2 |
| OneHidden | size1 | Size of the hidden layer as a fraction of the encoding dimensionality | 0.25 |
| TwoHidden | dropout | Dropout value for model | 0.2 |
| TwoHidden | size1 | Size of the hidden layer as a fraction of the encoding dimensionality | 0.25 |
| TwoHidden | size2 | Size of the second hidden layer as a fraction of the encoding dimensionality | 0.0625 |
| OneConv | dropout | Dropout value for model | 0.2 |
| OneConv | filter_choice | The width of the 1D convolutional window as a fraction of the number of positions in the combinatorial space | 0.5 |
| OneConv | n_filters1 | The number of filters used in the convolution as a fraction of the encoding dimensionality of a single position | 0.0625 |
| OneConv | flatten_choice | The method of flattening post convolution | "Average" |
| TwoConv | dropout | Dropout value for model | 0.2 |
| TwoConv | filter_arch | The widths of the two 1D convolutional windows as a fraction of the number of positions in the combinatorial space, given as a tuple | (0.5, 0.5) |
| TwoConv | n_filters1 | The number of filters used in the first convolution as a fraction of the encoding dimensionality of a single position | 0.0625 |
| TwoConv | n_filters2 | The number of filters used in the second convolution as a fraction of the encoding dimensionality of a single position | 0.007813 |
| TwoConv | flatten_choice | The method of flattening post convolution | "Average" |

##### *XGBoost*

Four gradient boosting approaches were implemented in MLDE using the Python package XGBoost:[7] both tree and linear base models were implemented with both “reg:squarederror” and “reg:tweedie” objectives. For reg:tweedie, `tweedie_variance_power` was set to 1.5. The identities and default values for tunable XGBoost hyperparameters are given in Table S3. Unless explicitly mentioned in Table S3, all XGBoost parameters were held at their default values as detailed in the official XGBoost documentation. The descriptions for all parameters can also be found in the official XGBoost documentation. All XGBoost models used in this work were implemented with early stopping: `eval_metric` was set to “rmse” when the “reg:squarederror” was used and “tweedie-nloglik@1.5” when the “reg:tweedie” objective was used; validation error was calculated against the cross-validation test data; training was terminated if validation error did not decrease for 10 epochs or 1000 total training epochs had passed.

**Table S3.** The tunable parameters with their default values for the base models used in the XGBoost models of MLDE.

| Base Model | Parameter | Default Value |
|---|---|---|
| Linear | lambda | 1 |
| Linear | alpha | 0 |
| Tree | eta | 0.3 |
| Tree | max_depth | 6 |
| Tree | lambda | 1 |
| Tree | alpha | 0 |

##### *Scikit-learn*

Only scikit-learn models were used in our original implementation of MLDE.[8] To remain consistent with this first implementation, the regressor models from scikit-learn that were effective while using default parameters in our previous implementation were also used in this new version.[9] The identities and default values for tunable scikit-learn hyperparameters are given in Table S4. Unless explicitly mentioned in Table S4, all scikit-learn parameters were held at their default values as detailed in the official scikit-learn documentation. The descriptions for all parameters can also be found in the official documentation.

**Table S4.** The tunable parameters with their default values for the scikit-learn models used in MLDE.

| Model | Parameter | Default Value |
|---|---|---|
| Linear | N/A | N/A |
| GradientBoostingRegressor | learning_rate | See sklearn docs |
| GradientBoostingRegressor | n_estimators | See sklearn docs |
| GradientBoostingRegressor | min_samples_split | See sklearn docs |
| GradientBoostingRegressor | min_samples_leaf | See sklearn docs |
| GradientBoostingRegressor | max_depth | See sklearn docs |
| RandomForestRegressor | n_estimators | See sklearn docs |
| RandomForestRegressor | min_samples_split | See sklearn docs |
| RandomForestRegressor | min_samples_leaf | See sklearn docs |
| RandomForestRegressor | max_depth | See sklearn docs |
| LinearSVR | tol | See sklearn docs |
| LinearSVR | C | See sklearn docs |
| LinearSVR | dual | See sklearn docs |
| ARDRegression | tol | See sklearn docs |
| ARDRegression | alpha_1 | See sklearn docs |
| ARDRegression | alpha_2 | See sklearn docs |
| ARDRegression | lambda_1 | See sklearn docs |
| ARDRegression | lambda_2 | See sklearn docs |
| KernelRidge | alpha | See sklearn docs |
| KernelRidge | kernel | See sklearn docs |
| BayesianRidge | tol | See sklearn docs |
| BayesianRidge | alpha_1 | See sklearn docs |
| BayesianRidge | alpha_2 | See sklearn docs |
| BayesianRidge | lambda_1 | See sklearn docs |
| BayesianRidge | lambda_2 | See sklearn docs |
| BaggingRegressor | n_estimators | See sklearn docs |
| BaggingRegressor | max_samples | See sklearn docs |
| LassoLarsCV | max_iter | See sklearn docs |
| LassoLarsCV | cv | 5 |
| LassoLarsCV | max_n_alphas | See sklearn docs |
| DecisionTreeRegressor | max_depth | See sklearn docs |
| DecisionTreeRegressor | min_samples_split | See sklearn docs |
| DecisionTreeRegressor | min_samples_leaf | See sklearn docs |
| SGDRegressor | alpha | See sklearn docs |
| SGDRegressor | l1_ratio | See sklearn docs |
| SGDRegressor | tol | See sklearn docs |
| KNeighborsRegressor | n_neighbors | See sklearn docs |
| KNeighborsRegressor | weights | See sklearn docs |
| KNeighborsRegressor | leaf_size | See sklearn docs |
| KNeighborsRegressor | p | See sklearn docs |
| ElasticNet | l1_ratio | See sklearn docs |
| ElasticNet | alpha | See sklearn docs |
| AdaBoostRegressor | n_estimators | See sklearn docs |
| AdaBoostRegressor | learning_rate | See sklearn docs |

#### Compute Environment

All MLDE code is written in Python using Anaconda as the environment manager. The Anaconda environment "mlde.yml" within the MLDE GitHub page can be used to build an environment in which MLDE is known to be stable.

#### Supplementary Methods/Results

##### Evaluation Metrics

The ranking metrics used in this work include (1) the mean normalized true fitness of the M-highest-ranked variants, (2) the max normalized true fitness of the M-highest-ranked variants, and (3) the ranking metric "normalized discounted cumulative gain" (NDCG) of all predictions (where "gain" is defined as the normalized fitness of unsampled variants). NDCG was calculated using scikit-learn's `ndcg_score()` function, which uses the form

$$NDCG = \left(\sum_{i=1}^{N} \frac{r_i}{\log_2(i+1)}\right) \Bigg/ \left(\sum_{i=1}^{N} \frac{r_i'}{\log_2(i+1)}\right), \qquad \text{Eq. 1}$$

where $r$ is the true fitness of all ($N$) unsampled variants ranked by predicted fitness and $r'$ is the true fitness of all unsampled variants ranked by true fitness (i.e. the ideal ordering). When evaluating a single MLDE simulation, the fitness was normalized to the highest-fitness variant in the unsampled data. Typically, this was equivalent to normalizing to the highest fitness in the entire GB1 dataset, as it was extremely unlikely that the highest-fitness variant in the dataset is drawn in the training set. Still, normalizing to the highest unsampled fitness allowed us to make more fair comparisons between MLDE simulations in the rare case that the highest-fitness value appeared in the training data.

##### Computational Hardware Information

Simulations were performed across three workstations and an r5.24xlarge Amazon Web Services (AWS) EC2 instance. The specs on all workstations are given in Table S5. Triad calculations were performed on Desktop2 (fixed backbone) and a c5.24xlarge AWS EC2 instance (flexible backbone). Information regarding which computer ran which specific simulation/computation with which specific piece of hardware is available upon request.

**Table S5.** Hardware information for the three workstations used in this work.

| ComputerID | OS | CPU | GPU1 | GPU2 |
|---|---|---|---|---|
| **Desktop1** | Ubuntu 18.04.3 LTS | AMD Ryzen 9 3900X | NVIDIA GeForce RTX 2070 | NVIDIA GeForce RTX 2070 |
| **Desktop2** | Ubuntu 18.04.3 LTS | Intel i7-8700 | NVIDIA Titan V | NVIDIA GeForce RTX 2070 |
| **Desktop3** | Ubuntu 18.04.3 LTS | Intel i7-8700 | NVIDIA Titan V | NVIDIA GeForce RTX 2070 |

##### EVcouplings Alignments

Multiple sequence alignments (MSAs) were generated using the parent GB1 sequence (See *Encoding Preparation* above) and the EVcouplings webapp.[10] The alignments were performed against the UniRef100 database for bitscore inclusion thresholds of 0.10, 0.30, 0.50, and 0.70, keeping all other settings at their default values (Alignment threshold type = Bitscore; Search iterations = 5; Position filter = 70%; Sequence fragment filter = 50%; Removing similar sequences = 90%; Downweighting similar sequences = 80%). At a bitscore inclusion threshold of 0.50, this alignment returned at most 27 redundancy-reduced sequences which covered all variable positions in the GB1 landscape at ≥70% coverage. However, a bitscore sequence inclusion threshold of 0.30 returned at most 1914 redundancy-reduced sequences with ≥70% coverage for positions 39, 40, and 41 (but not 54), so we decided to fine-tune the bitscore threshold to try and increase the number of redundancy-reduced sequences covering all GB1 landscape positions with ≥70% coverage. Testing bitscores of 0.35, 0.36, 0.37, 0.38, 0.39, 0.40 and 0.45 yielded at most 63 redundancy-reduced sequences covering all positions with ≥70% coverage (at a bitscore inclusion threshold of 0.40). Given these results, we concluded that achieving the ≥10L (where L is the length of the protein) number of redundancy-reduced sequences targeted by the authors in the evolutionary conservation-based zero-shot prediction tools EVmutation and DeepSequence would not be achievable for the GB1 combinatorial landscape.[11,12]

We next evaluated the frequency of different sequence motifs in the best alignments (in terms of number of sequences returned) when considering only the first three variable positions in the GB1 landscape (positions 39,

40 and 41) as well as when considering all variable positions in the GB1 landscape. We define a "sequence motif" to be the combination of amino acids aligned to the variable positions in GB1 in the output (.a2m) alignment files of the EVcouplings webapp. At a bitscore inclusion threshold of 0.1, 17,301 redundancy-reduced sequences were returned which covered the first three positions in the GB1 landscape at ≥70% coverage, consisting of 1769 unique motifs. As expected, the most common motif was "VDG", corresponding to the wild-type GB1 protein (which has V39 D40 G41 V54). The motif with the highest associated GB1 fitness was "VAA", which can be found in the full combination "VAAA" with fitness of 0.70. Testing the alignment output for the bitscore inclusion threshold of 0.4 (the tested bitscore that yielded the most redundancy-reduced sequences covering all four positions in the GB1 landscape at ≥70% coverage), we find 29 unique sequence motifs with the most common motif ("VDGV") again corresponding to wild type. The motif with the highest associated GB1 fitness for the 0.4 bitscore alignments was "VNAA" with fitness of 0.62.

The lack of sequence motifs corresponding to the highest-fitness GB1 variants in sequences related to GB1 provides a potential explanation for why simulated MLDE using embeddings for encodings can achieve NDCG scores superior to one-hot encoding but not a higher maximum fitness. As proposed by Biswas *et al.*, unsupervised learning (and so the embeddings generated from unsupervised models) potentially serves to guide a downstream supervised search away from sequences deemed to be "unnatural", where an unnatural sequence is a sequence distant from those on which the unsupervised model was trained.[13] Because there are no direct motifs in closely related sequences to GB1 that correspond to the highest-fitness GB1 variants, it is possible that the embeddings used in this work push MLDE away from the "unnatural" highest-fitness variants. This bias toward favoring more "natural" variants need not necessarily diminish the general ranking capabilities of MLDE models trained using embeddings (thus still enabling higher-than-one-hot NDCG values), just limit their ability to identify the highest-fitness variants.

#### ΔΔG Calculations

ΔΔG calculations were performed using a local copy of the Triad software suite (Protabit, Pasadena, CA, USA: https://triad.protabit.com/). To begin, the template protein crystal structure (PDB: 2GI9) was prepared for calculations via the below command:

```
$ ~/triad-2.1.2/triad.sh ~/triad-2.1.2/apps/preparation/proteinProcess.py -struct
2GI9.pdb --crosetta
```

This command generated two files: 2GI9_process.pdb and 2GI9_prepared.pdb.[14] The "_process" pdb file is the 2GI9.pdb file prepared for downstream Triad calculations but without any structural minimization. The "_prepared" pdb file is the 2GI9.pdb file prepared for downstream Triad calculations but with an added constrained minimization. The flexible backbone calculations were run on an AWS c5.24x-large EC2 instance using the standard Rosetta scoring function and the "_prepared" pdb file. The command line call is below:

```
$ ~/triad-2.1.2/tools/openmpi/bin/mpirun -np 96 ~/triad-2.1.2/triad.sh
~/triad-2.1.2/apps/cleanSequences_BjwMod.py -struct
./2GI9_prepared.pdb -inputSequences
2GI9.mut -crosetta -calculateRmsd --minDesign -inputSequenceFormat
pid --floatNearbyResidues 2>&1 | tee $OUTPUT
```

Note that the cleanSequences_BjwMod.py file is a version of the inbuilt Triad script cleanSequences.py modified to also output root mean squared deviation (RMSD) of the protein backbone. The 'inputSequences' file '2GI9.mut' describes the mutations for all 149,360 GB1 variants relative to the parent GB1 protein. The fixed backbone calculations were run on Desktop 2 (Table S5) with the "_process" pdb file using a Rosetta scoring function that has a Van der Waals term with a softer inner wall, reducing the chance that steric clashes produce overly high energies. The command line call for the flexible backbone calculations is below:

```
$ ~/triad-2.1.2/tools/openmpi/bin/mpirun -np 12 ~/triad-2.1.2/triad.sh
~/triad-2.1.2/apps/cleanSequences.py -struct ../pdbs/2GI9_process.pdb -inputSequences
../2GI9.mut -rosetta -inputSequenceFormat pid --floatNearbyResidues -soft 2>&1 | tee
$OUTPUT
```

There was no output file directly produced by the cleanSequence.py script, hence the captured output. The captured output file generated was parsed to extract ΔG values for each protein variant, which were in turn used to calculate ΔΔG values relative to the parent protein. In this work, we defined a negative ΔΔG to be stabilizing and a positive ΔΔG to be destabilizing relative to the parent protein; this necessitated flipping the sign of the literature ΔΔG values of Nisthal *et al.*, who defined opposite meanings of the sign of ΔΔG.[15]

#### Supplementary Results/Figures

This section contains all supplementary figures and tables referenced in the main text. For simplicity, subsections are named identically to the main text; relevant supplementary figures/tables for the main text sections can be found in these supplementary information counterparts.

##### More Informative Encodings Improve MLDE Outcome

This section contains all supplementary figures for the investigation of different encodings on MLDE outcome. Because the training and cross-validation indices were kept the same between encodings for each simulation, pairwise comparisons between simulation results can be made. Figure S6 shows the pairwise NDCG scores of 2000 MLDE simulations for each encoding compared against the others. Figures S7–S12 show the distributions of the differences in fitness summary statistics over all MLDE simulations between one-hot encoding and each more-informative encoding, considering testing 1 through 96 variants with the highest predicted fitness. Specifically, in each of Figures S7–S12, the x-axis describes the number of top predictions tested (e.g. when the x-axis is "10", the boxplot above shows the distribution of differences in mean or max fitness achieved over 2000 simulations when testing the 10 unsampled variants with highest predicted fitness), while the y-axis describes the difference in fitness metrics (e.g. if one-hot achieved a maximum fitness of "0.5" in a simulation and the other encoding a maximum fitness of "0.7", the difference would be "0.2"—the results of 2000 such differences for the 2000 simulations performed make up the boxplot above each x-value considered). In each of Figures S7–S12, subplot A shows the difference in mean fitness achieved and subplot B shows the difference in maximum fitness achieved. All differences are calculated as the more informative encoding results minus the one-hot encoding results. The red line in each figure highlights "zero" (no difference in results). Simulations performed with learned embeddings tended to achieve both a higher NDCG score and mean fitness (except for UniRep-derived encodings, which achieved a higher NDCG score but lower mean fitness) than simulations performed with one-hot encodings, but a lower maximum fitness. Simulations performed with physicochemical encodings ("Georgiev") consistently achieved a higher NDCG, mean fitness, and max fitness than simulations performed with one-hot encodings.

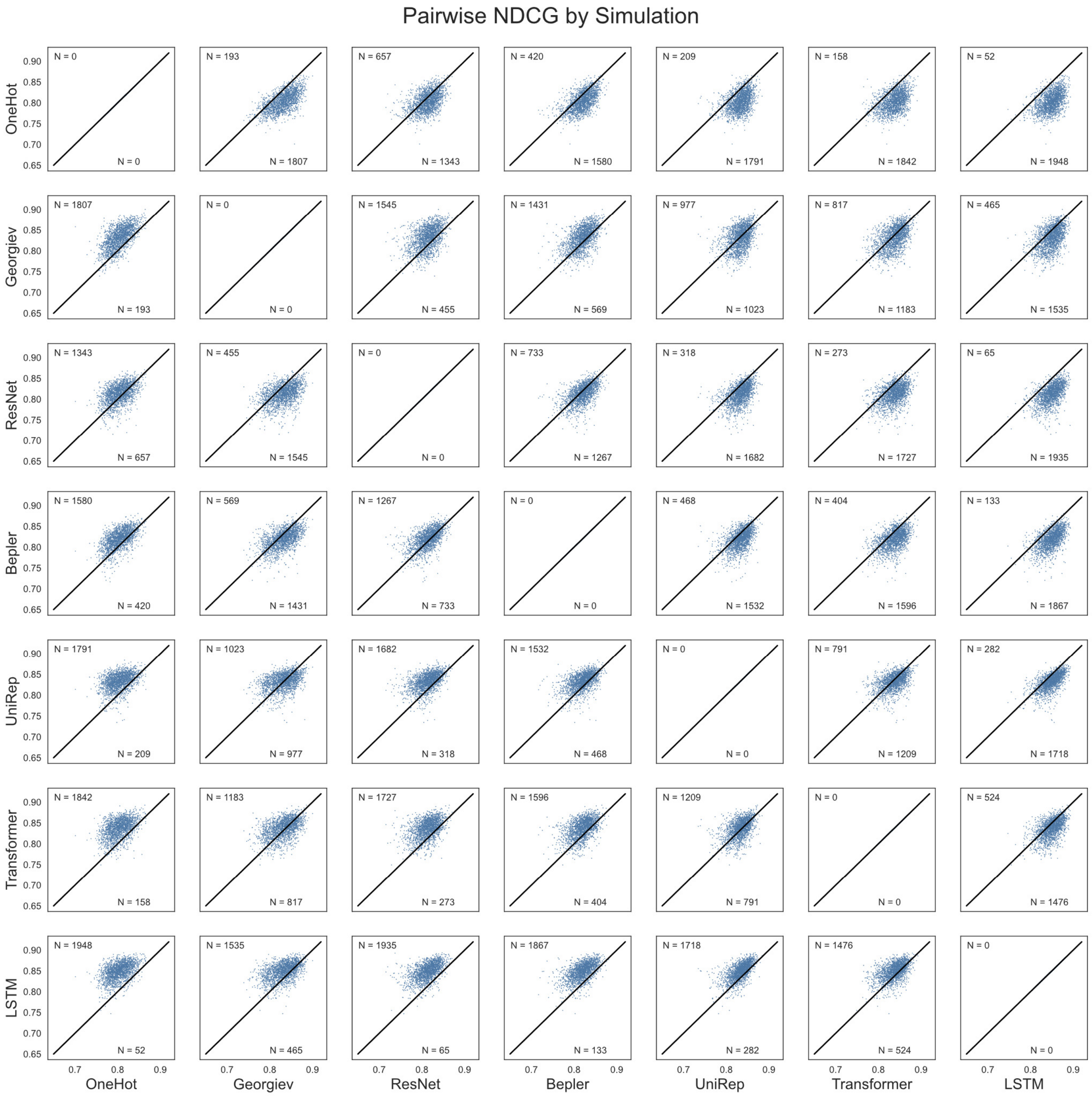

**Figure S6.** A pairwise comparison of NDCG for all encodings for each of the 2000 MLDE simulations run for comparing encoding strategies.

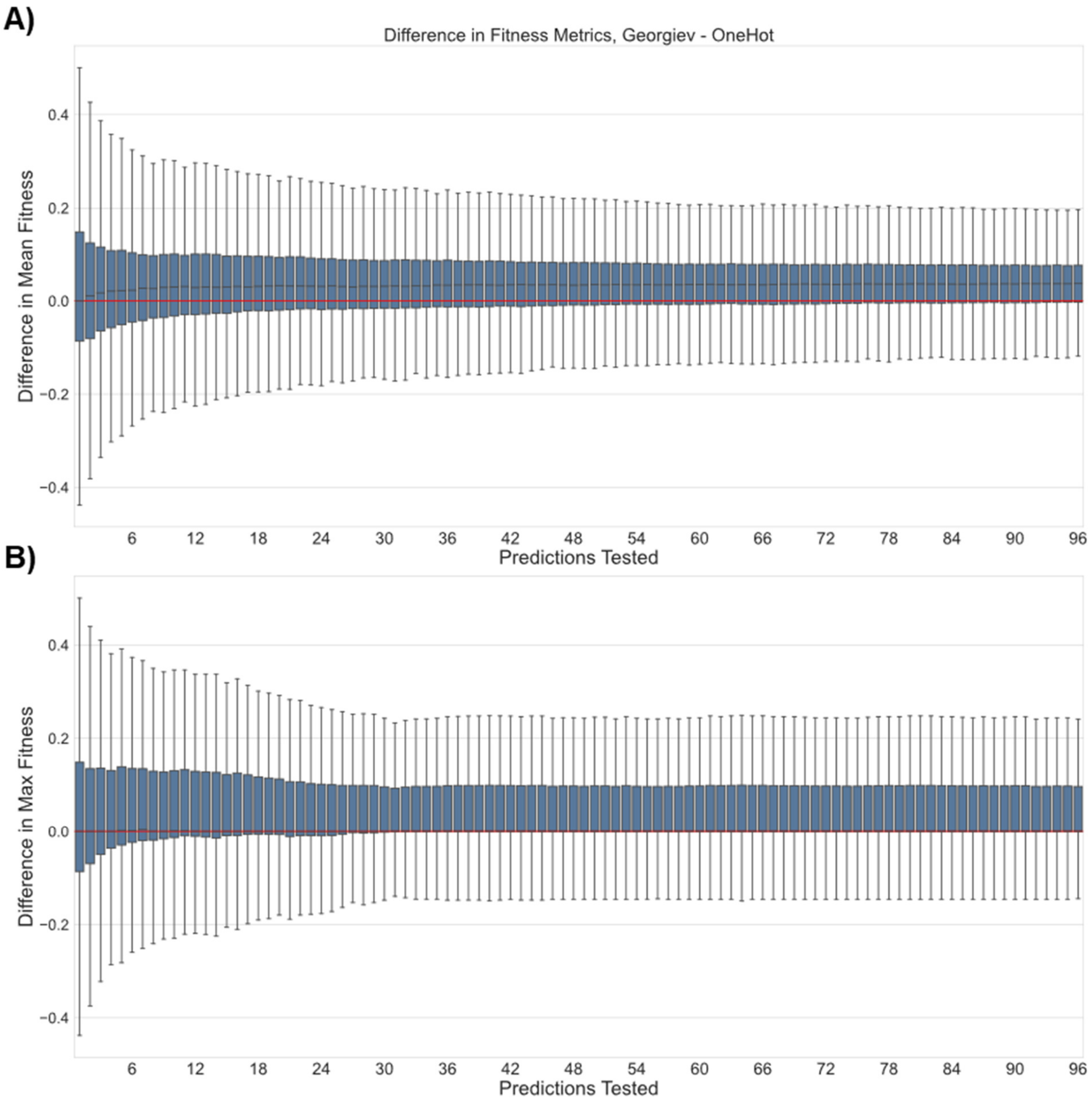

**Figure S7.** Pairwise comparison of fitness results for Georgiev and one-hot encodings over 2000 simulations when testing different numbers of the top-predicted variants. Extra details on the meaning and construction of this figure can be found in paragraph at the top of this section. The x-axis gives the number of top predictions tested (e.g. when x = 42, the 42 variants predicted to have the highest fitness by MLDE were evaluated for each encoding strategy) and the y-axis gives the difference in either (A) the mean or (B) the maximum fitness achieved in that top sample. All differences are reported as Georgiev – one-hot. The red line serves as a reference for 0 difference. Probability mass above the red line indicates a superior result using Georgiev encodings.

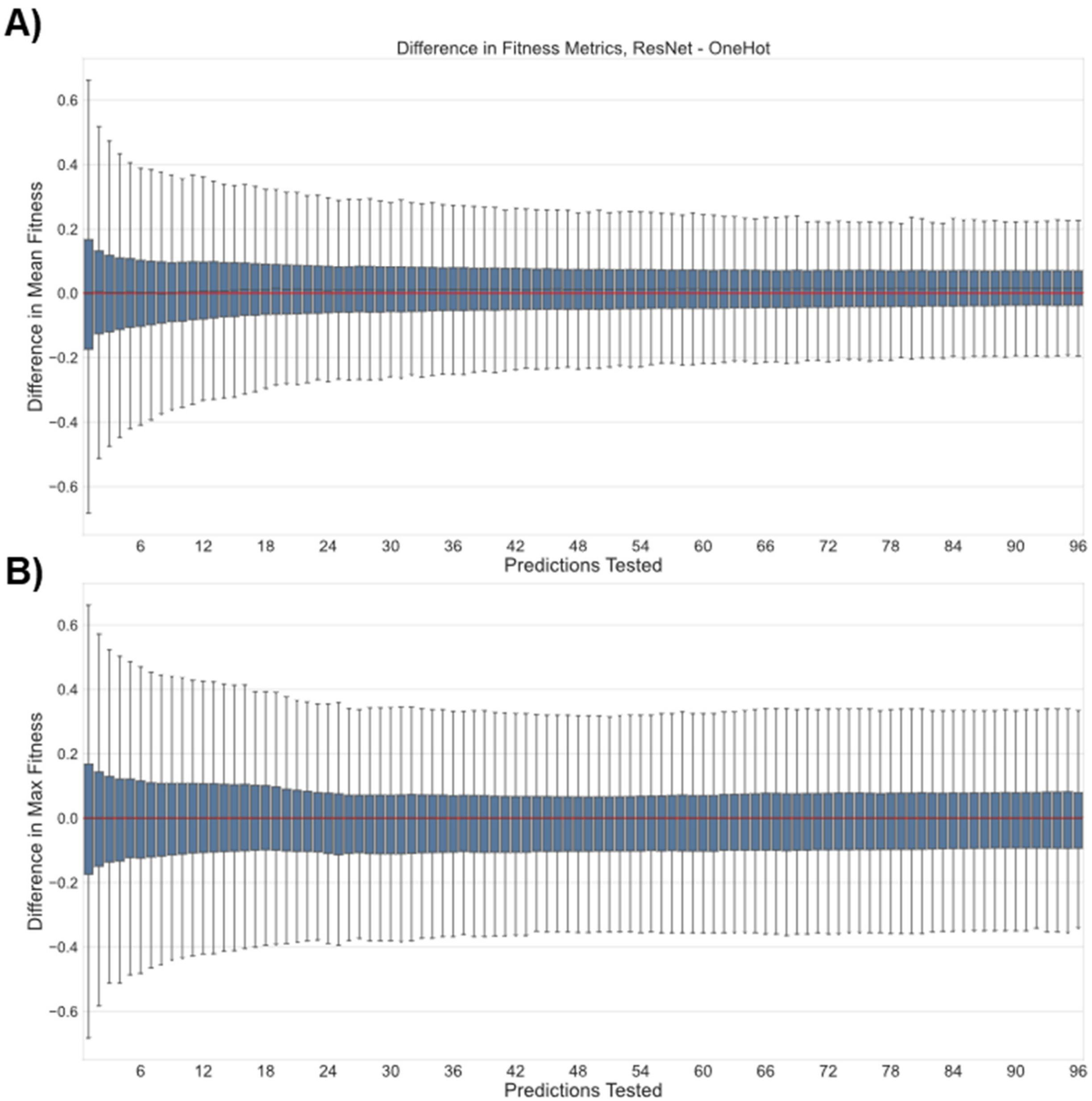

**Figure S8.** Pairwise comparison of fitness results for ResNet-derived and one-hot encodings over 2000 simulations when testing different numbers of the top-predicted variants. Extra details on the meaning and construction of this figure can be found in paragraph at the top of this section. The x-axis gives the number of top predictions tested (e.g. when x = 42, the 42 variants predicted to have the highest fitness by MLDE were evaluated for each encoding strategy) and the y-axis gives the difference in either (A) the mean or (B) the maximum fitness achieved in that top sample. All differences are reported as ResNet – one-hot. The red line serves as a reference for 0 difference. Probability mass above the red line indicates a superior result using ResNet-derived encodings.

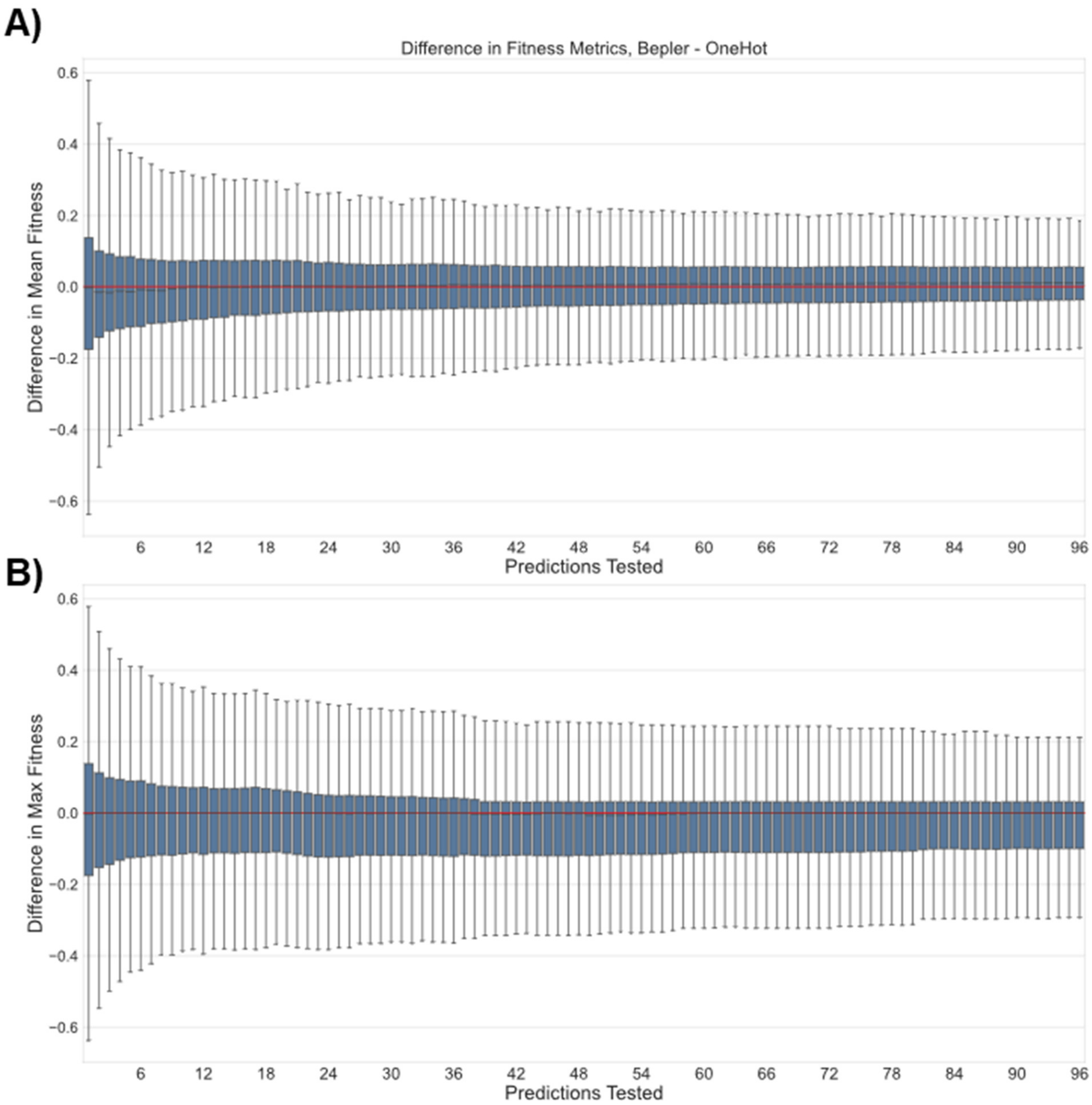

**Figure S9.** Pairwise comparison of fitness results for Bepler-derived and one-hot encodings over 2000 simulations when testing different numbers of the top-predicted variants. Extra details on the meaning and construction of this figure can be found in paragraph at the top of this section. The x-axis gives the number of top predictions tested (e.g. when x = 42, the 42 variants predicted to have the highest fitness by MLDE were evaluated for each encoding strategy) and the y-axis gives the difference in either (A) the mean or (B) the maximum fitness achieved in that top sample. All differences are reported as Bepler – one-hot. The red line serves as a reference for 0 difference. Probability mass above the red line indicates a superior result using Bepler-derived encodings.

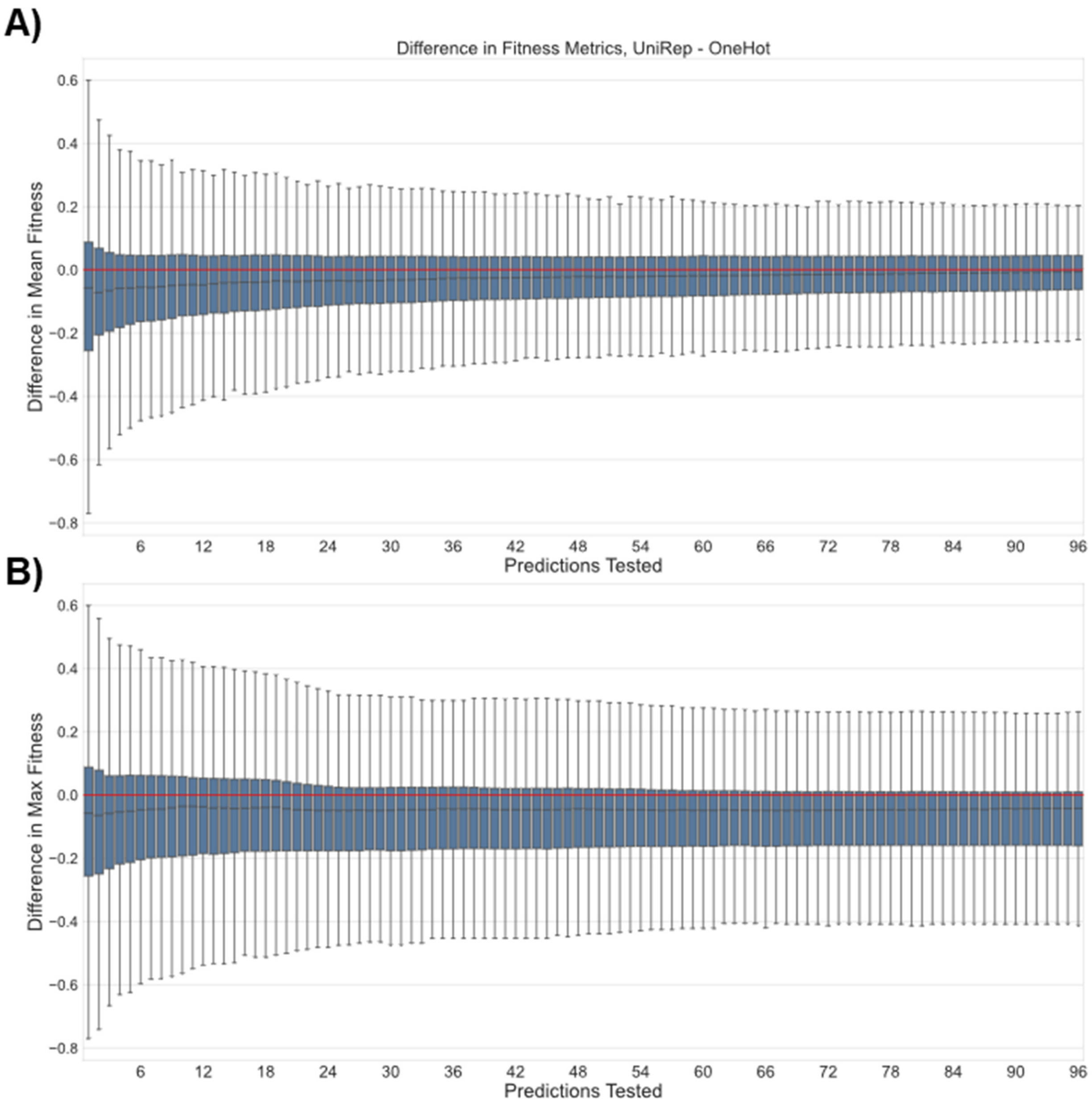

**Figure S10.** Pairwise comparison of fitness results for UniRep-derived and one-hot encodings over 2000 simulations when testing different numbers of the top-predicted variants. Extra details on the meaning and construction of this figure can be found in paragraph at the top of this section. The x-axis gives the number of top predictions tested (e.g. when x = 42, the 42 variants predicted to have the highest fitness by MLDE were evaluated for each encoding strategy) and the y-axis gives the difference in either (A) the mean or (B) the maximum fitness achieved in that top sample. All differences are reported as UniRep – one-hot. The red line serves as a reference for 0 difference. Probability mass above the red line indicates a superior result using UniRep-derived encodings.

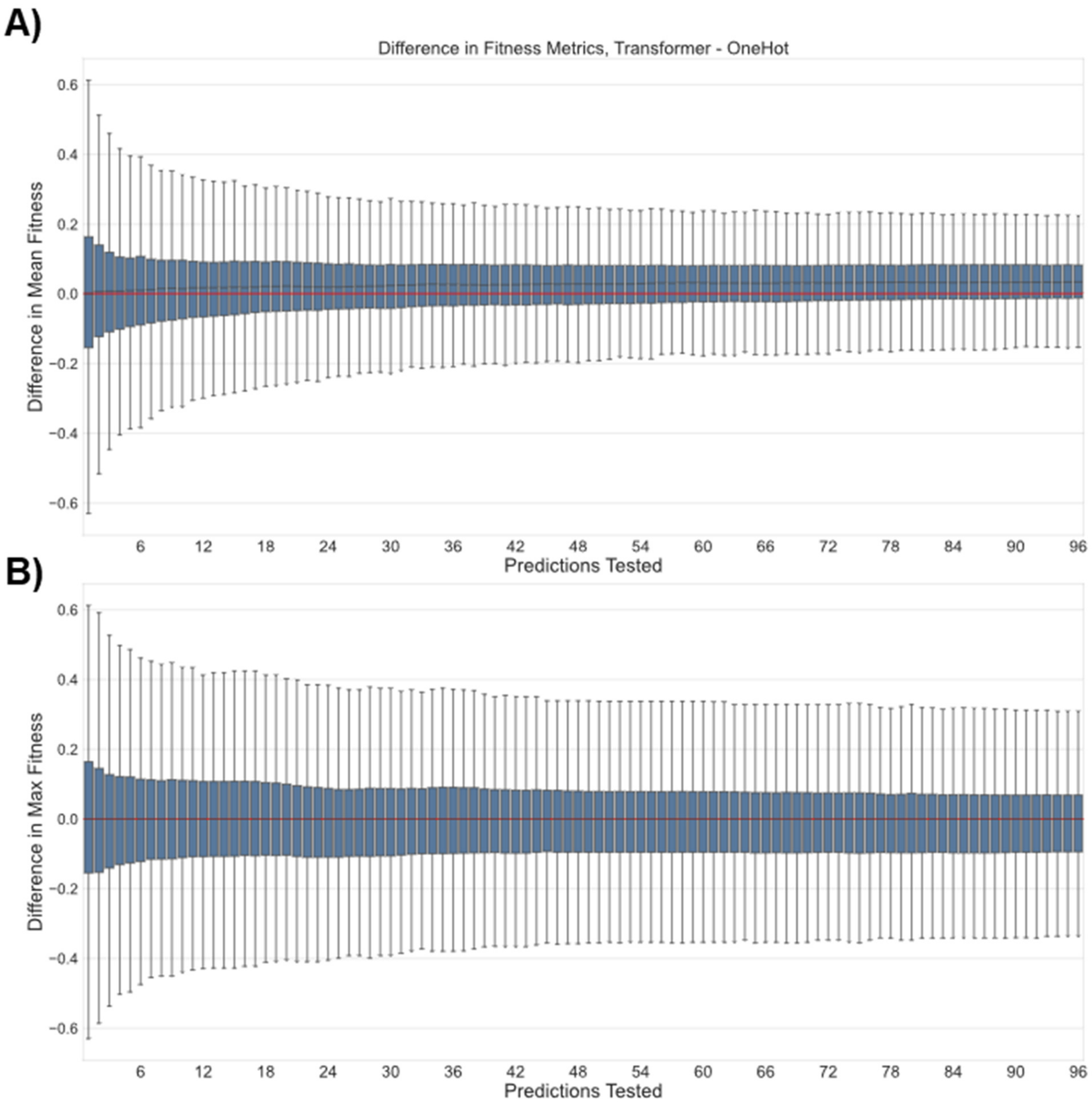

**Figure S11.** Pairwise comparison of fitness results for Transformer-derived and one-hot encodings over 2000 simulations when testing different numbers of the top-predicted variants. Extra details on the meaning and construction of this figure can be found in paragraph at the top of this section. The x-axis gives the number of top predictions tested (e.g. when x = 42, the 42 variants predicted to have the highest fitness by MLDE were evaluated for each encoding strategy) and the y-axis gives the difference in either (A) the mean or (B) the maximum fitness achieved in that top sample. All differences are reported as Transformer – one-hot. The red line serves as a reference for 0 difference. Probability mass above the red line indicates a superior result using Transformer-derived encodings.

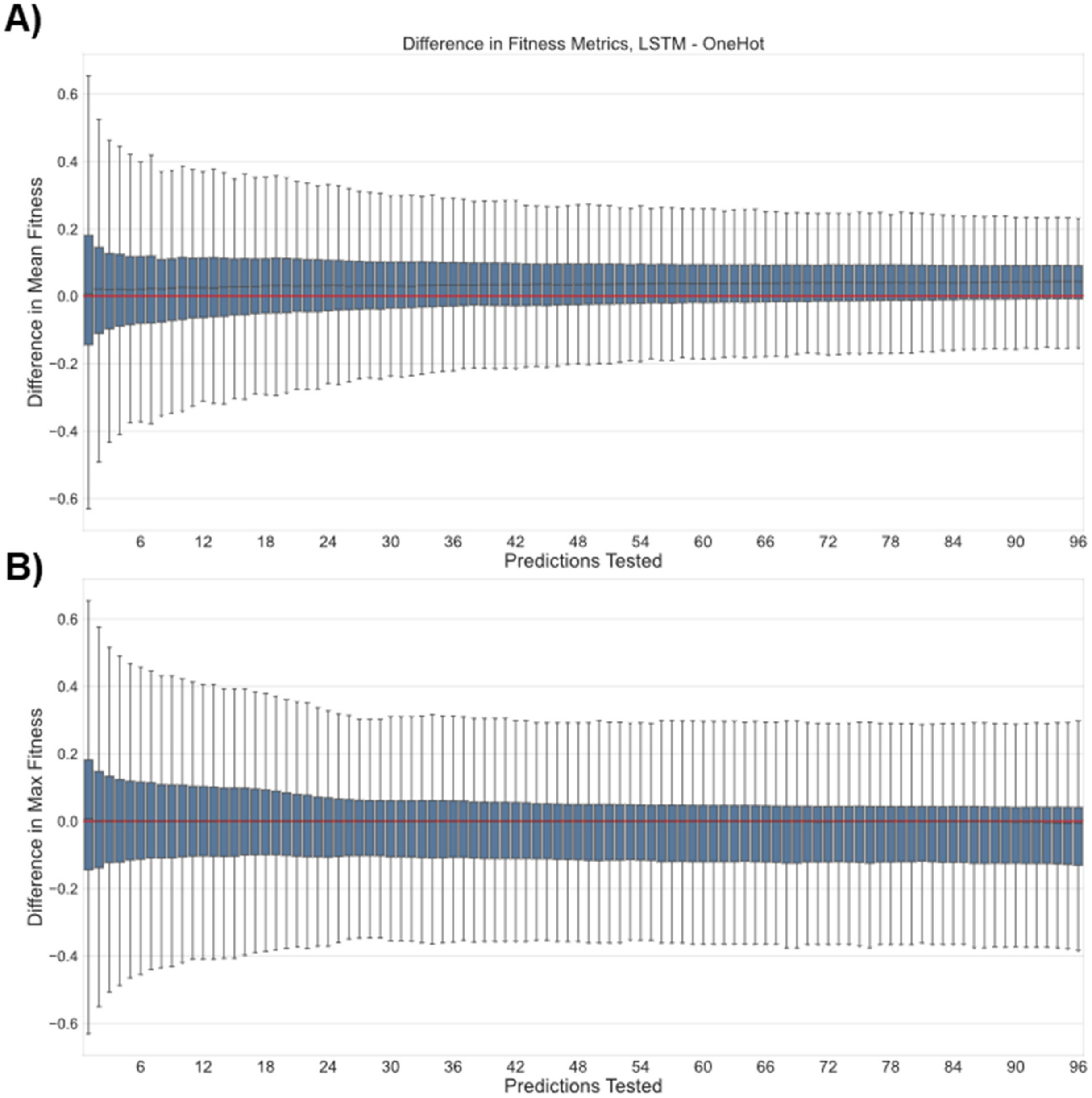

**Figure S12.** Pairwise comparison of fitness results for LSTM-derived and one-hot encodings over 2000 simulations when testing different numbers of the top-predicted variants. Extra details on the meaning and construction of this figure can be found in paragraph at the top of this section. The x-axis gives the number of top predictions tested (e.g. when x = 42, the 42 variants predicted to have the highest fitness by MLDE were evaluated for each encoding strategy) and the y-axis gives the difference in either (A) the mean or (B) the maximum fitness achieved in that top sample. All differences are reported as LSTM – one-hot. The red line serves as a reference for 0 difference. Probability mass above the red line indicates a superior result using LSTM-derived encodings.

#### Models/Training Procedures More Tailored for Combinatorial Fitness Landscapes Improve MLDE Predictive Performance

This section contains supporting information highlighting the new models incorporated into the MLDE workflow. During the training stage of MLDE, 22 different models of varied architecture are trained. To make predictions, these models are first ranked by cross-validation error, then the predictions of the top-N models are averaged and returned. Figures S13–S19 depict the frequency with which major model classes were ranked in each position across the 2000 runs of simulated MLDE for each encoding type tested in this work. The major model classes represented are: “Convolutional”, which includes the two 1D convolutional neural network architectures; “FeedForward”, which includes the two fully connected neural network architectures that have at least one hidden layer; “NoHidden”, which includes the fully connected neural network architecture with no hidden layers; “Sklearn”, which includes the 13 scikit-learn models; “XGBoost”, which includes the two XGBoost models trained with “reg:squarederror” as the learning objective; and “XGBoost-Tweedie”, which includes the two XGBoost models trained with “reg:tweedie” as the learning objective. Descriptions of the major models can be found above in *Inbuilt Models*. The x-axis in Figures S13–S19 is the rank achieved, and the y-axis is the total number of times models from each major model class were ranked in the position given by the x-axis over the 2000 runs of simulated MLDE performed for encoding comparison. Note that these results are not normalized to the number of models in the major class. Figures are shown as stacked bar plots to enable comparison between different model classes at each rank. Each figure in Figures S13–S19 shows the results for a different encoding. Table S6 details the frequency with which 1D convolutional neural networks were found in the top three models for each encoding type.

The remaining figures in this section (Figures S20–S34) make pairwise comparisons of the results of MLDE simulations using XGBoost models trained with the “reg:squarederror” objective versus XGBoost models trained with the “reg:tweedie” objective. Figure S20 compares NDCG scores achieved when using the “reg:squarederror” objective (x-axes) versus using the “reg:tweedie” objective (y-axes) for both a tree base model (Figure S20A) and a linear base model (Figure S20B) for all different encodings tested, considering the results of 2000 simulations per encoding. Figures S21–S34 show the distributions of the differences in fitness summary statistics between the results from models trained with the “reg:tweedie” objective and “reg:squarederror” objective from 2000 MLDE simulations; Figures S21–S27 are the results when a tree base model is used and Figures S28–S34 are the results when a linear base model is used; each figure in these two groups corresponds to the results from simulations with a different encoding. In each of Figures S21–S34, the x-axis describes the number of top predictions tested (e.g. when the x-axis is “10”, the boxplot above shows the distribution of differences in mean or max fitness achieved over 2000 simulations when testing the 10 unsampled variants with highest predicted fitness), while the y-axis describes the difference in fitness metrics (e.g. if the model trained with the “reg:tweedie” objective achieved a maximum fitness of “0.7” in a simulation and the model trained with the “reg:squarederror” objective achieved a maximum fitness of “0.5”, the difference would be “0.2”—the results of 2000 such differences for the 2000 simulations performed make up the boxplot above each x-value considered). In each of Figures S21–S34, subplot A shows the difference in mean fitness achieved and subplot B shows the difference in maximum fitness achieved. All differences are calculated as the result of the model trained with the “reg:tweedie” objective minus the result of the model trained with the “reg:squarederror” objective. Regardless of encoding, models with a tree base that were trained with the “reg:tweedie” objective achieved superior NDCG, mean, and max fitness. Likewise, models with a linear base that were trained with the “reg:tweedie” objective achieved a superior NDCG, but tended to achieve a worse mean and max fitness.

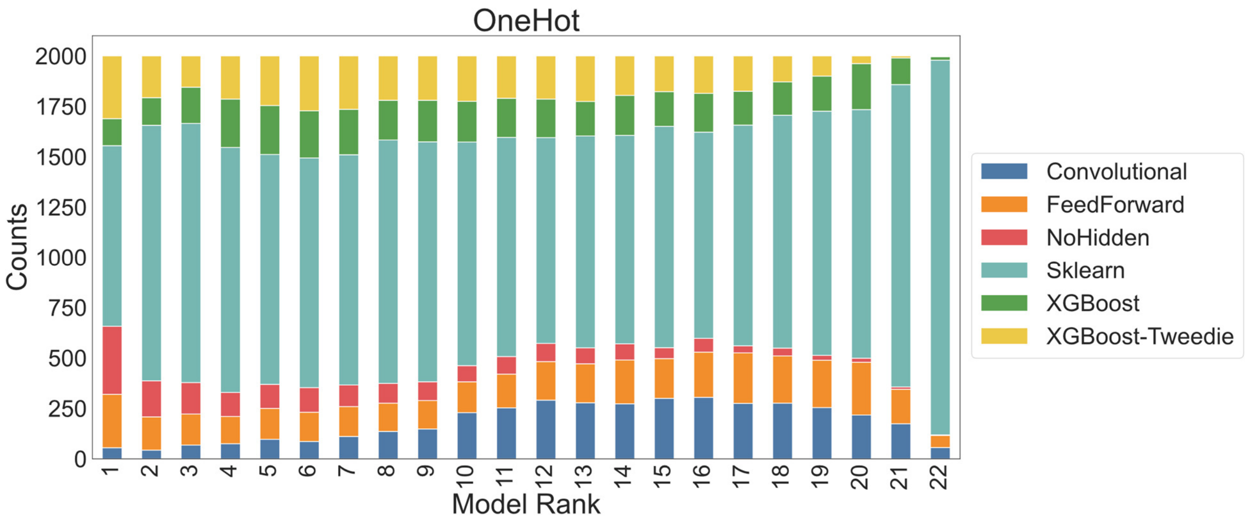

**Figure S13.** The frequency of ranking by cross-validation error of all model types over 2000 rounds of simulated MLDE using one-hot encoding. Extra details on the meaning and construction of this figure can be found in paragraphs at the top of this section.

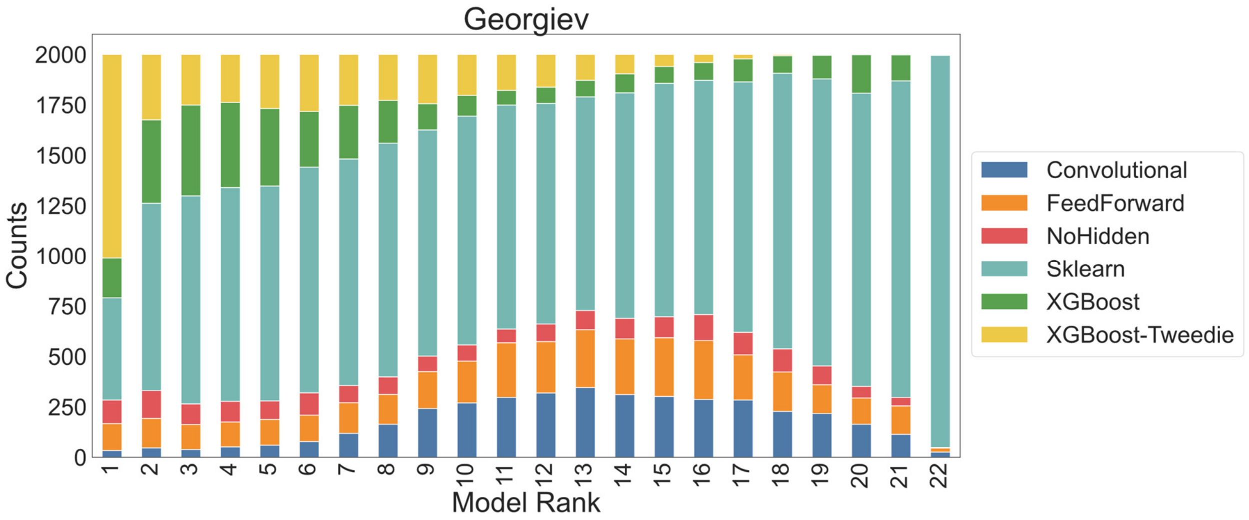

**Figure S14.** The frequency of ranking by cross-validation error of all model types over 2000 rounds of simulated MLDE using Georgiev encodings. Extra details on the meaning and construction of this figure can be found in paragraphs at the top of this section.

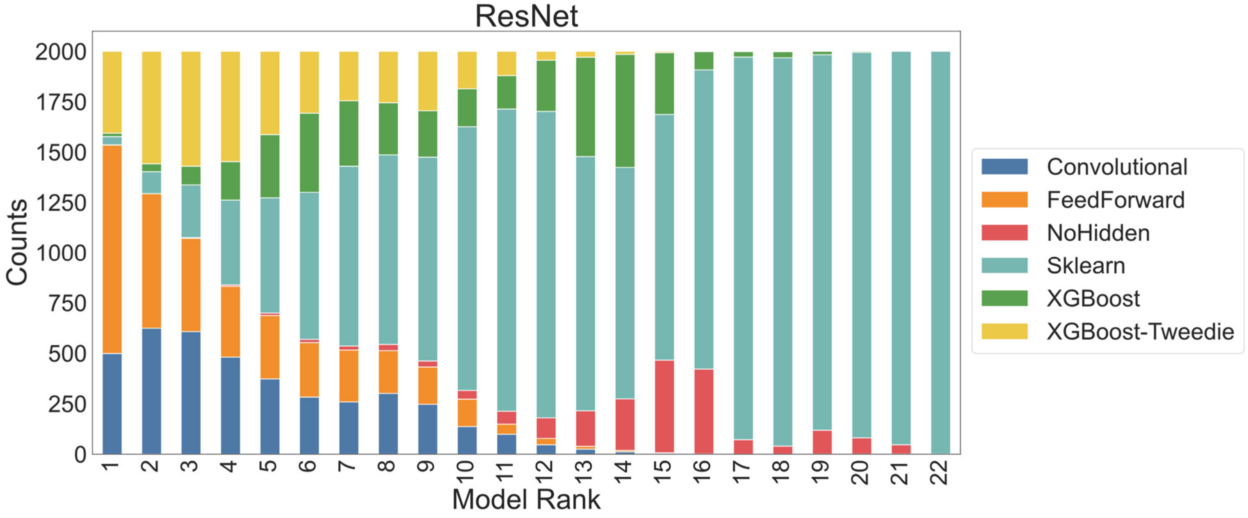

**Figure S15.** The frequency of ranking by cross-validation error of all model types over 2000 rounds of simulated MLDE using ResNet embeddings for encoding. Extra details on the meaning and construction of this figure can be found in paragraphs at the top of this section.

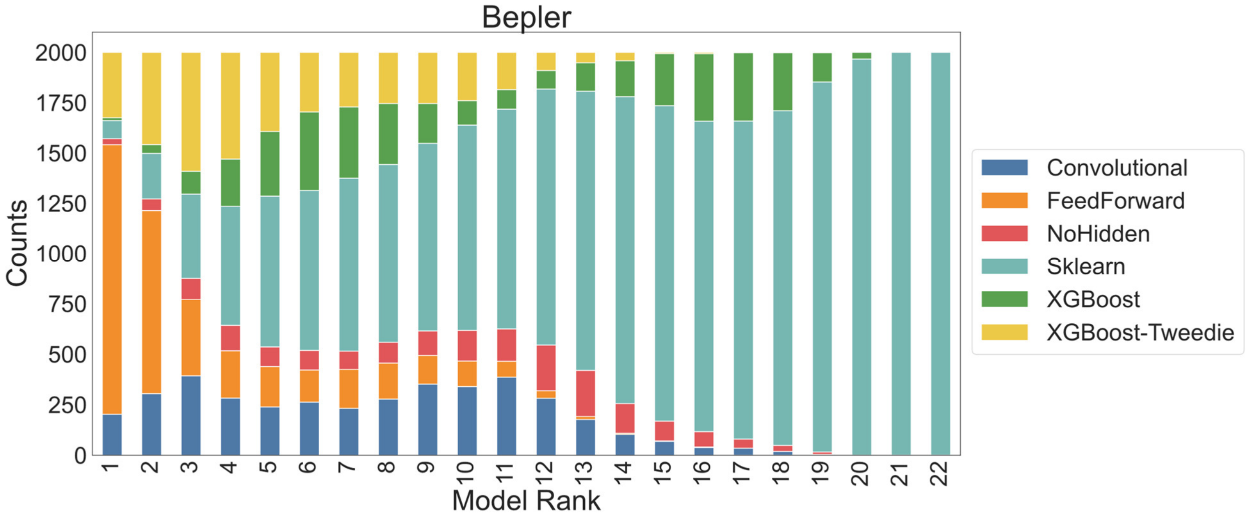

**Figure S16.** The frequency of ranking by cross-validation error of all model types over 2000 rounds of simulated MLDE using Bepler embeddings for encoding. Extra details on the meaning and construction of this figure can be found in paragraphs at the top of this section.

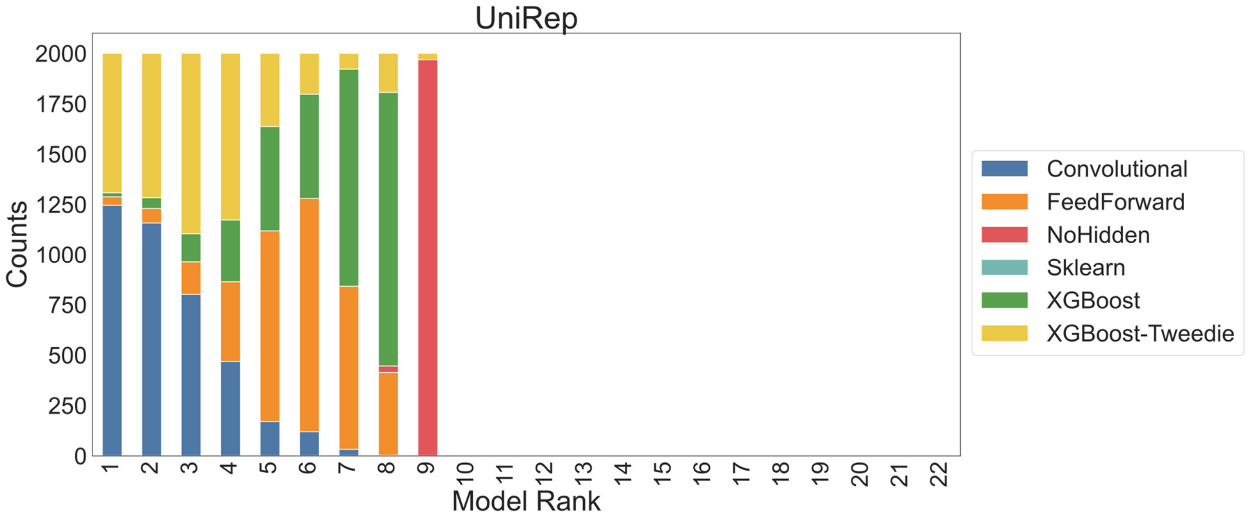

**Figure S17.** The frequency of ranking by cross-validation error of all model types over 2000 rounds of simulated MLDE using UniRep embeddings for encoding. Note that scikit-learn models were not tested for this encoding due to computational limitations, hence why the highest rank is "9". Extra details on the meaning and construction of this figure can be found in paragraphs at the top of this section.

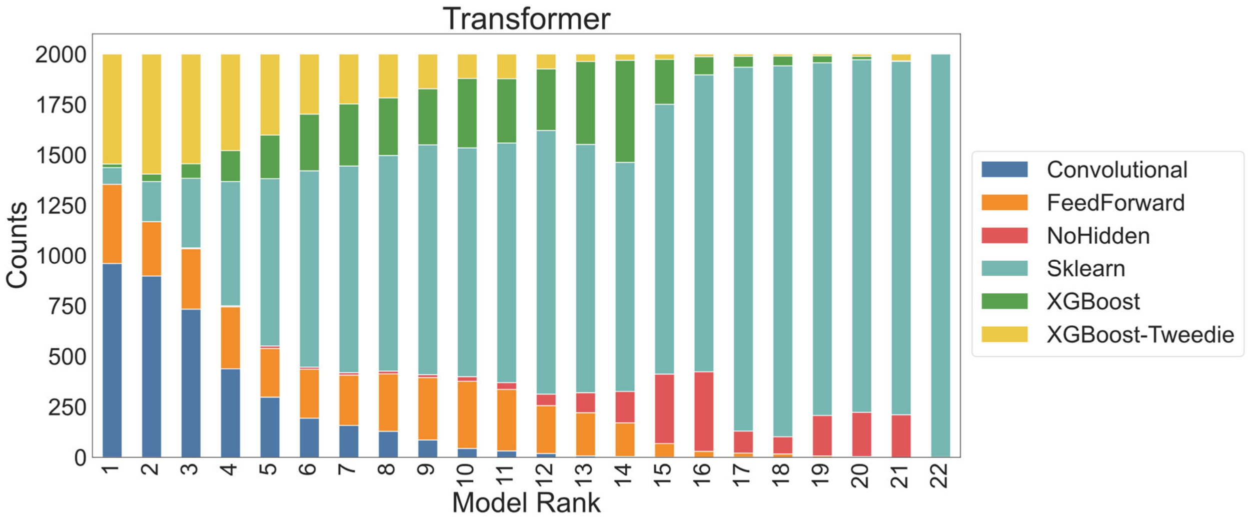

**Figure S18.** The frequency of ranking by cross-validation error of all model types over 2000 rounds of simulated MLDE using Transformer embeddings for encoding. Extra details on the meaning and construction of this figure can be found in paragraphs at the top of this section.

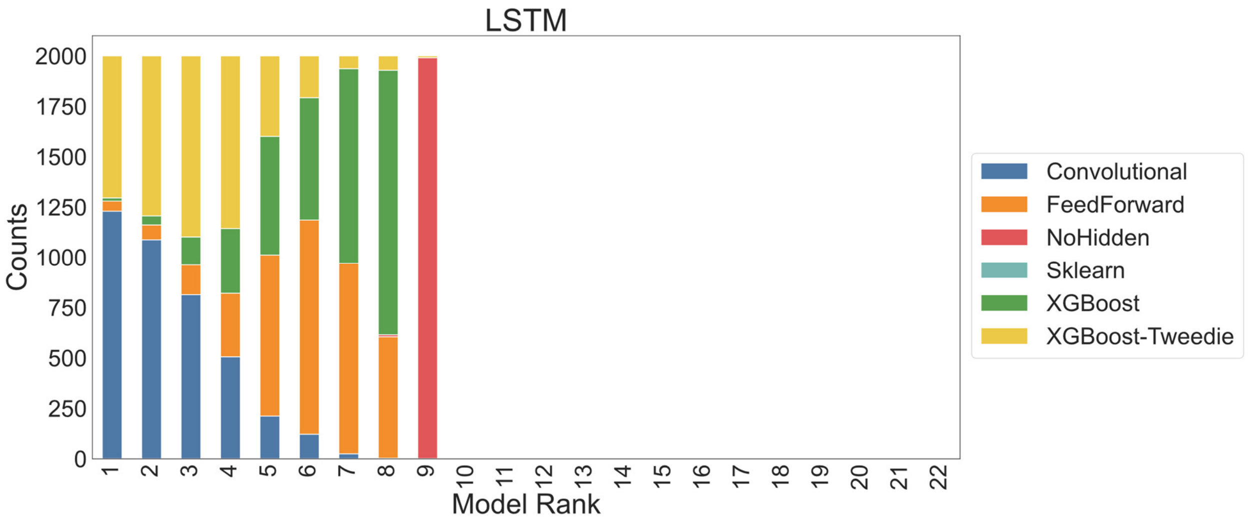

**Figure S19.** The frequency of ranking by cross-validation error of all model types over 2000 rounds of simulated MLDE using LSTM embeddings for encoding. Note that scikit-learn models were not tested for this encoding due to computational limitations, hence why the highest rank is “9”. Extra details on the meaning and construction of this figure can be found in paragraphs at the top of this section.

**Table S6.** The frequency with which at least one 1D convolutional neural network (1D CNN) appeared in the top 3 models over 2000 rounds of simulated MLDE for each encoding type. Note that the frequency with which 1D CNNs appear in the top 3 models increases with the dimensionality of the encoding. Note also that simulations using UniRep- and LSTM-derived encodings were trained only on Keras and XGBoost models while the other simulations were trained using Keras, XGBoost, and scikit-learn models.

| Encoding | Amino Acid Encoding Dimensionality | Frequency CNN in Top 3 |
|---|---|---|
| **Georgiev** | 19 | 2.00% |
| **OneHot** | 20 | 2.77% |
| **Bepler** | 100 | 15.02% |
| **ResNet** | 256 | 28.83% |
| **Transformer** | 512 | 43.15% |
| **UniRep*** | 1900 | 53.42% |
| **LSTM*** | 2048 | 52.20% |

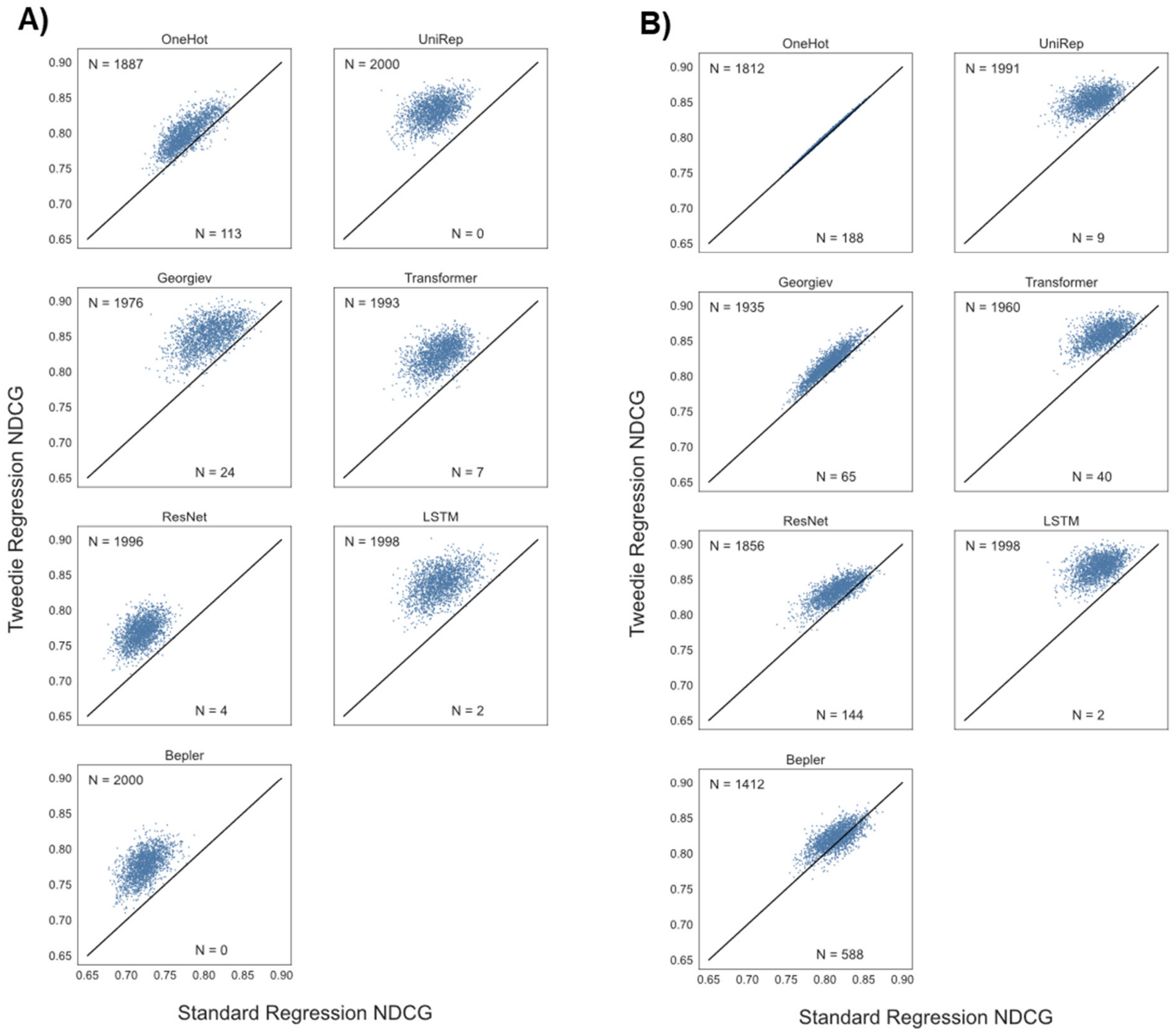

**Figure S20.** A pairwise comparison of NDCG for different XGBoost regression strategies over 2000 MLDE simulations for each possible encoding strategy. Each x-axis corresponds to the NDCG achieved using the default XGBoost regression objective (reg:squarederror) while each y-axis corresponds to the NDCG achieved using the Tweedie regression objective (reg:tweedie). Each subplot corresponds to the results when using a different encoding. Subplots under (A) are the results when using a base tree model, while subplots under (B) are the results when using a base linear model. Extra details on the meaning and construction of this figure can be found in paragraphs at the top of this section.

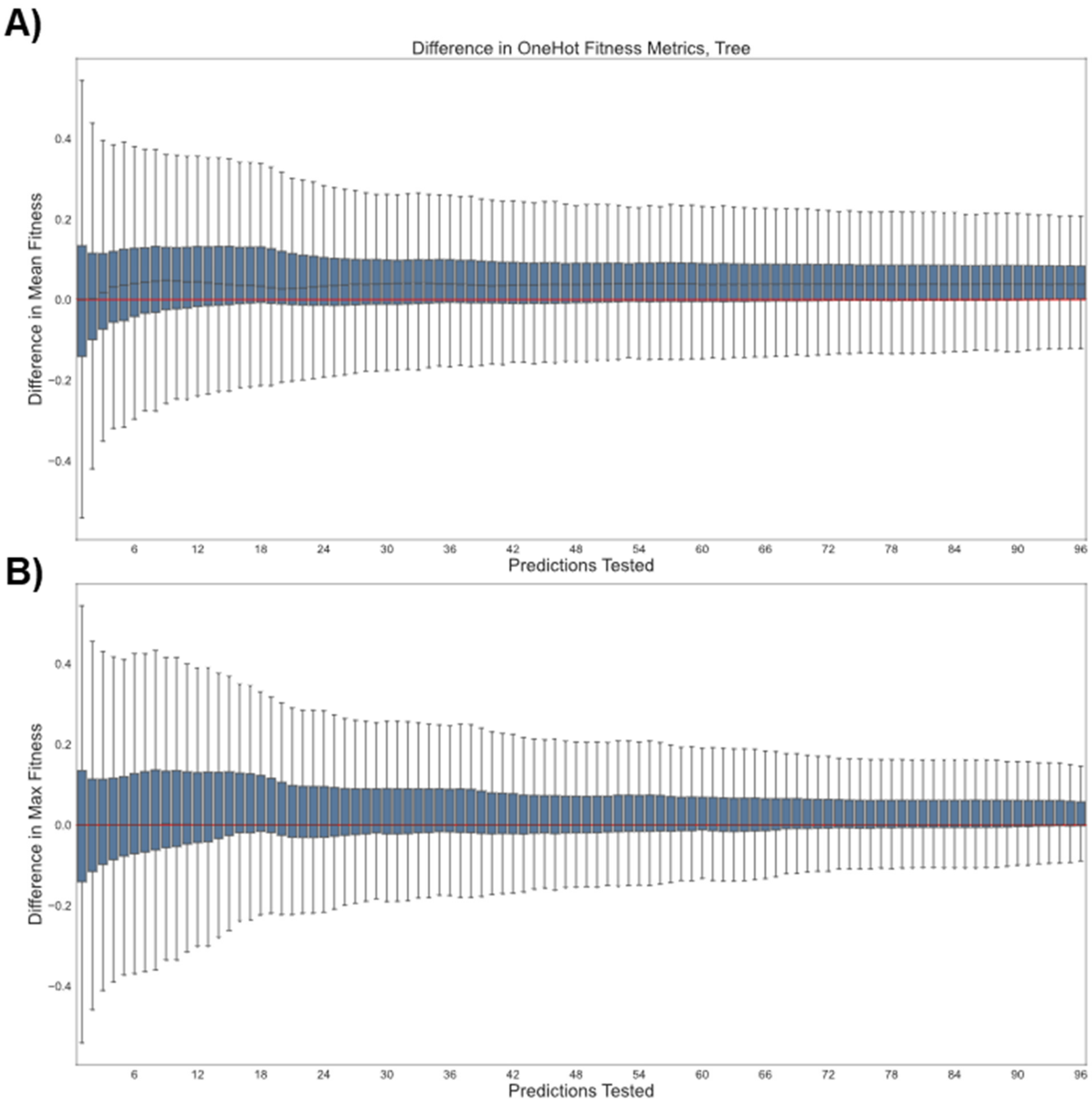

**Figure S21.** Pairwise comparison of fitness results for XGBoost with the Tweedie regression objective (reg:tweedie) and the default regression objective (reg:squarederror) over 2000 simulations using one-hot encoding and a base tree model. The x-axis gives the number of top predictions tested (e.g. when x = 42, the 42 variants predicted to have the highest fitness by MLDE were evaluated for each regression strategy) and the y-axis gives the difference in either (A) the mean or (B) the maximum fitness achieved in that top sample. All differences are reported as reg:tweedie result – reg:squarederror result. The red line serves as a reference for 0 difference. Probability mass above the red line indicates a superior result using Tweedie regression. Extra details on the meaning and construction of this figure can be found in paragraphs at the top of this section.

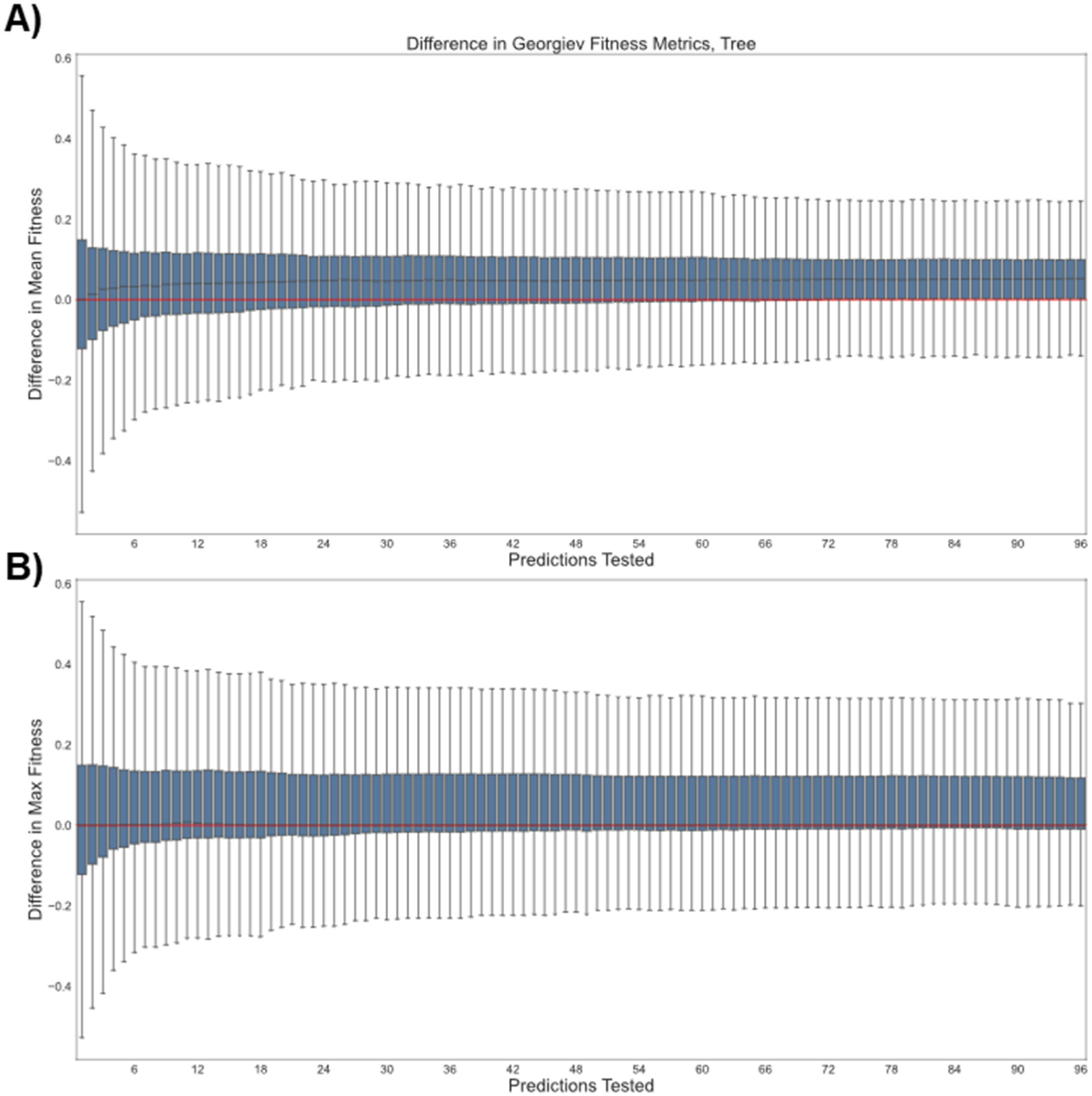

**Figure S22.** Pairwise comparison of fitness results for XGBoost with the Tweedie regression objective (reg:tweedie) and the default regression objective (reg:squarederror) over 2000 simulations using Georgiev encodings and a base tree model. The x-axis gives the number of top predictions tested (e.g. when x = 42, the 42 variants predicted to have the highest fitness by MLDE were evaluated for each regression strategy) and the y-axis gives the difference in either (A) the mean or (B) the maximum fitness achieved in that top sample. All differences are reported as reg:tweedie result – reg:squarederror result. The red line serves as a reference for 0 difference. Probability mass above the red line indicates a superior result using Tweedie regression. Extra details on the meaning and construction of this figure can be found in paragraphs at the top of this section.

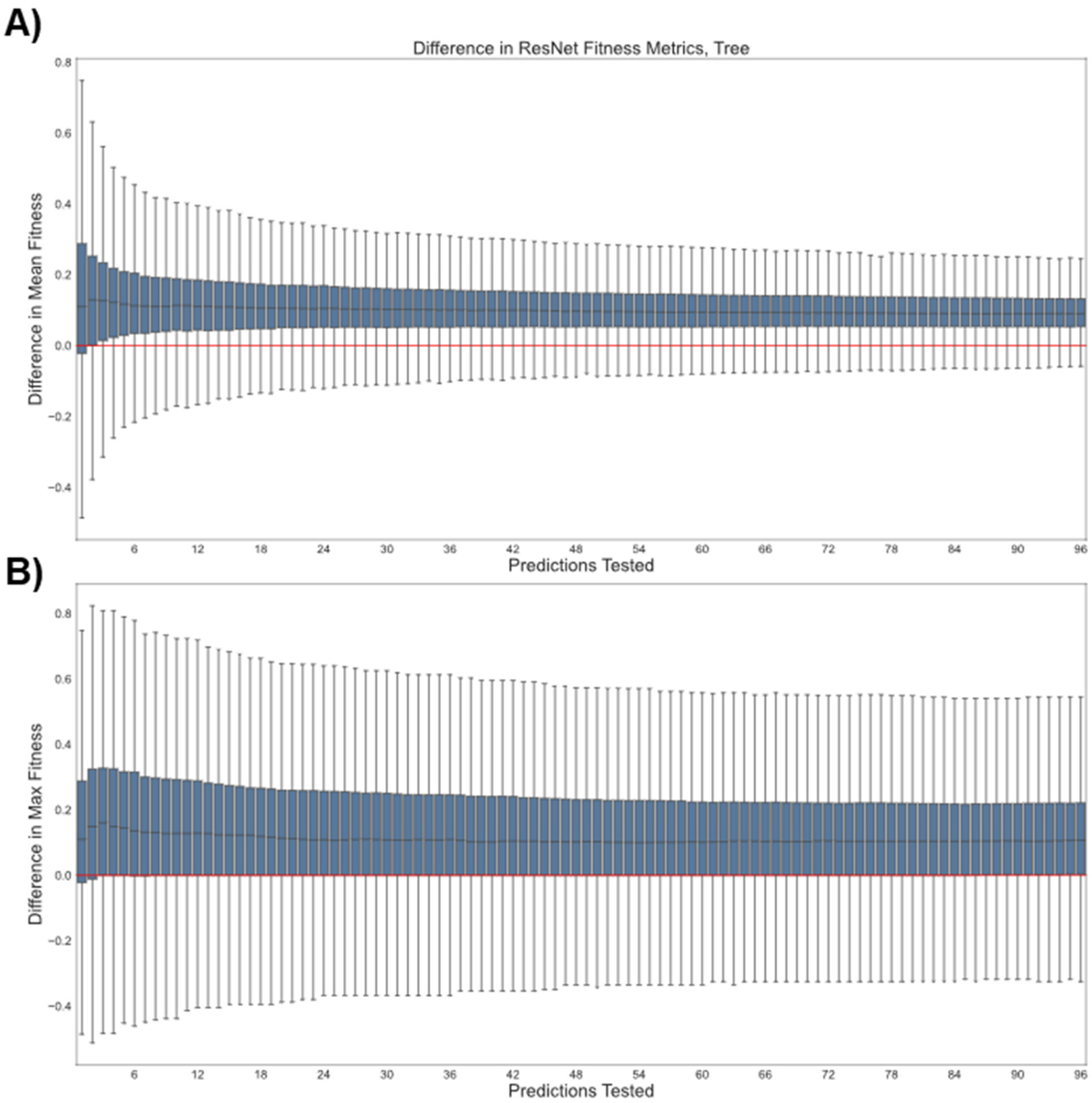

**Figure S23.** Pairwise comparison of fitness results for XGBoost with the Tweedie regression objective (reg:tweedie) and the default regression objective (reg:squarederror) over 2000 simulations using ResNet-derived encodings and a base tree model. The x-axis gives the number of top predictions tested (e.g. when x = 42, the 42 variants predicted to have the highest fitness by MLDE were evaluated for each regression strategy) and the y-axis gives the difference in either (A) the mean or (B) the maximum fitness achieved in that top sample. All differences are reported as reg:tweedie result – reg:squarederror result. The red line serves as a reference for 0 difference. Probability mass above the red line indicates a superior result using Tweedie regression. Extra details on the meaning and construction of this figure can be found in paragraphs at the top of this section.

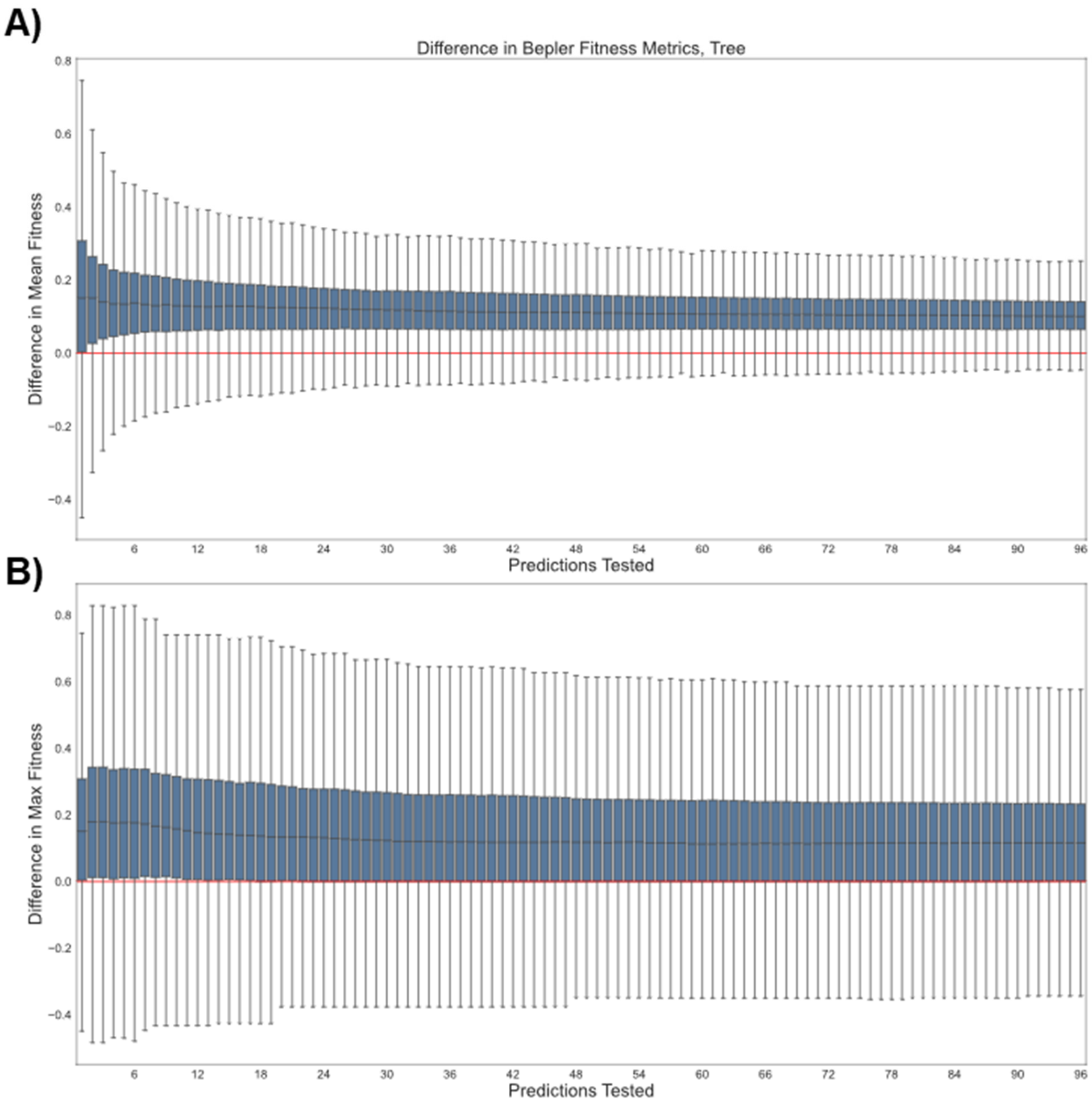

**Figure S24.** Pairwise comparison of fitness results for XGBoost with the Tweedie regression objective (reg:tweedie) and the default regression objective (reg:squarederror) over 2000 simulations using Bepler-derived encodings and a base tree model. The x-axis gives the number of top predictions tested (e.g. when x = 42, the 42 variants predicted to have the highest fitness by MLDE were evaluated for each regression strategy) and the y-axis gives the difference in either (A) the mean or (B) the maximum fitness achieved in that top sample. All differences are reported as reg:tweedie result – reg:squarederror result. The red line serves as a reference for 0 difference. Probability mass above the red line indicates a superior result using Tweedie regression. Extra details on the meaning and construction of this figure can be found in paragraphs at the top of this section.

**A)**

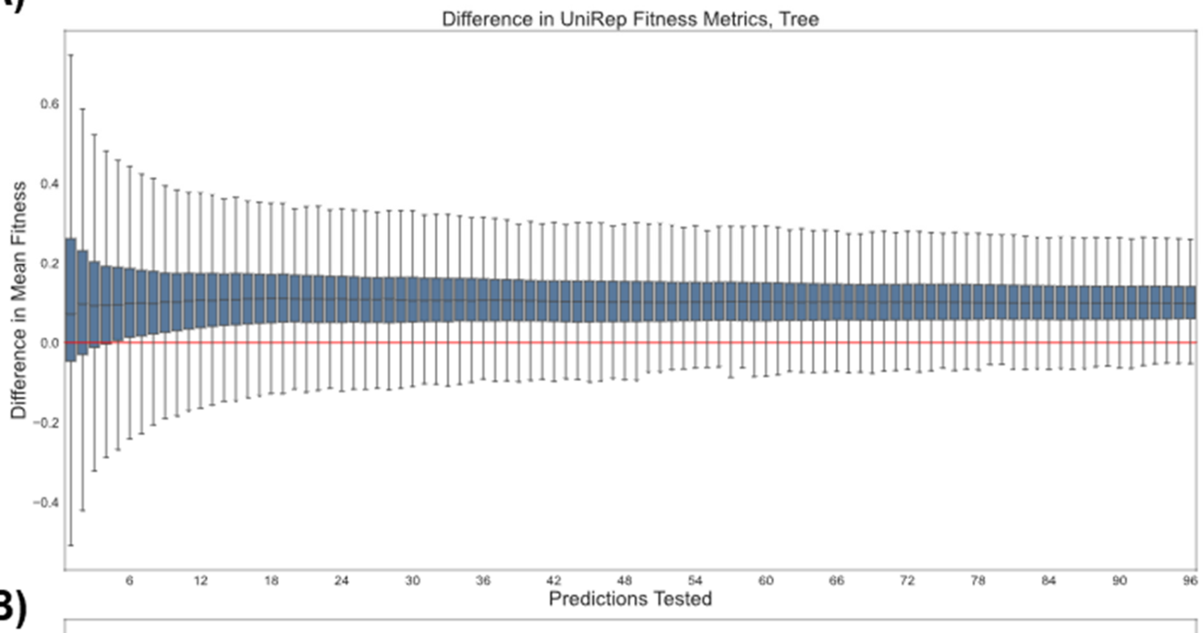

**B)**

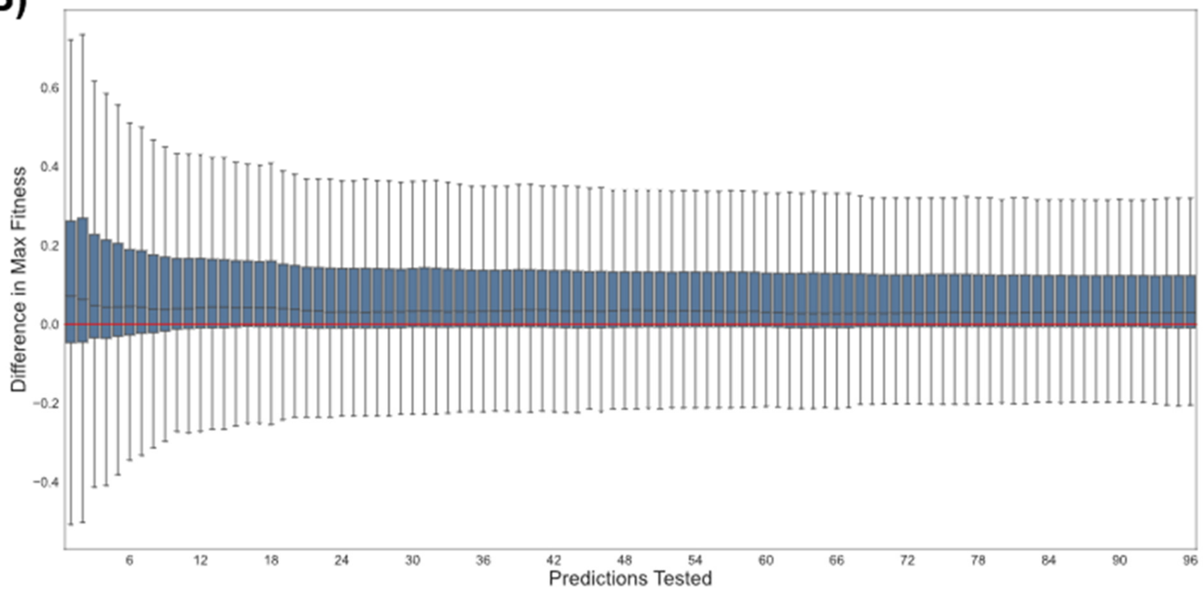

**Figure S25.** Pairwise comparison of fitness results for XGBoost with the Tweedie regression objective (reg:tweedie) and the default regression objective (reg:squarederror) over 2000 simulations using UniRep-derived encodings and a base tree model. The x-axis gives the number of top predictions tested (e.g. when x = 42, the 42 variants predicted to have the highest fitness by MLDE were evaluated for each regression strategy) and the y-axis gives the difference in either (A) the mean or (B) the maximum fitness achieved in that top sample. All differences are reported as reg:tweedie result – reg:squarederror result. The red line serves as a reference for 0 difference. Probability mass above the red line indicates a superior result using Tweedie regression. Extra details on the meaning and construction of this figure can be found in paragraphs at the top of this section.

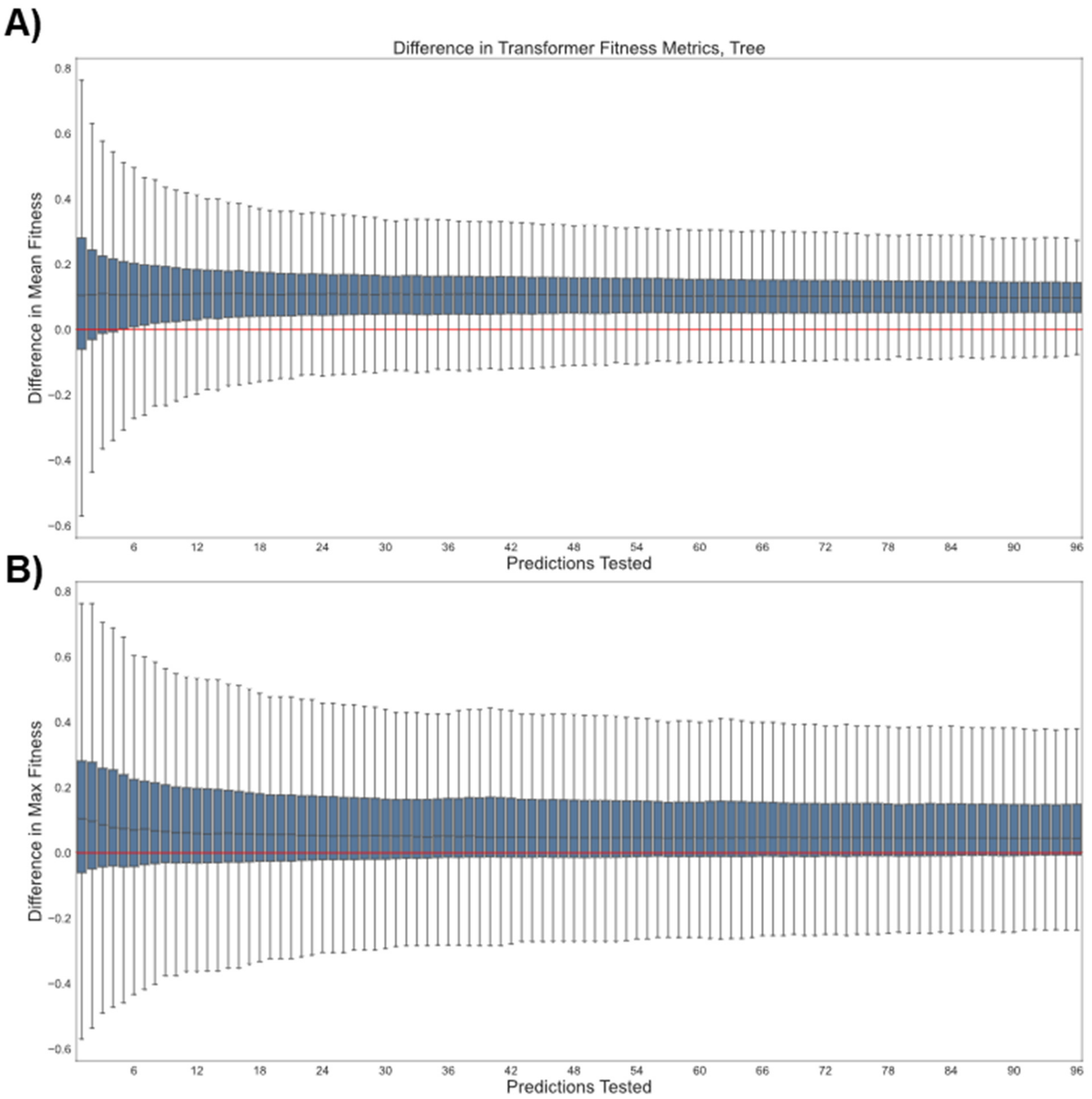

**Figure S26.** Pairwise comparison of fitness results for XGBoost with the Tweedie regression objective (reg:tweedie) and the default regression objective (reg:squarederror) over 2000 simulations using Transformer-derived encodings and a base tree model. The x-axis gives the number of top predictions tested (e.g. when x = 42, the 42 variants predicted to have the highest fitness by MLDE were evaluated for each regression strategy) and the y-axis gives the difference in either (A) the mean or (B) the maximum fitness achieved in that top sample. All differences are reported as reg:tweedie result – reg:squarederror result. The red line serves as a reference for 0 difference. Probability mass above the red line indicates a superior result using Tweedie regression. Extra details on the meaning and construction of this figure can be found in paragraphs at the top of this section.

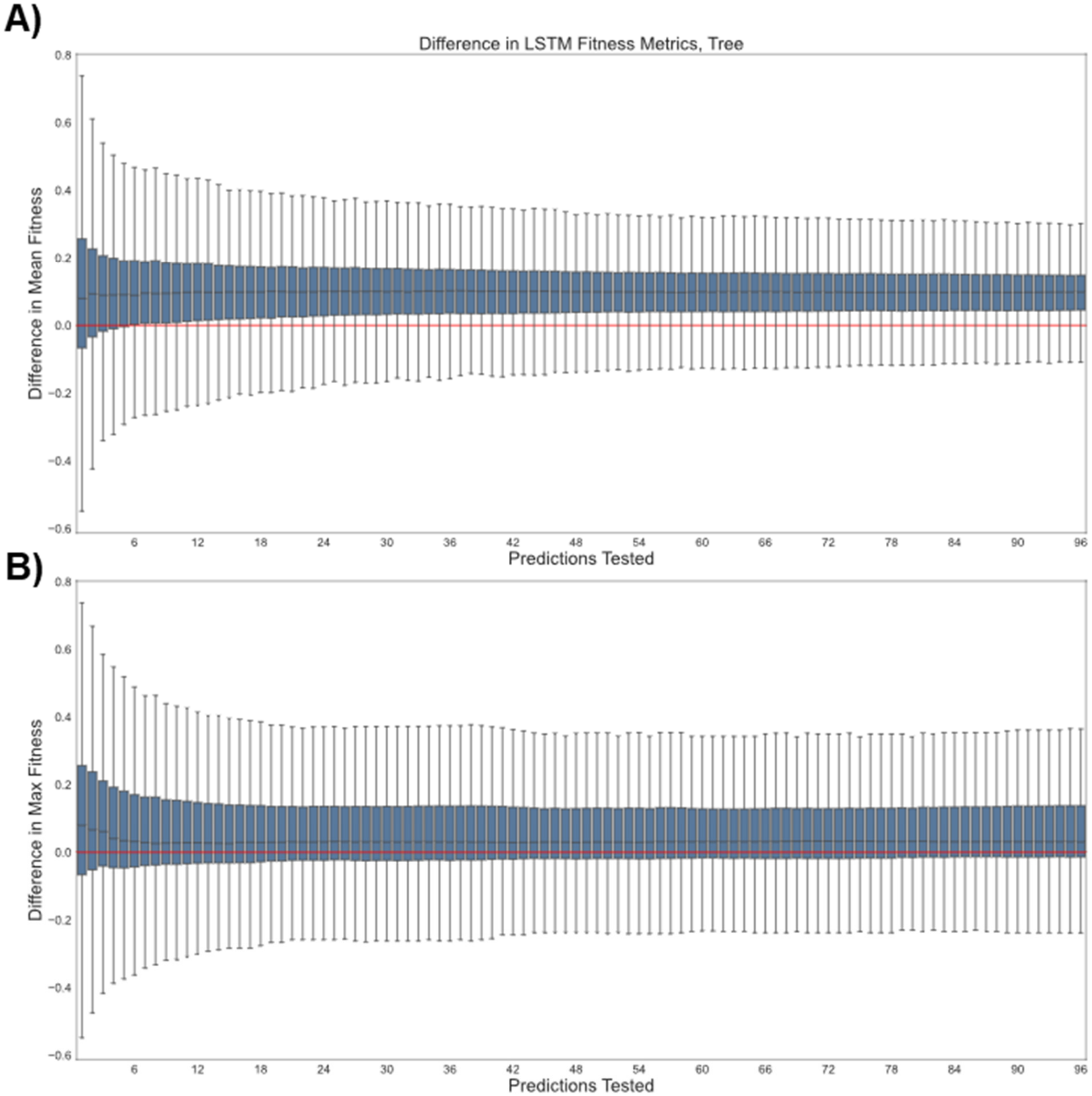

**Figure S27.** Pairwise comparison of fitness results for XGBoost with the Tweedie regression objective (reg:tweedie) and the default regression objective (reg:squarederror) over 2000 simulations using LSTM-derived encodings and a base tree model. The x-axis gives the number of top predictions tested (e.g. when x = 42, the 42 variants predicted to have the highest fitness by MLDE were evaluated for each regression strategy) and the y-axis gives the difference in either (A) the mean or (B) the maximum fitness achieved in that top sample. All differences are reported as reg:tweedie result – reg:squarederror result. The red line serves as a reference for 0 difference. Probability mass above the red line indicates a superior result using Tweedie regression. Extra details on the meaning and construction of this figure can be found in paragraphs at the top of this section.

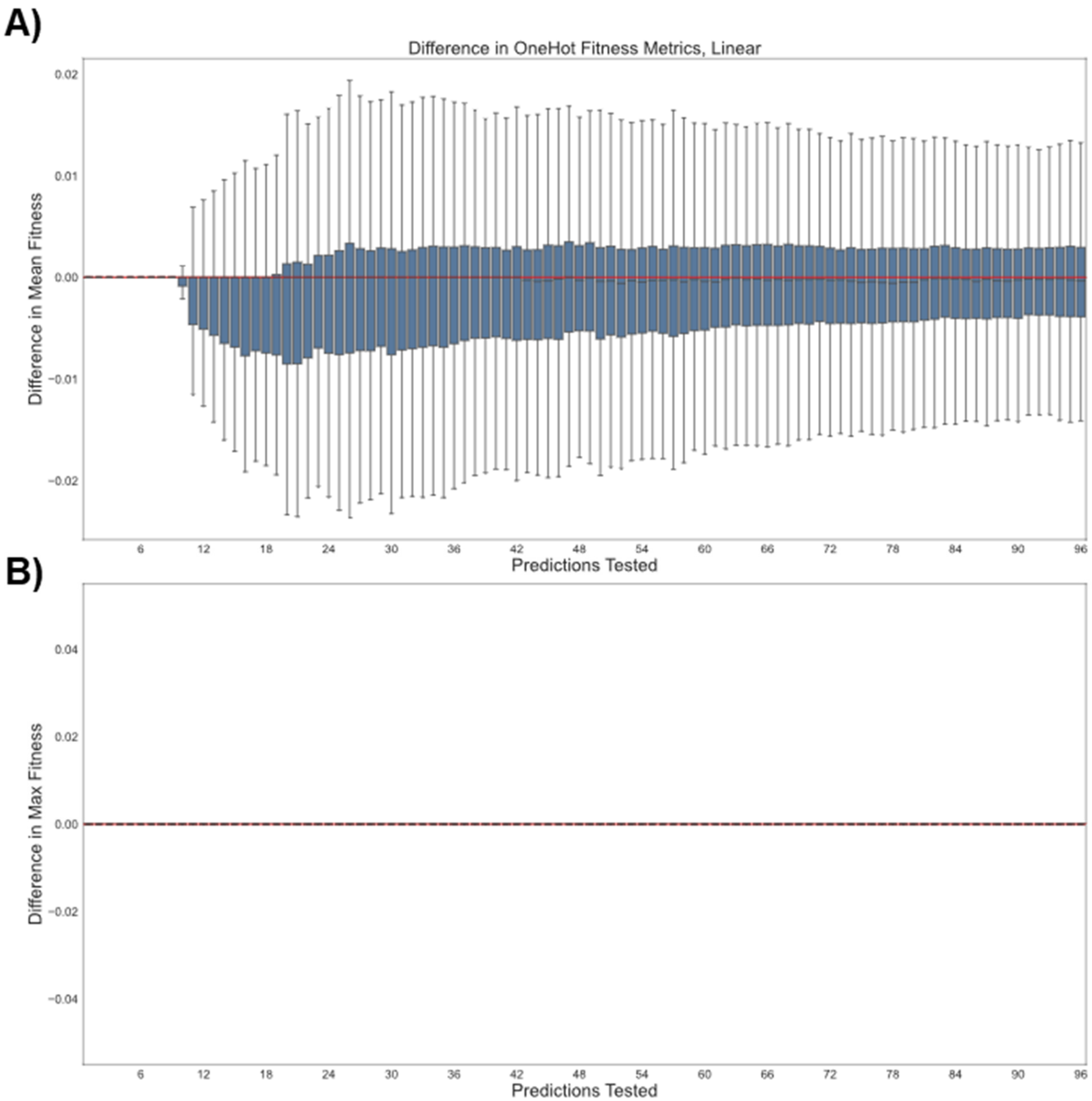

**Figure S28.** Pairwise comparison of fitness results for XGBoost with the Tweedie regression objective (reg:tweedie) and the default regression objective (reg:squarederror) over 2000 simulations using one-hot encoding and a base linear model. The x-axis gives the number of top predictions tested (e.g. when x = 42, the 42 variants predicted to have the highest fitness by MLDE were evaluated for each regression strategy) and the y-axis gives the difference in either (A) the mean or (B) the maximum fitness achieved in that top sample. All differences are reported as reg:tweedie result – reg:squarederror result. The red line serves as a reference for 0 difference. Probability mass above the red line indicates a superior result using Tweedie regression. **Note that all simulations achieved the same maximum fitness regardless of regression strategy, hence the lack of distributions.**

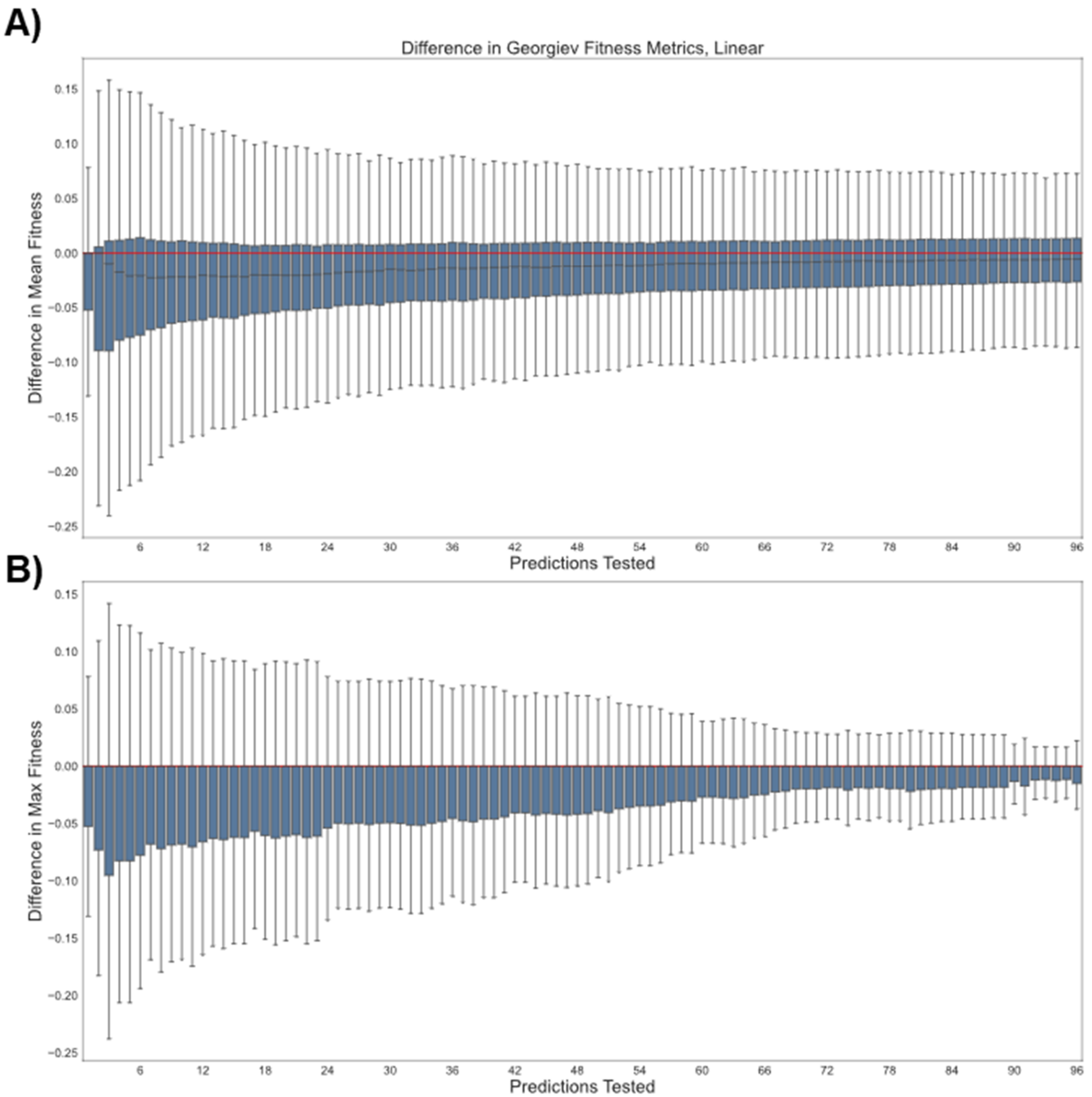

**Figure S29.** Pairwise comparison of fitness results for XGBoost with the Tweedie regression objective (reg:tweedie) and the default regression objective (reg:squarederror) over 2000 simulations using Georgiev encodings and a base linear model. The x-axis gives the number of top predictions tested (e.g. when x = 42, the 42 variants predicted to have the highest fitness by MLDE were evaluated for each regression strategy) and the y-axis gives the difference in either (A) the mean or (B) the maximum fitness achieved in that top sample. All differences are reported as reg:tweedie result – reg:squarederror result. The red line serves as a reference for 0 difference. Probability mass above the red line indicates a superior result using Tweedie regression. Extra details on the meaning and construction of this figure can be found in paragraphs at the top of this section.

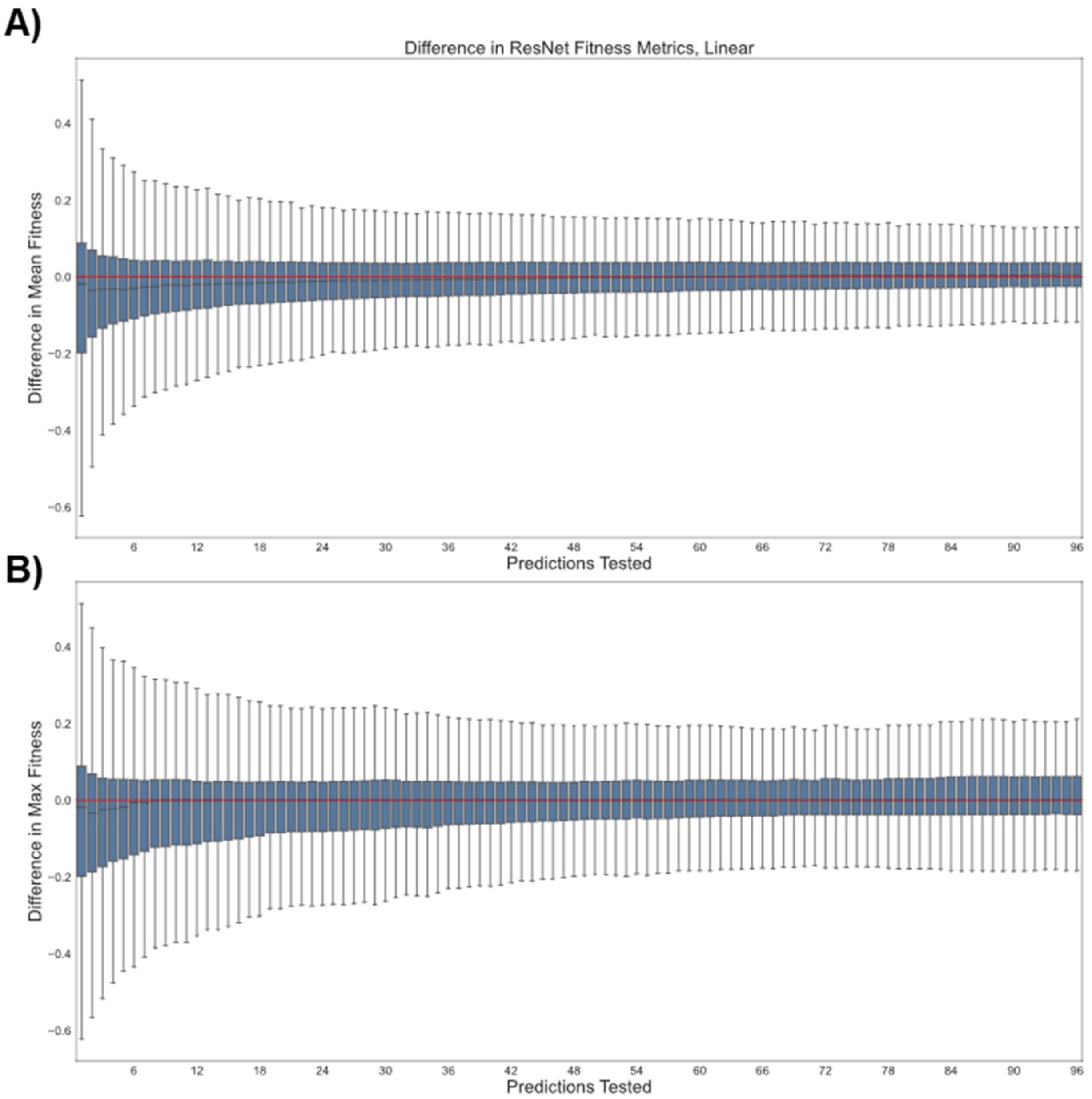

**Figure S30.** Pairwise comparison of fitness results for XGBoost with the Tweedie regression objective (reg:tweedie) and the default regression objective (reg:squarederror) over 2000 simulations using ResNet-derived encodings and a base linear model. The x-axis gives the number of top predictions tested (e.g. when x = 42, the 42 variants predicted to have the highest fitness by MLDE were evaluated for each regression strategy) and the y-axis gives the difference in either (A) the mean or (B) the maximum fitness achieved in that top sample. All differences are reported as reg:tweedie result – reg:squarederror result. The red line serves as a reference for 0 difference. Probability mass above the red line indicates a superior result using Tweedie regression. Extra details on the meaning and construction of this figure can be found in paragraphs at the top of this section.

A)

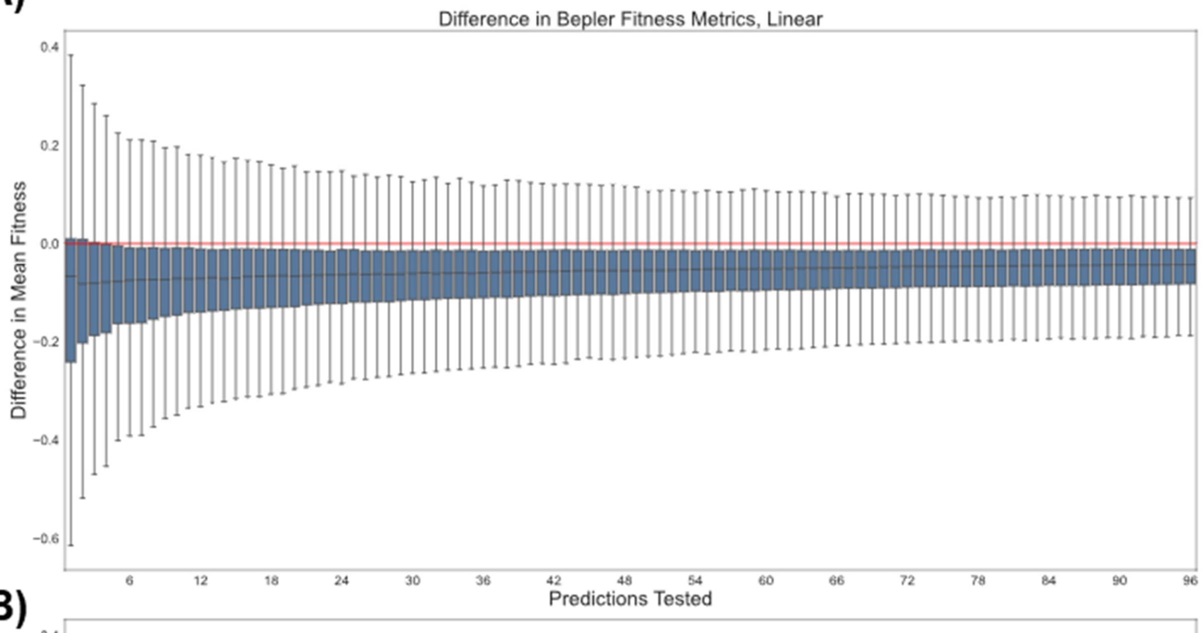

B)

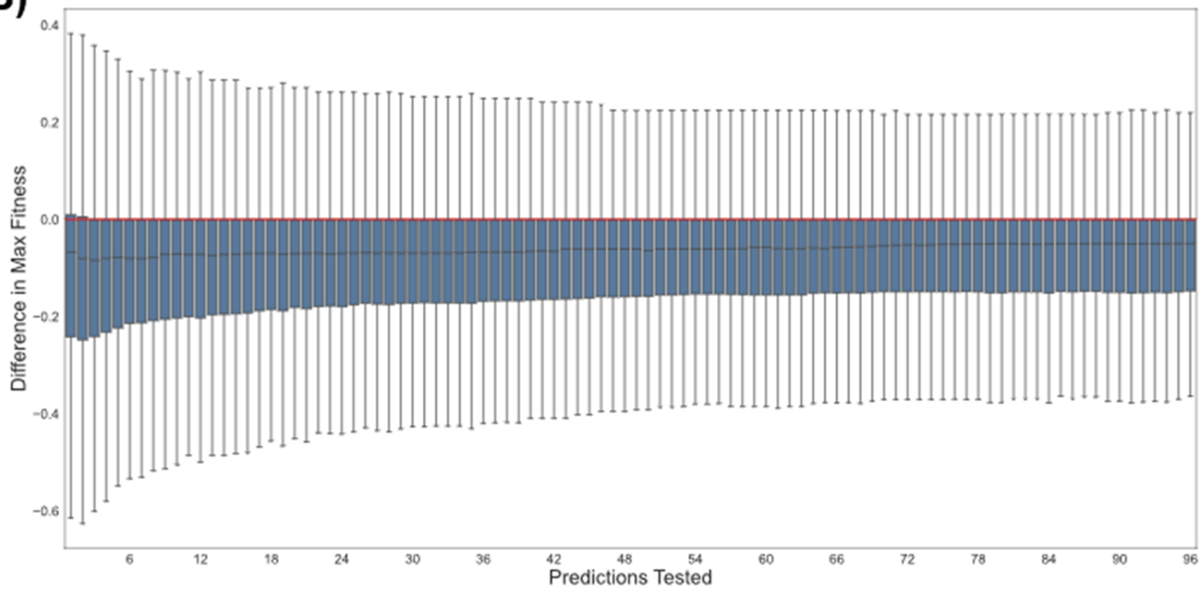

**Figure S31.** Pairwise comparison of fitness results for XGBoost with the Tweedie regression objective (reg:tweedie) and the default regression objective (reg:squarederror) over 2000 simulations using Bepler-derived encodings and a base linear model. The x-axis gives the number of top predictions tested (e.g. when x = 42, the 42 variants predicted to have the highest fitness by MLDE were evaluated for each regression strategy) and the y-axis gives the difference in either (A) the mean or (B) the maximum fitness achieved in that top sample. All differences are reported as reg:tweedie result – reg:squarederror result. The red line serves as a reference for 0 difference. Probability mass above the red line indicates a superior result using Tweedie regression. Extra details on the meaning and construction of this figure can be found in paragraphs at the top of this section.

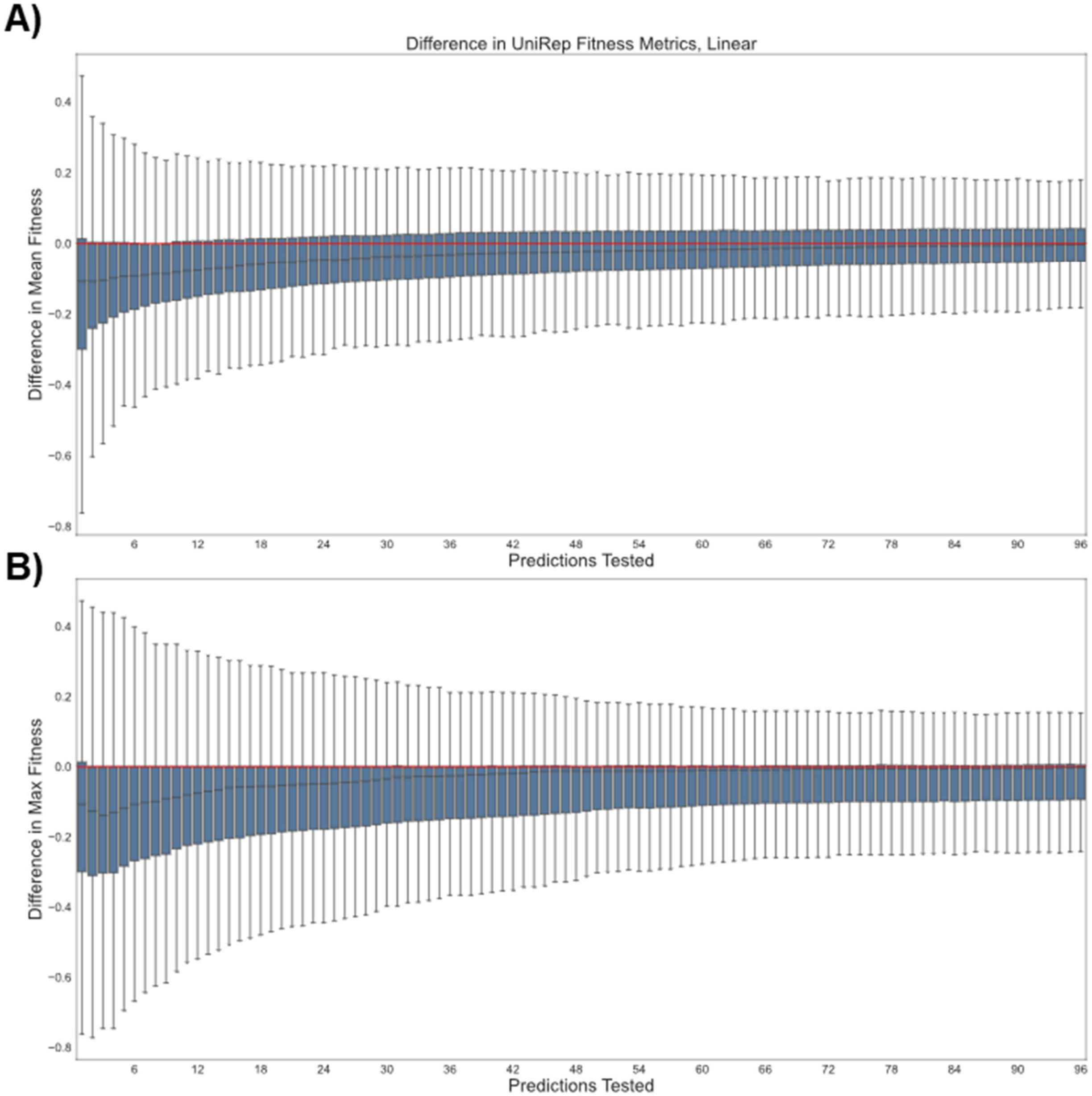

**Figure S32.** Pairwise comparison of fitness results for XGBoost with the Tweedie regression objective (reg:tweedie) and the default regression objective (reg:squarederror) over 2000 simulations using UniRep-derived encodings and a base linear model. The x-axis gives the number of top predictions tested (e.g. when x = 42, the 42 variants predicted to have the highest fitness by MLDE were evaluated for each regression strategy) and the y-axis gives the difference in either (A) the mean or (B) the maximum fitness achieved in that top sample. All differences are reported as reg:tweedie result – reg:squarederror result. The red line serves as a reference for 0 difference. Probability mass above the red line indicates a superior result using Tweedie regression. Extra details on the meaning and construction of this figure can be found in paragraphs at the top of this section.

A)

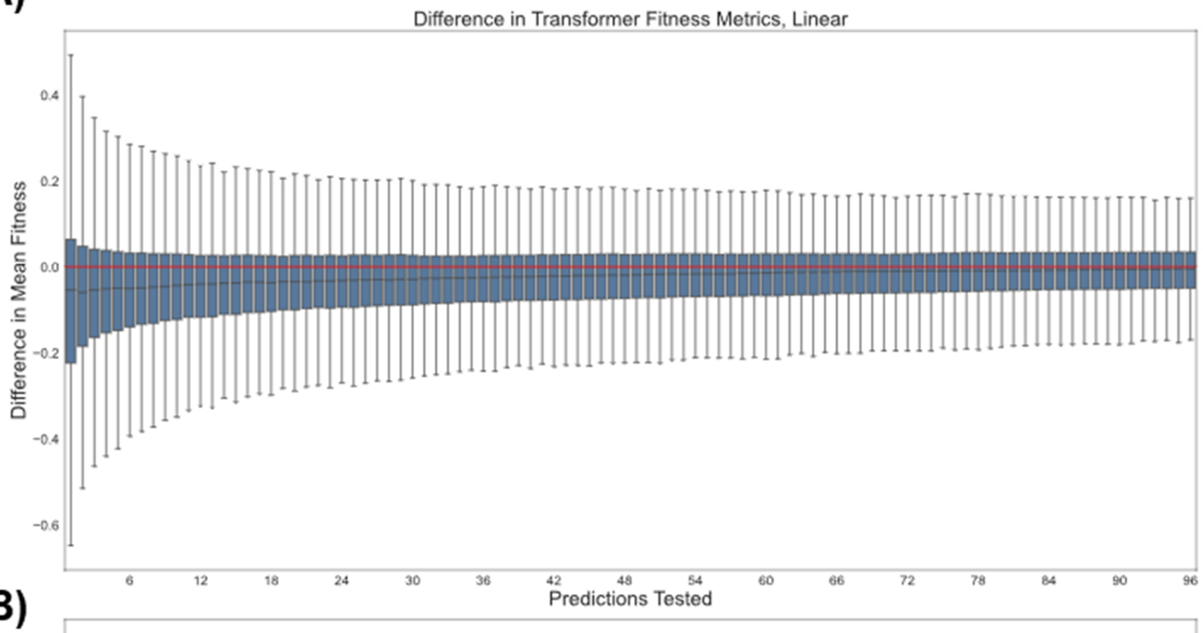

B)

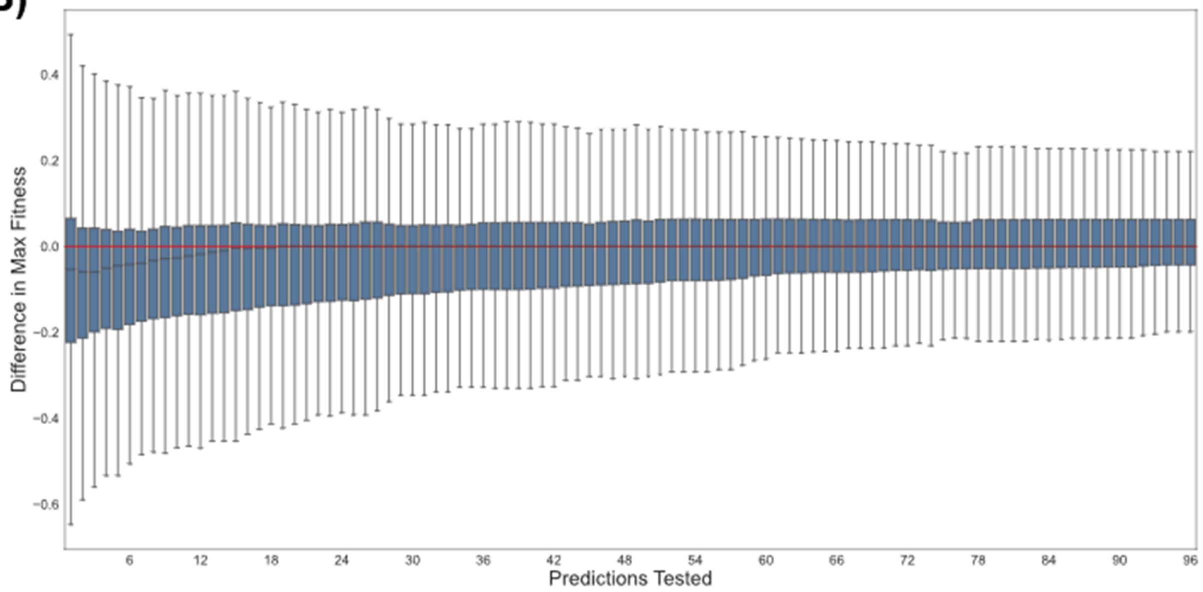

**Figure S33.** Pairwise comparison of fitness results for XGBoost with the Tweedie regression objective (reg:tweedie) and the default regression objective (reg:squarederror) over 2000 simulations using Transformer-derived encodings and a base linear model. The x-axis gives the number of top predictions tested (e.g. when x = 42, the 42 variants predicted to have the highest fitness by MLDE were evaluated for each regression strategy) and the y-axis gives the difference in either (A) the mean or (B) the maximum fitness achieved in that top sample. All differences are reported as reg:tweedie result – reg:squarederror result. The red line serves as a reference for 0 difference. Probability mass above the red line indicates a superior result using Tweedie regression. Extra details on the meaning and construction of this figure can be found in paragraphs at the top of this section.

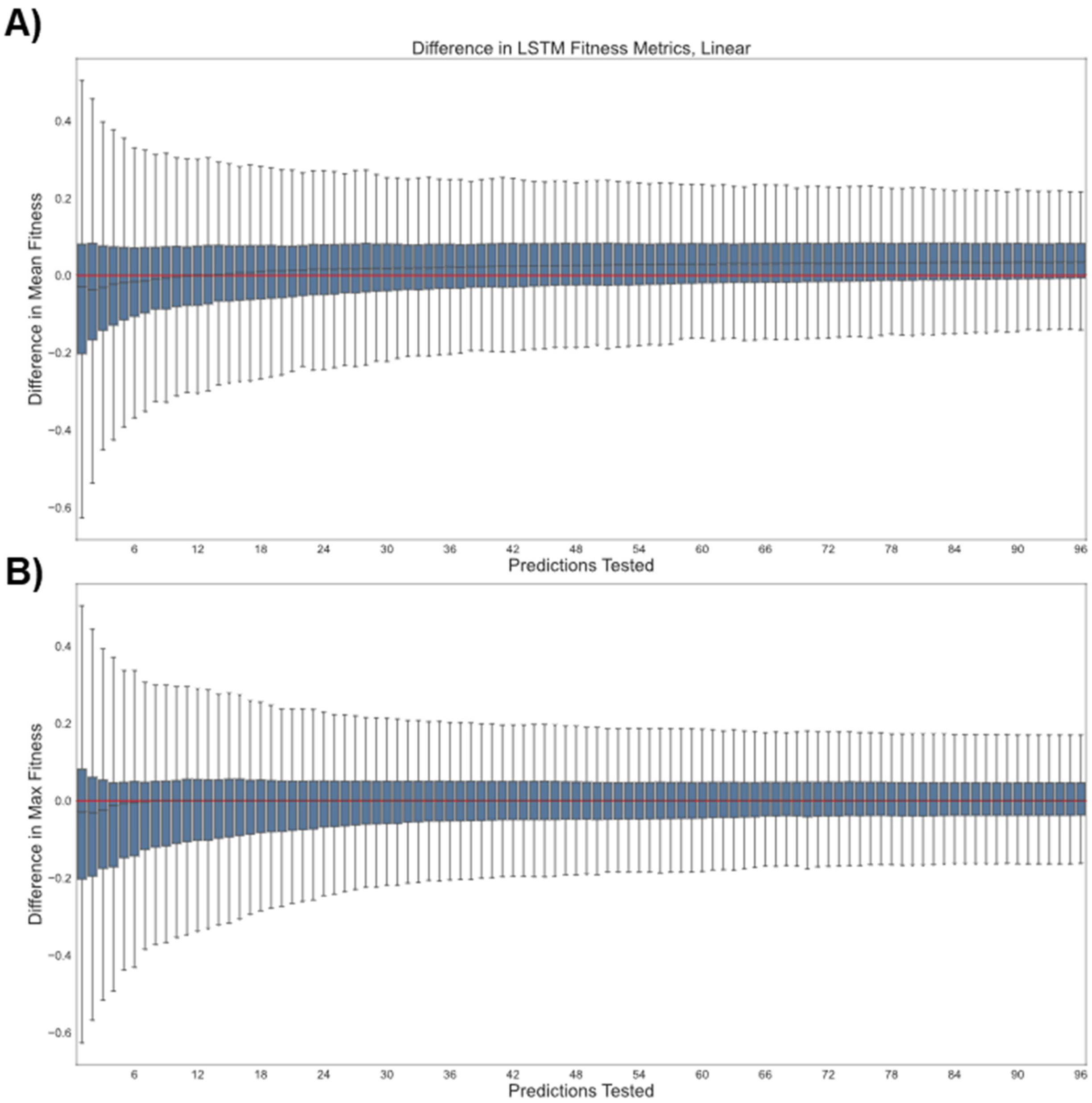

**Figure S34.** Pairwise comparison of fitness results for XGBoost with the Tweedie regression objective (reg:tweedie) and the default regression objective (reg:squarederror) over 2000 simulations using LSTM-derived encodings and a base linear model. The x-axis gives the number of top predictions tested (e.g. when x = 42, the 42 variants predicted to have the highest fitness by MLDE were evaluated for each regression strategy) and the y-axis gives the difference in either (A) the mean or (B) the maximum fitness achieved in that top sample. All differences are reported as reg:tweedie result – reg:squarederror result. The red line serves as a reference for 0 difference. Probability mass above the red line indicates a superior result using Tweedie regression. Extra details on the meaning and construction of this figure can be found in paragraphs at the top of this section.

#### The Challenge of Holes in Combinatorial Fitness Landscapes and the Importance of Informative Training Data

This section provides supporting information for the rounds of simulated focused training MLDE (ftMLDE) performed using simulated classifiers. Figure S35 provides summary statistics for the training sets used in these simulations. Each empirical cumulative distribution function (ECDF) in the below figure is generated from 2000 sets of 384 training variants; Figure S35A plots the mean fitness of each of these sets; Figure S35B plots the mean pairwise hamming distance between the members within each set. Table S7 provides summary statistics for rounds of simulated ftMLDE performed using these training sets.

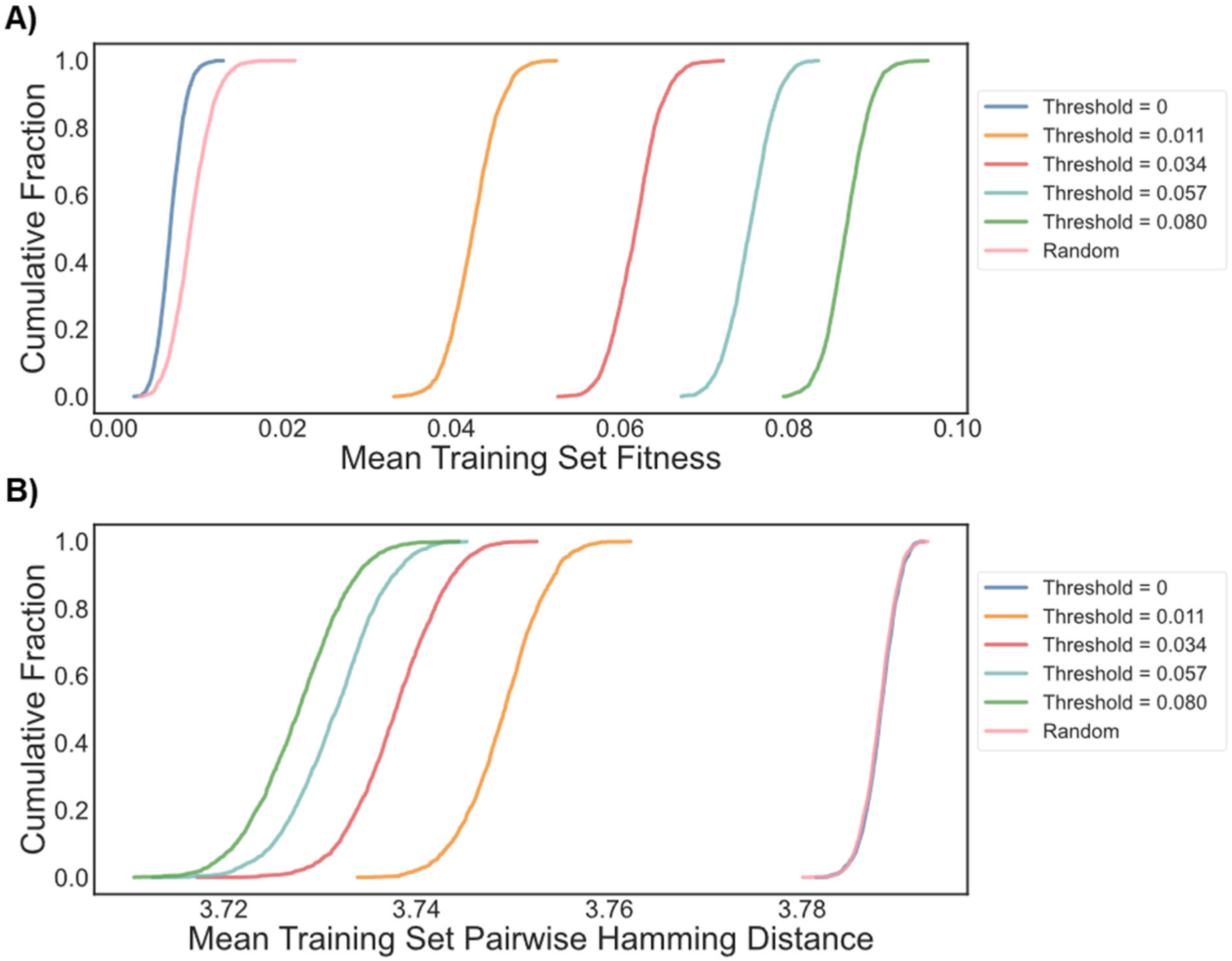

**Figure S35.** Summary statistics (shown as empirical cumulative distribution functions) for the 2000 training sets (each consisting of 384 samples) generated using the simulated weak classifiers. For a given threshold, 50% of variants in the training data have fitness greater than or equal to the threshold and the remainder have fitness less than or equal to the threshold. When the threshold = 0, 100% of variants in the training data have fitness greater than or equal to 0. For all classifiers (including the one with a threshold at 0), the maximum allowed fitness in the training data is 0.34. The random sample is equivalent to the classifier with a threshold at 0, but does not have this upper bound on training fitness. (A) Plots of the mean fitness of all variants in a training set for all 2000 training sets derived from each classifier. As the classification threshold increases, the mean fitness rises as expected. (B) Plots of the mean pairwise hamming distance between all members of a training set for all 2000 training sets derived from each classifier. A higher mean pairwise hamming distance indicates greater sequence diversity in the training data. As the classification threshold increases, the mean pairwise hamming distance decreases. This is because the training data is increasingly restricted to the narrow regions of sequence space that contain higher-fitness variants.

**Table S7.** Expected NDCG, mean of the top 96 predictions, and max of the top 96 predictions for the 2000 ftMLDE simulations performed using each of the simulated weak classifiers for training data generation.

| Classification Threshold | NDCG | Mean | Max |
|---|---|---|---|
| **0** | 0.819 | 0.209 | 0.654 |
| **0.011** | 0.884 | 0.313 | 0.777 |
| **0.034** | 0.894 | 0.329 | 0.797 |
| **0.057** | 0.899 | 0.340 | 0.816 |
| **0.08** | 0.901 | 0.341 | 0.815 |

#### Predicted ΔΔG of Stabilization for the Design of Fitness-Enriched Training Data

This section provides supplementary results pertaining to the predicted ΔΔG of GB1 protein stability (procedure detailed above in *ΔΔG Calculations*). Figure S36 compares literature values of ΔΔG for single-mutant variants of GB1 present in the fitness landscape used in this study against (A) the fitness of those variants, (B) the ΔΔG predicted from Triad calculations using a fixed backbone, and (C) the ΔΔG predicted from Triad calculations using a flexible backbone.[15] Figure S37 is an alternate representation of Figure 4A from the main text, showing a rolling mean with a linear-scale y-axis (as opposed to the rolling median on a log-scale axis used in the main text). Figure S38 shows the results of zero-shot predictions using fixed-backbone calculations when only considering quadruple mutants. The fact that high-enrichment is maintained suggests that Triad calculations are able to identify higher-fitness mutants away from the parent protein. Figure S39 shows the results of zero-shot predictions using flexible-backbone calculations—essentially no enrichment is observed, despite the ability of flexible-backbone calculations to predict experimental ΔΔG. Figure S40 shows the results of zero-shot predictions using flexible-backbone calculations when ranking GB1 variants by backbone root mean squared deviation (RMSD) rather than predicted ΔΔG. Enrichment is observed when ranking by RMSD, suggesting that structural conservation may be the factor that makes predicted ΔΔG an effective zero-shot strategy for GB1. Figure S41 is the crystal structure (PDB: 2GI9) used as the scaffold for ΔΔG calculations.[14]

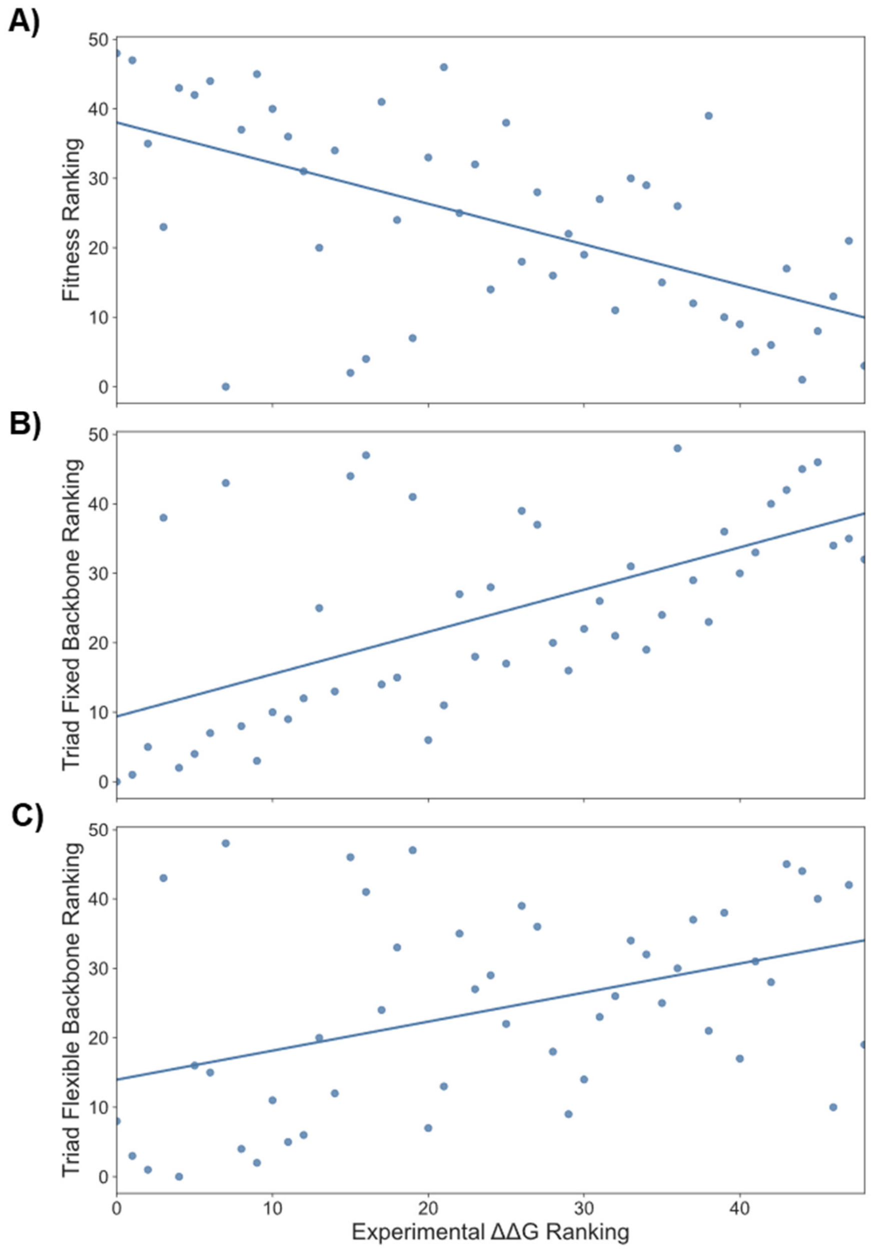

**Figure S36.** (A) Relationship between experimentally determined ΔΔG and GB1 fitness. The fitness of GB1 (at least at the considered positions (V39, D40, G41, and V54)) is loosely correlated with ΔΔG, but is clearly not the only determinant, with some lower-fitness variants having low ΔΔG and some higher-fitness variants having high ΔΔG. (B) Comparison of predicted ΔΔG upon mutation for GB1 variants using fixed backbone calculations to experimentally measured values of ΔΔG. (C) Comparison of predicted ΔΔG for GB1 variants using flexible backbone calculations to experimentally measured values of ΔΔG. Note that the experimentally reported ΔΔG values of -4 (those protein that did not express) were ignored in this analysis.

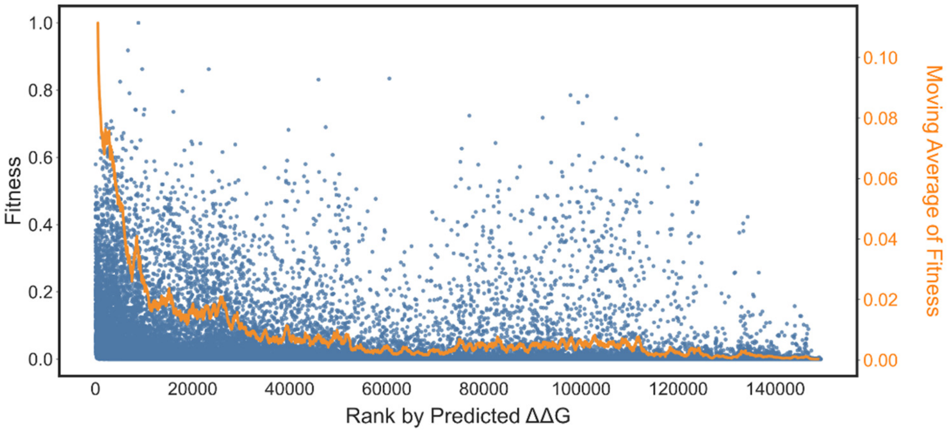

**Figure S37.** The fitness of all GB1 variants plotted against the rank (from lowest to highest ΔΔG) given by Triad calculations using fixed backbone calculations. Blue dots are all individual variants while the orange line is the sliding mean (window size = 1000) of fitness.

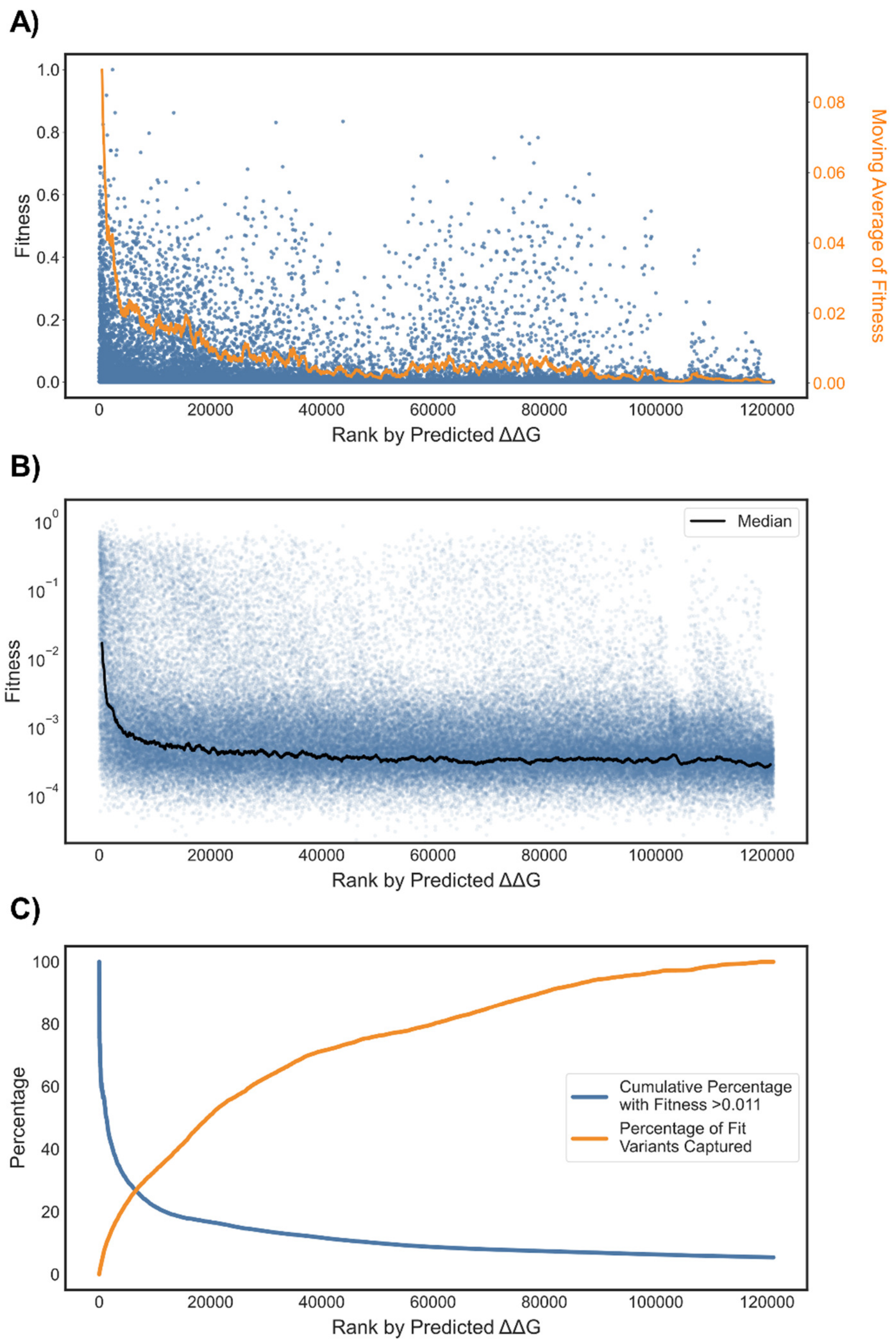

**Figure S38.** Results of zero-shot prediction using fixed backbone Triad ΔΔG calculations when only considering quadruple mutants relative to the parent GB1 protein (V39 D40 G41 V54) (|Spearman ρ| = 0.19). (A) The fitness of quadruple mutants plotted against the rank (from lowest to highest ΔΔG) given by Triad calculations. Blue dots are all individual variants while the orange line is the sliding mean (window size = 1000) of fitness. (B) The log-fitness of all quadruple mutants plotted against the rank given by Triad calculations. Blue dots are all individual variants while the black line is the sliding median (window size = 1000) of fitness. (C) Cumulative fitness metrics for quadruple mutants ranked by Triad score. The blue curve gives the percentage of variants ranked up to and including a given Triad rank that have fitness greater than 0.011 (the cutoff of the weakest simulated classifier in main text Figure 3). The orange curve gives the percentage of all "fit" (defined as fitness greater than 0.011) variants encompassed in the set up to and including a given Triad rank.

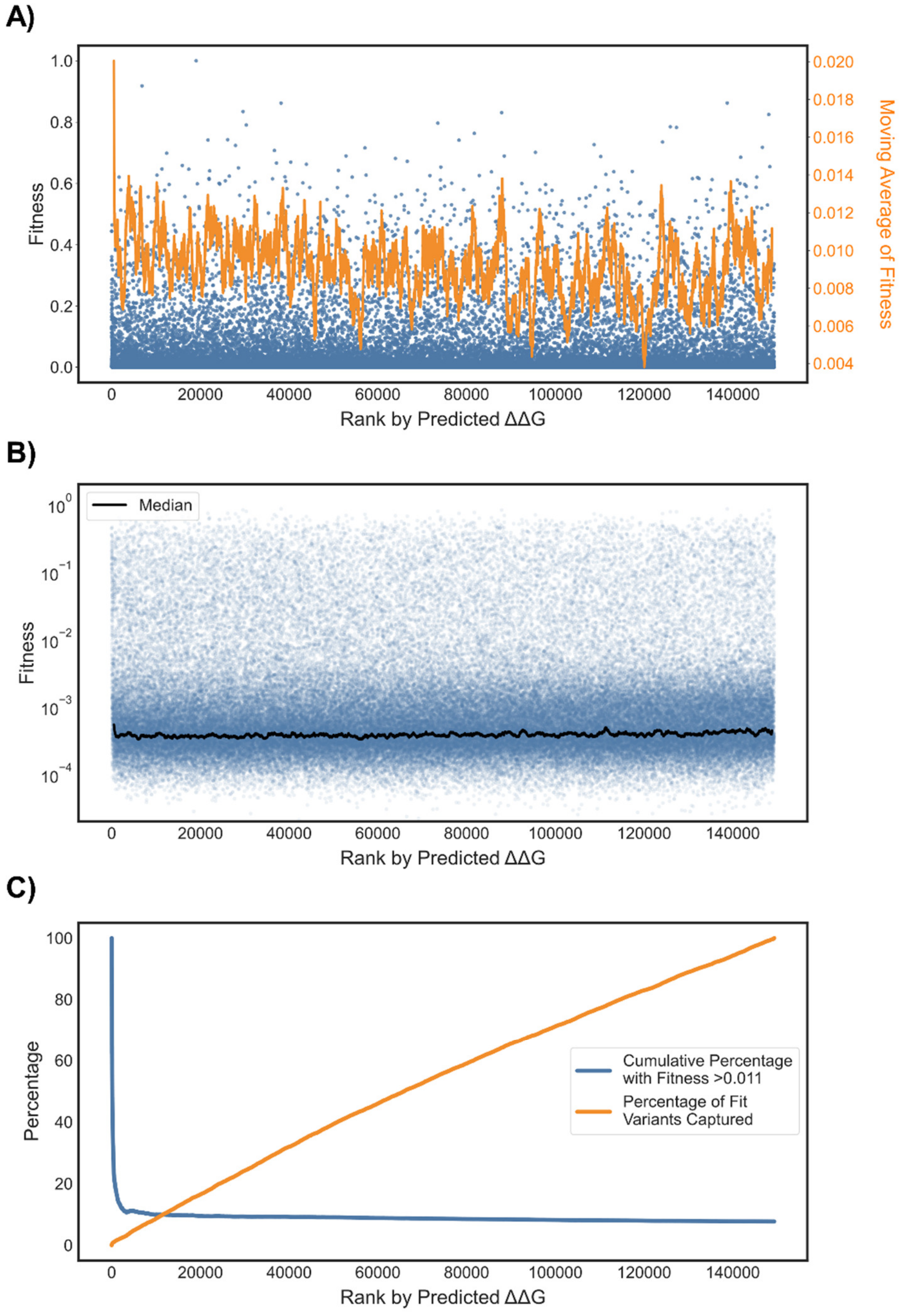

**Figure S39.** Results of zero-shot prediction using flexible backbone Triad ΔΔG calculations. (A) The fitness of all GB1 variants plotted against the rank (from lowest to highest ΔΔG) given by Triad calculations. Blue dots are all individual variants while the orange line is the sliding mean (window size = 1000) of fitness. (B) The log-fitness of all GB1 variants plotted against the rank given by Triad calculations. Blue dots are all individual variants while the black line is the sliding median (window size = 1000) of fitness. (C) Cumulative fitness metrics for all GB1 variants ranked by Triad score. The blue curve gives the percentage of variants ranked up to and including a given Triad rank that have fitness greater than 0.011 (the cutoff of the weakest simulated classifier in main text Figure 3). The orange curve gives the percentage of all "fit" (defined as fitness greater than 0.011) variants encompassed in the set up to and including a given Triad rank.

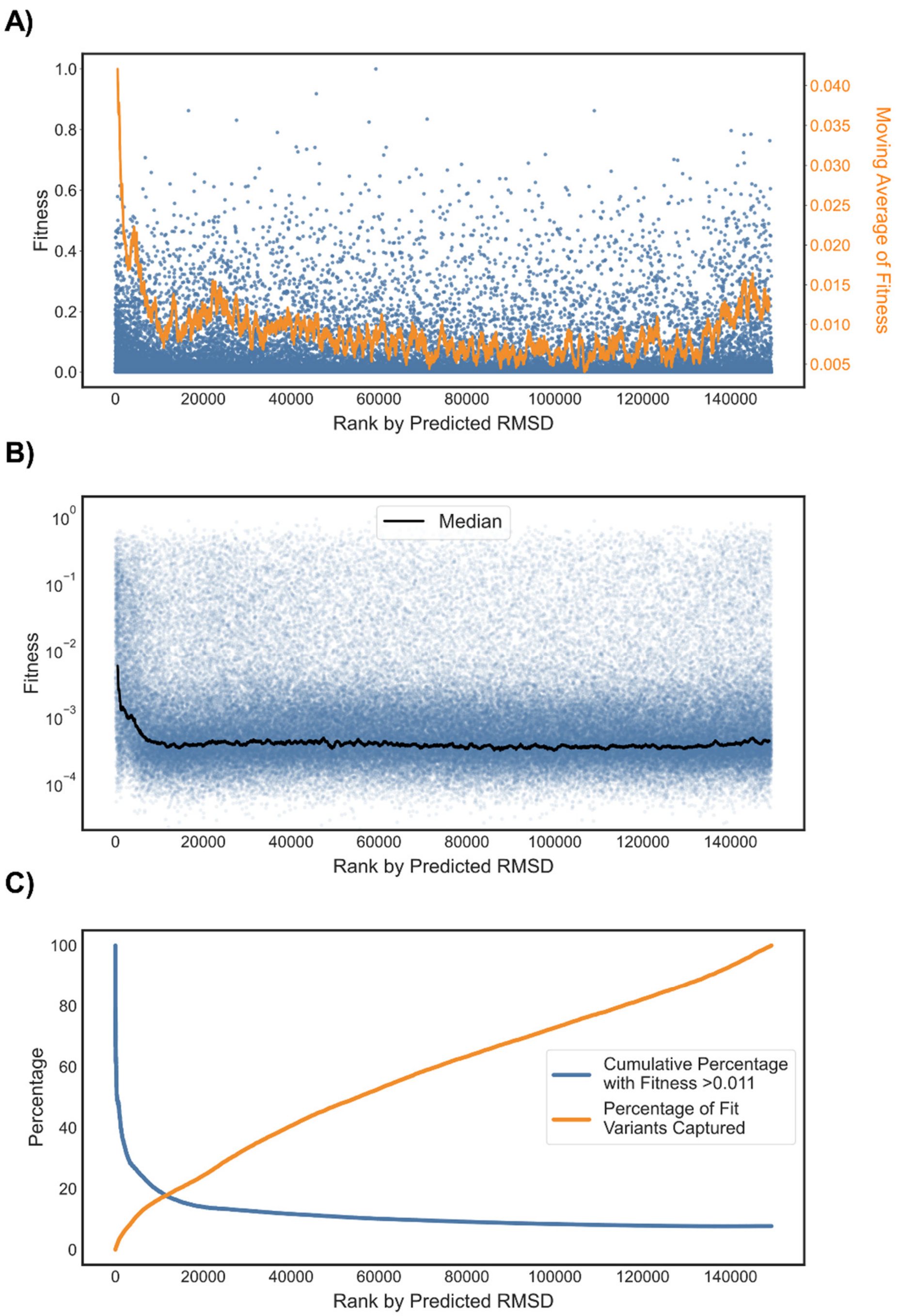

**Figure S40.** Results of zero-shot prediction using flexible backbone Triad root mean squared deviation (RMSD) calculations. (A) The fitness of all GB1 variants plotted against the rank (from lowest to highest RMSD) given by Triad calculations. Blue dots are all individual variants while the orange line is the sliding mean (window size = 1000) of fitness. (B) The log-fitness of all GB1 variants plotted against the rank given by Triad calculations. Blue dots are all individual variants while the black line is the sliding median (window size = 1000) of fitness. (C) Cumulative fitness metrics for all GB1 variants ranked by Triad score. The blue curve gives the percentage of variants ranked up to and including a given Triad rank that have fitness greater than 0.011 (the cutoff of the weakest simulated classifier in main text Figure 3). The orange curve gives the percentage of all "fit" (defined as fitness greater than 0.011) variants encompassed in the set up to and including a given Triad rank.

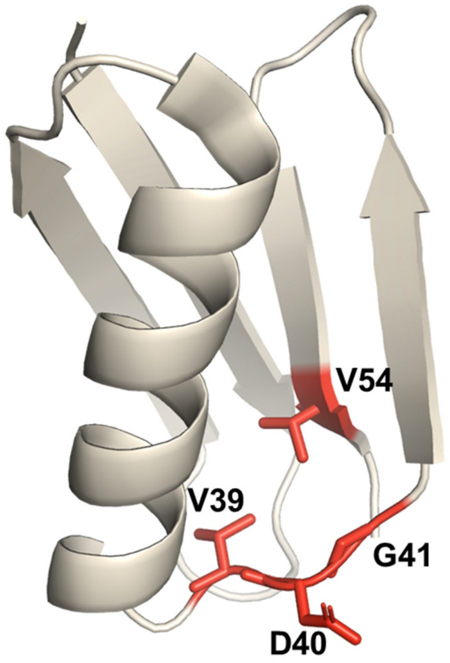

**Figure S41.** GB1 crystal structure (PDB: 2GI9) with the positions mutated in the GB1 combinatorial landscape highlighted in red.

#### Predicted ΔΔG of Stabilization for Training Set Design Enables Highly Effective ftMLDE on the GB1 Landscape

This section provides supplementary results pertaining to the ftMLDE simulations run using training data enriched in fitness using predicted ΔΔG as a zero-shot fitness-prediction strategy. Figure S42 provides summary statistics for the training sets containing 384 variants used in these simulations. Each empirical cumulative distribution function (ECDF) in the below figure is generated from 2000 sets of 384 training variants; Figure S42A plots the mean fitness of each of these sets; Figure S42B plots the mean pairwise hamming distance between the members within each set. Table S8 gives the summary statistics for all conditions tested during simulations using training data generated from Triad zero-shot predictions.

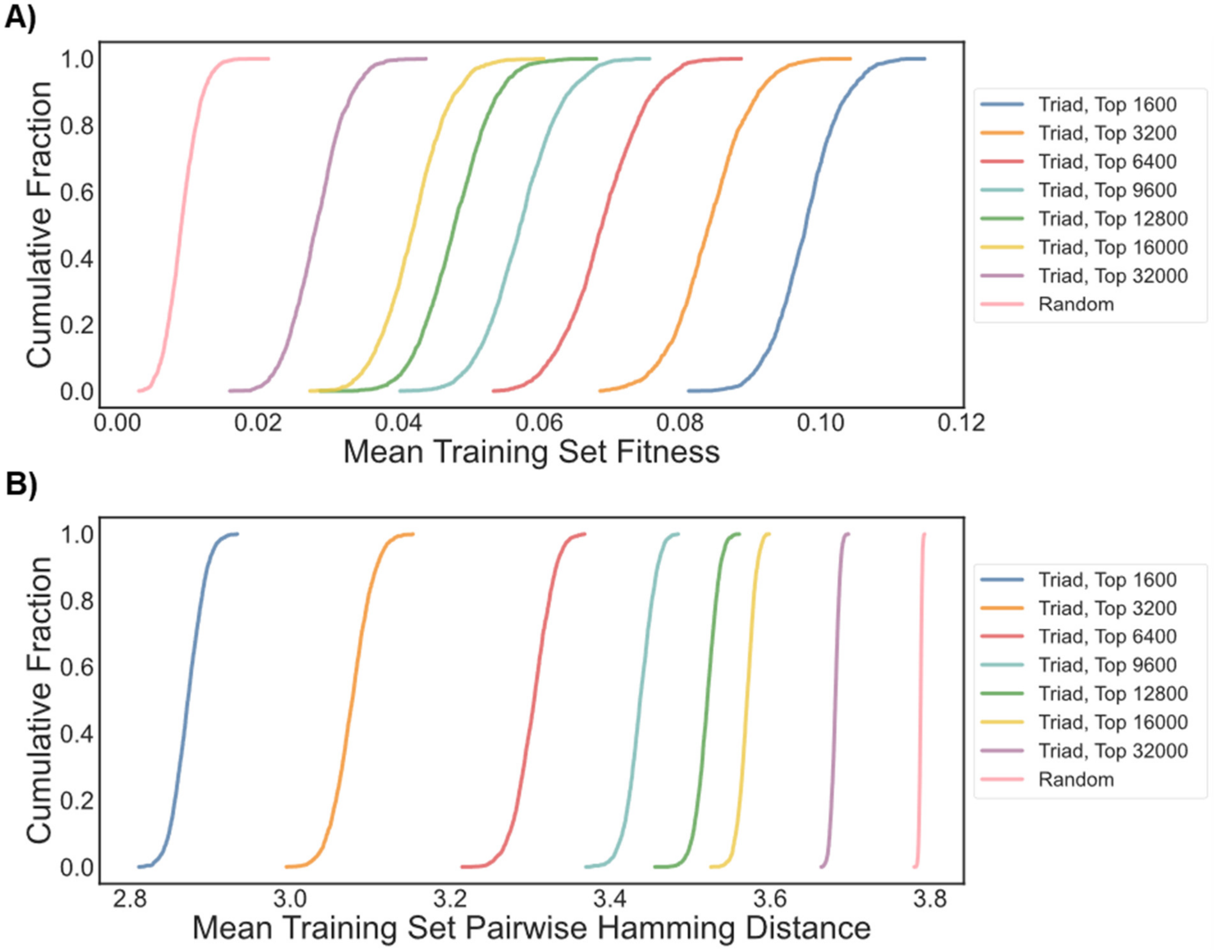

**Figure S42.** Summary statistics (shown as empirical cumulative distribution functions) for the 384-sample training sets generated using Triad zero-shot prediction. (A) Plots of the mean fitness of all variants in a training set for all 2000 training sets derived for each sampling threshold. As the threshold increases, the mean fitness decreases as more low-fitness variants have the potential to be included in the training data. (B) Plots of the mean pairwise hamming distance between all members of a training set for all 2000 training sets derived from each classifier. A higher mean pairwise hamming distance indicates greater sequence diversity in the training data. As the threshold increases, the mean pairwise hamming distance also increases. This is because predictive algorithms will tend to group similar sequences as having similar properties. For Triad, this means that sequences close in rank-order (by ΔΔG) will be similar. By increasing the range of rank sampled from, the range of sequences sampled from is thus also increased.

**Table S8.** Summary statistics for the ftMLDE simulations performed using Triad zero-shot predictions for training set design. Expected mean and max fitness values are reported for the tested predictions. "Frequency Unsampled Max Achieved" gives the frequency with which the maximum value in the unlabeled datapoints was identified over all simulated rounds of MLDE; note that the maximum value in the unlabeled datapoints is identical to the global maximum of the GB1 fitness landscape unless the global maximum is found in the training data. The column "Frequency Global Max Achieved" gives the frequency with which the highest-fitness variant in the GB1 dataset was identified in either the training variants or the top-predictions tested. For reference, a greedy walk achieves the global optimum 1.2% and has an expected max fitness achieved of 0.45.

| Training Points | Tested Predictions | Sampling Limit | Expected Max Fitness Achieved | Expected Mean Fitness Achieved | Frequency Unsampled Max Achieved | Frequency Global Max Achieved |
|---|---|---|---|---|---|---|
| 24 | 56 | 1600 | 0.599 | 0.198 | 4.95% | 4.95% |
| 24 | 56 | 3200 | 0.628 | 0.181 | 5.80% | 5.80% |
| 24 | 56 | 6400 | 0.627 | 0.174 | 5.55% | 5.55% |
| 24 | 56 | 9600 | 0.604 | 0.156 | 4.05% | 4.25% |
| 24 | 56 | 12800 | 0.587 | 0.141 | 3.60% | 3.70% |
| 24 | 56 | 16000 | 0.573 | 0.137 | 2.40% | 2.40% |
| 24 | 56 | 32000 | 0.524 | 0.119 | 1.30% | 1.30% |
| 24 | 56 | Random | 0.418 | 0.073 | 0.40% | 0.50% |
| 48 | 32 | 1600 | 0.618 | 0.263 | 5.75% | 5.75% |
| 48 | 32 | 3200 | 0.672 | 0.240 | 9.60% | 9.60% |
| 48 | 32 | 6400 | 0.674 | 0.249 | 7.30% | 7.30% |
| 48 | 32 | 9600 | 0.650 | 0.228 | 6.40% | 6.65% |
| 48 | 32 | 12800 | 0.632 | 0.217 | 5.30% | 5.45% |
| 48 | 32 | 16000 | 0.614 | 0.205 | 4.15% | 4.40% |
| 48 | 32 | 32000 | 0.560 | 0.178 | 2.10% | 2.10% |
| 48 | 32 | Random | 0.476 | 0.130 | 0.50% | 0.60% |
| 384 | 96 | 1600 | 0.723 | 0.347 | 14.85% | 14.85% |
| 384 | 96 | 3200 | 0.985 | 0.389 | 91.80% | 91.80% |
| 384 | 96 | 6400 | 0.976 | 0.445 | 82.05% | 82.05% |
| 384 | 96 | 9600 | 0.970 | 0.426 | 76.95% | 77.20% |
| 384 | 96 | 12800 | 0.948 | 0.380 | 66.30% | 66.55% |
| 384 | 96 | 16000 | 0.898 | 0.336 | 44.65% | 44.95% |
| 384 | 96 | 32000 | 0.851 | 0.322 | 29.35% | 29.90% |
| 384 | 96 | Random | 0.751 | 0.267 | 8.65% | 8.75% |

#### Supplementary References

(1) Georgiev, A. G. Interpretable Numerical Descriptors of Amino Acid Space. *J. Comput. Biol.* **2009**, *16*, 703–723. https://doi.org/10.1089/cmb.2008.0173.

(2) Ofer, D.; Linial, M. ProFET: Feature Engineering Captures High-Level Protein Functions. *Bioinformatics* **2015**, *31*, 3429–3436. https://doi.org/10.1093/bioinformatics/btv345.

(3) Rao, R.; Bhattacharya, N.; Thomas, N.; Duan, Y.; Chen, X.; Canny, J.; Abbeel, P.; Song, Y. S. Evaluating

Protein Transfer Learning with TAPE. *arXiv* **2019**. arXiv:1906.08230.
(4) Bepler, T.; Berger, B. Learning Protein Sequence Embeddings Using Information from Structure. *arXiv* **2019**. arXiv:1902.08661.
(5) Alley, E. C.; Khimulya, G.; Biswas, S.; AlQuraishi, M.; Church, G. M. Unified Rational Protein Engineering with Sequence-Based Deep Representation Learning. *Nat. Methods* **2019**, *16*, 1315–1322. https://doi.org/10.1038/s41592-019-0598-1.
(6) Bergstra, J.; Yamins, D.; Cox, D. D. Making a Science of Model Search: Hyperparameter Optimization in Hundreds of Dimensions for Vision Architectures. *Proc. 30th Int. Conf. Mach. Learn.* **2013**.
(7) Chen, T.; Guestrin, C. XGBoost: A Scalable Tree Boosting System. *arXiv* **2016**. arXiv:1603.02754.
(8) Wu, Z.; Kan, S. B. J.; Lewis, R. D.; Wittmann, B. J.; Arnold, F. H. Machine Learning-Assisted Directed Protein Evolution with Combinatorial Libraries. *Proc. Natl. Acad. Sci.* **2019**, *116*, 8852–8858. https://doi.org/10.1073/pnas.1901979116.
(9) Buitinck, L.; Louppe, G.; Blondel, M.; Pedregosa, F.; Mueller, A.; Grisel, O.; Niculae, V.; Prettenhofer, P.; Gramfort, A.; Grobler, J.; Layton, R.; Vanderplas, J.; Joly, A.; Holt, B.; Varoquaux, G. API Design for Machine Learning Software: Experiences from the Scikit-Learn Project. *arXiv* **2013**. arXiv:1309.0238.
(10) Hopf, T. A.; Green, A. G.; Schubert, B.; Mersmann, S.; Schärfe, C. P. I.; Ingraham, J. B.; Toth-Petroczy, A.; Brock, K.; Riesselman, A. J.; Palmedo, P.; Kang, C.; Sheridan, R.; Draizen, E. J.; Dallago, C.; Sander, C.; Marks, D. S. The EVcouplings Python Framework for Coevolutionary Sequence Analysis. *Bioinformatics* **2019**, *35*, 1582–1584. https://doi.org/10.1093/bioinformatics/bty862.
(11) Hopf, T. A.; Ingraham, J. B.; Poelwijk, F. J.; Schärfe, C. P. I.; Springer, M.; Sander, C.; Marks, D. S. Mutation Effects Predicted from Sequence Co-Variation. *Nat. Biotechnol.* **2017**, *35*, 128–135. https://doi.org/10.1038/nbt.3769.
(12) Riesselman, A. J.; Ingraham, J. B.; Marks, D. S. Deep Generative Models of Genetic Variation Capture the Effects of Mutations. *Nat. Methods* **2018**, *15*, 816–822. https://doi.org/10.1038/s41592-018-0138-4.
(13) Biswas, S.; Khimulya, G.; Alley, E. C.; Esvelt, K. M.; Church, G. M. Low-N Protein Engineering with Data-Efficient Deep Learning. *bioRxiv* **2020**. https://doi.org/10.1101/2020.01.23.917682.
(14) Franks, W. T.; Wylie, B. J.; Stellfox, S. A.; Rienstra, C. M. Backbone Conformational Constraints in a Microcrystalline U-$^{15}$N-Labeled Protein by 3D Dipolar-Shift Solid-State NMR Spectroscopy. *J. Am. Chem. Soc.* **2006**, *128*, 3154–3155. https://doi.org/10.1021/ja058292x.
(15) Nisthal, A.; Wang, C. Y.; Ary, M. L.; Mayo, S. L. Protein Stability Engineering Insights Revealed by Domain-Wide Comprehensive Mutagenesis. *Proc. Natl. Acad. Sci.* **2019**, *116*, 16367–16377. https://doi.org/10.1073/pnas.1903888116.
